# A single cell atlas of the healthy breast tissues reveal clinically relevant clusters of breast epithelial cells

**DOI:** 10.1101/2020.06.25.171793

**Authors:** Poornima Bhat-Nakshatri, Hongyu Gao, Patrick C. McGuire, Xiaoling Xuei, Liu Sheng, Jun Wan, Yunlong Liu, Sandra K. Althouse, Austyn Colter, George Sandusky, Anna Maria Storniolo, Harikrishna Nakshatri

## Abstract

Single cell RNA sequencing is an evolving field to elucidate cellular architecture of adult organs. Using normal breast tissues from healthy volunteers and a rapid procurement/processing/sequencing protocol, 13 breast epithelial cell clusters were identified. Approximately 90% of breast cancers were enriched for cell-of-origin signatures derived from differentiated luminal clusters and two minor luminal progenitor clusters. Expression of cell cycle and chromosome segregation-related genes were higher in one of the minor clusters and breast tumors with this cluster signature displayed the highest mutation rate and poor outcome. We identified TBX3 and PDK4 as genes co-expressed with estrogen receptor (ER) in the normal breasts and their expression analyses in >550 breast cancers enabled prognostically relevant cell-of-origin based subclassification of ER+ breast cancers.

**Significance:** This study elucidates different epithelial cell types of the normal breasts and identifies a minor subpopulation of cells from which the majority of breast cancers may originate. This observation should help to develop methods to characterize breast tumors based on cell-of-origin. Although it was suggested that intrinsic subtypes of breast cancers have distinct cells of origins, this study suggests multiple cell-of-origin for an intrinsic subtype of breast cancer, including for hormone responsive breast cancers. Cell-of-origin signatures allowed survival-associated subclassification of intrinsic subtypes. Critically, this normal breast cell atlas would allow for the classification of genes differentially expressed in a breast tumor compared to normal breast due to the cell-of-origin of tumor and those that are acquired due to genomic aberrations.

## Introduction

Breast cancers are subclassified into multiple subtypes based on gene expression analyses and genomic aberrations (Curtis et al., 2012; Sotiriou et al., 2003). These classifications have played a significant role in clinical decision making. Among these classifications, intrinsic subtype classification based on gene expression, which classifies breast cancer into luminal-A, luminal-B, HER2+, basal, normal-like and claudin-low, is suggested to reflect cell-of-origin of breast cancer (Prat and Perou, 2009). Flow cytometry-based marker profiling and gene expression portraits have identified three major epithelial cell types in the breast including basal/stem (CD49f+/EpCAM-), luminal progenitors (CD49f+/EpCAM+), and mature luminal (CD49f-/EpCAM+) cells (Lim et al., 2009; Visvader and Stingl, 2014). Cell-type enriched transcription factor networks such as *TP63/NFIB, ELF5/EHF* and *FOXA1/ESR1* control gene expression patterns in basal/stem, luminal progenitor and mature luminal cells, respectively (Pellacani, 2016). It is suggested that while claudin-low subtype of breast cancers originates from basal/stem cells, luminal progenitors are the source of basal-like breast cancers (Lim et al., 2009; Prat et al., 2010; Proia et al., 2011). While HER2+ breast cancers may originate from luminal progenitors and mature luminal cells, luminal-A/B breast cancers likely originate from mature luminal cells (Prat and Perou, 2009). However, it is acknowledged that heterogeneity exists within basal/stem, luminal progenitors and mature luminal cells as defined by CD49f/EpCAM cell surface marker profiling (Anjanappa et al., 2017; Colacino et al., 2018).

A recent integrative analysis of 10,000 tumors from 33 types of cancer emphasized the dominant role of cell-of-origin patterns in cancers (Hoadley et al., 2018). Since normal tissue itself is composed of multiple cell types, fine mapping of these cell types and identifying potential cancer vulnerable cell population in normal tissues would aid in identifying and characterizing organ-specific cell-of-origin of cancers. Recent advances in single cell techniques including scRNA-seq, sc-Epigenetics, scDNA-seq and scProteomics-atlas are enabling further refinement of cell types within normal and diseased tissues (Lim et al., 2020). For example, using reduction mammoplasty samples and cells flow sorted based on CD49f/EpCAM, Nguyen et al identified three epithelial cell types in the normal breasts (Nguyen et al., 2018). Using the same technique and mouse mammary tissues at different development stages, Pal et al described seven epithelial cell types in the mouse mammary gland (Pal et al., 2017). However, Bach et al observed 15 epithelial cell types in the mouse mammary gland (Bach et al., 2017). A concern has been raised about the reproducibility of data, which is likely influenced by the types of tissues used, duration between tissue collection and sequencing, and dissociation protocol (Lim et al., 2020). While these issues can be standardized for studies involving mouse tissues, standardizing is difficult for studies that utilize human tissues collected after a surgical procedure. In this regard, in an elegant review, Lim et al recently proposed the need to establish a rapid tissue dissociation program to advance single cell technology for clinical applications (Lim et al., 2020).

A decade ago, our institution established a normal breast tissue bank where clinically healthy women donate breast biopsies for research purposes. This resource has enabled others and us to demonstrate clear differences between our “normal” and both reduction mammoplasty and tumor-adjacent normal tissues, which have been the most common sources of “normal controls” for breast cancer studies in the literature, including the single cell transcriptome study detailed above. We and others have been able to show clear histologic and molecular abnormalities in these surrogate sources of normal breast tissue (Degnim et al., 2012; Nakshatri et al., 2015; Nakshatri et al., 2019). In this study, we first performed single cell RNA sequencing of five freshly collected samples that included 18704 cells and 20647 genes. Results were analyzed at both single sample levels as well as in an integrative manner. To confirm the results of first sequencing, we repeated integrated single cell analyses of five new cryopreserved samples covering 7582 cells and 25,842 genes. Using the expression patterns of CD49f and EpCAM as well as basal/stem, luminal progenitor and mature luminal cell transcription factor networks, we performed refined analyses of epithelial cells. Epithelial cluster specific gene signatures were then applied on TCGA and METABRIC datasets to determine the impact of putative “cell-of-origin” on breast cancer outcomes (Cancer Genome Atlas, 2012; Curtis et al., 2012). Since there is limited subclassification of estrogen receptor positive (ER+) breast cancers and it is exceeding difficult to characterize ER+ breast epithelial cells from the normal breasts to identify genes co-expressed with ER in normal and tumor cells (Rosenbluth et al., 2020), we performed additional studies on *TBX3* and *PDK4*, two genes that are co-expressed at different levels in ER+ clusters of the normal breasts.

## Results

### Establishment of rapid tissue procurement and single cell analyses protocol

Although the primary intention of establishing the Susan G. Komen Tissue Bank at IU Simon Cancer Center (Komen Tissue Bank) was to provide a source of healthy breast tissue to be used as normal controls for research, we took advantage of tissue collection procedure easily accessible in clinic instead of surgical room to limit time lapse between tissue collection and utilization of tissues for research purposes typically associated with tissue collection after surgical procedures. Because these “collection events” have 1:2 donor: volunteer ratio, we were abundantly staffed and able to reduce of time of tissue collection to placement in media or cold ischemia time for cryopreservation to ∼6 minutes. Since specimen cellularity varied between individuals, 50% of fresh or cryopreserved tissues provided high quality data with respect to number of viable cells and minimal ambient RNA contamination of single cell data. Also note that all tissues used have undergone histologic characterization and are free of abnormalities. Table S1 provides information about donors. Nine samples were from Caucasian women, one was from Asian, and one was from African American woman. Genetic ancestry mapping has been performed using 41-SNP genetic ancestry informative markers. Two out of 11 were nulliparous and two out of 11 were post-menopausal women. Five donors had a family history of breast cancer.

### Normal breast contains 13 epithelial cell types

Unlike the previous studies which purified breast epithelial cells using CD49f/EpCAM markers prior to single cell analyses (Nguyen et al., 2018), we subjected single cells after dissociation directly to RNA-sequencing and then used CD49f/EpCAM as well as transcriptional regulators known to specify basal/stem, luminal progenitors and mature luminal cells to subcluster epithelial cells (Pellacani, 2016). Uniform manifold approximation and projection (UMAP) plot of combined samples is shown in Figure 1A. As expected, the normal breasts contained a variety of cell types in addition to epithelial cells including monocytes, T cells, NK cells, endothelial cells, and fibroblast-like cells (Figure 1A). Fibroblast-like and endothelial cells displayed three closely related clusters suggesting heterogeneity within these cells. Heterogeneity in endothelial cells, driven largely by metabolic plasticity, has been previously described in other organ and disease conditions (Rohlenova et al., 2020). Similarly, heterogeneity in fibroblasts of mouse mammary gland with mammary tumors have also been described. (Bartoschek et al., 2018). Our studies show the heterogeneity in these cells within the normal breast itself. Epithelial cell types were dominant. Using CD49f/EpCAM expression pattern as well as *TP63/NFIB*, *ELF5/EHF* and *FOXA1/ESR1* as functional markers of basal/stem, luminal progenitors and mature luminal cells, we performed subcluster analyses of epithelial cells, whivch revealed 13 different epithelial cells (Figure 1B and C). Number of cells in each cluster and average expression value of genes that differentiated these clusters are shown in Table S2. A heatmap of average expression levels of top marker genes of these clusters is shown in Figure 1D. CD49f+/EpCAM- basal/stem cells contained three closely related subclusters (clusters 5, 7 and 9). Each of these clusters within basal/stem cells can be distinguished through expression of specific genes. For example, cluster 5 expressed higher levels of *NDUFA4L2*, a mitochondrial NADPH dehydrogenase (Figure 2A). This cluster also expressed higher levels of *CD36*, a lipid transporter associated with breast cancer metastasis, as well as *Vimentin*, a marker of basal cells (Pal et al., 2017; Pascual et al., 2017). Cluster 7 expressed *ACKR1*, a decoy receptor for CCL2 and IL-8 (Davis et al., 2015). Cluster 9 is enriched for *MECOM* (*EVI1*), a stem cell associated transcription factor (Sato et al., 2014).

**Figure 1:**
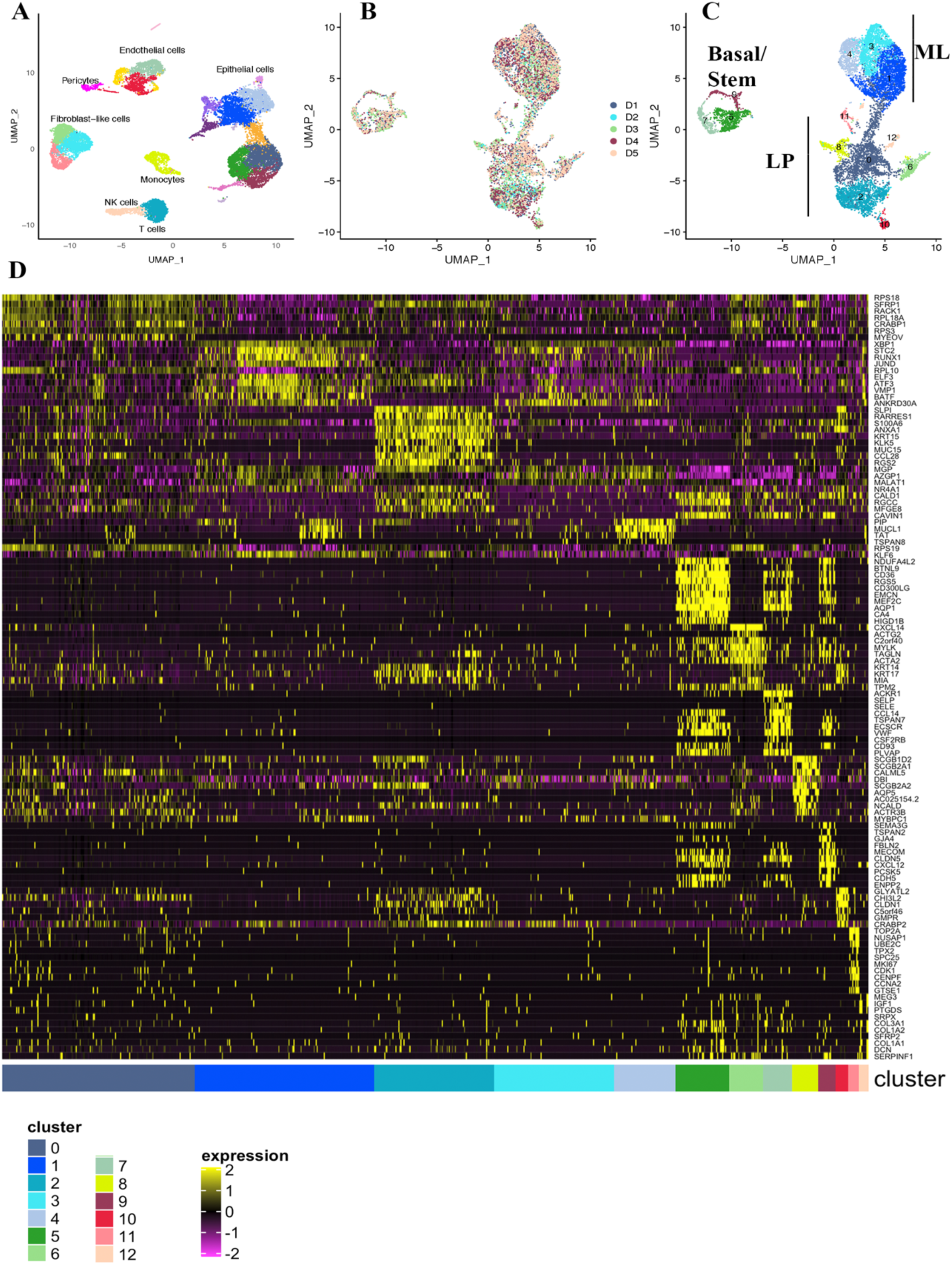
The normal breast contains 13 epithelial clusters. A) Integrated analysis of single cells of the normal breast biopsies of five healthy donors. Epithelial cells dominate among cell types. B) Subclustering of epithelial cell types using *CD49f/EpCAM* as well as *NFIB, TP63, EHF, ELF5, ESR1* and *FOXA1* expression patterns. D1-D5 corresponds to numbering of samples. C) Representation of various cell types in each sample. Subclusters in individual sample are shown in Figure S1A. D). Hierarchical clustering of top cluster-enriched genes.

**Figure 2:**
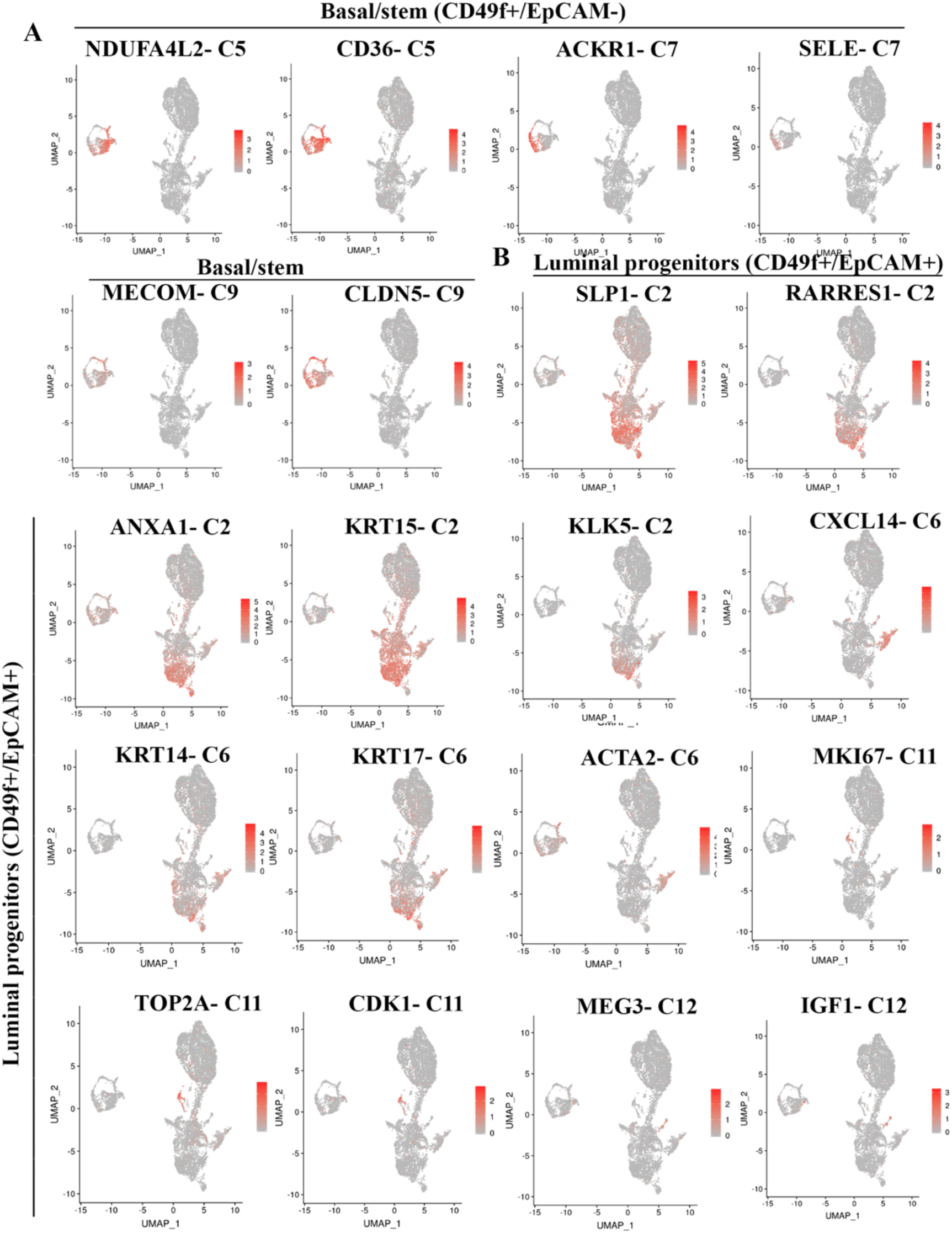
Expression patterns of representative cluster-enriched genes. A) Genes enriched in basal/stem cell clusters. B) Genes enriched in various clusters within luminal progenitor cells.

CD49f+/EpCAM+ cells contained two clusters that appeared as continuum of cells (clusters 0 and 2) and five other well-separated clusters (Clusters 6, 8, 10-12). Although cluster 0 and 2 appeared as a continuum of cells, significant differences in gene expression are evident (Figure 1D and 2B). For example, while all luminal progenitor cells expressed *Secreted Fizzled Related Protein 1* (*SFRP1*), a modulator of Wnt signaling (Baharudin et al., 2020), genes such as *SLP1, ANXA1, RARRES1, KLK5,* and *KRT15* were enriched in cluster 2. *KRT14* and *KRT17* expression were enriched in cluster 2 but not in cluster 0. Cluster 6 was enriched for *CXCL14* and *ACTA2*. Cluster 8 was enriched for *SCGB2A1* and *CALML5*. Cluster 10 was *KRT14*-positive and enriched for the expression of *GLYATL2*. Cluster 11 was enriched for the expression of multiple genes including *TOP2A, NUSAP1, UBE2C, TPX2, SPC25, MKI67, CDK1, CENPF* and *CCNA2* (Figure 1D and 2B). In fact, this cluster displayed a higher number of genes that are differentially expressed than other clusters and constituted a major signaling network associated with regulation of cell cycle, chromosome segregation, and spindle checkpoint to name few (Table S2). Cluster 12 was characterized by elevated expression of *MEG3, IGF1* and *PTGDS*.

CD49f-/EpCAM+ mature luminal cells were comprised of three clusters, which appeared as a continuum of cells (clusters 1, 3 and 4), although there were distinct differences in gene expression. All three of these clusters expressed *ESR1* and pioneering factors *FOXA1* and *GATA3* (Zaret and Carroll, 2011). *TBX3* and *PDK4* are two other genes that showed variable expression in these clusters. *XBP1* and *STC2*, *ESR*1 target genes (McBryan et al., 2007), were uniformly expressed at higher in all three of these clusters (Figure 3A). In fact, while expression level of *SFRP1* was able to distinguish luminal progenitors from mature luminal cells, expression levels of *XBP*1 and *STC2* were able to distinguish mature luminal cells from luminal progenitors. Cluster 1 showed enrichment of *RUNX1* and *BATF*. Cluster 3 was enriched for *ANKRD30A*, whereas Cluster 4 was enriched for *PIP, MUCL1, TAT* and *TSPAN8*.

**Figure 3:**
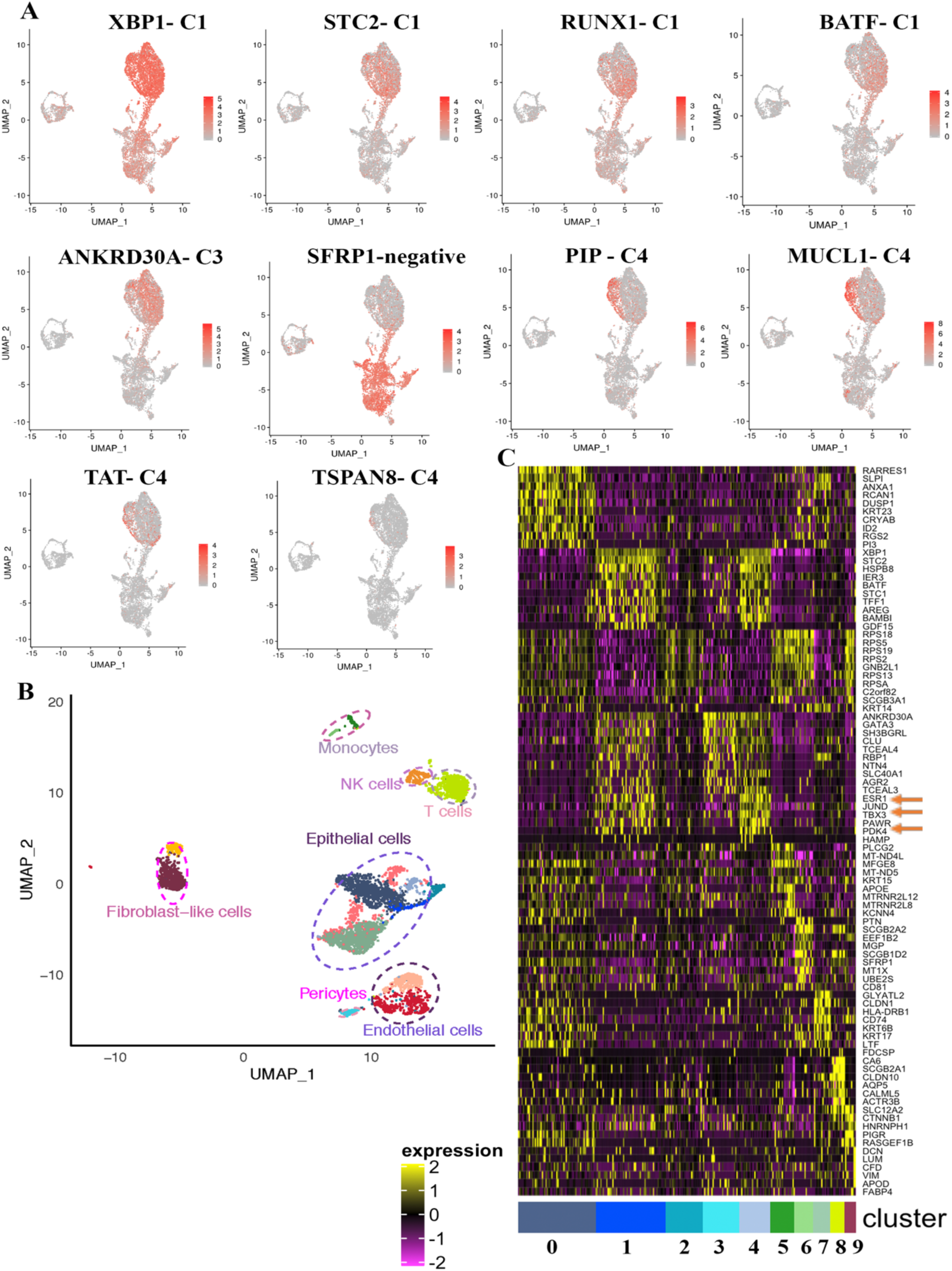
Mature luminal cells are enriched for *ESR1* and *XBP1*, whereas *SFRP1* is enriched in luminal progenitor cells. A) Genes enriched in mature luminal cells. Note that cluster 4 within mature luminal cells is distinctly enriched for *MUCL1* and *PIP*. B) Various cell types in the normal breast of a donor. C) Identification of *ESR1* expressing subclusters and genes co-expressed with *ESR1* in the normal breasts.

To highlight the presence of three clusters of cells expressing *ESR1* and to identify genes co-expressed with *ESR1*, we performed hierarchical clustering of breast epithelial cells from a 44-year old African American donor (donor 3). Various cell types present in the breast of this donor are shown in Figure 3B, and gene expression patterns in 10 epithelial cell clusters from this donor are shown in Figure 3C. Clusters 1, 3 and 4 were *ESR1+*. *JUND, TBX3, PAWR, PDK4*, and *HAMP* were the genes that distinguished cluster 4 from cluster 3, whereas cluster 1 contained lower levels of *GATA3* compared to clusters 3 and 4. Thus, there are at least three clusters of estradiol-responsive breast epithelial cells with varying levels of *ESR1* as well as pioneer factor expression.

The distribution pattern of epithelial clusters in the individual samples is shown in Figure S1A and number of cells per cluster is indicated in Table S2. Almost every sample contained similar levels of most of the clusters; two minor clusters, clusters 11 and 12, showed inter-sample variability.

### Reproducibility of cluster analyses

To determine whether epithelial clusters identified in the above analyses can be reproduced using cryopreserved tissues from healthy donors, we isolated cells from five cryopreserved tissues, pooled cells, and analyzed them all together. Since five samples were combined, there were enough cells to divide samples into two and perform cDNA synthesis and library preparation by two independent labs. In addition, we used the latest version of the library preparation with improved chemistry, paired-end sequencing and better efficiency. Because pooled samples contained more lymphocytes, lymphocyte related cells were removed from the analyses. Without lymphocyte removal, there were 28 clusters (Table S2). Side-by-side comparisons of the second set of pooled samples and re-analyses of first five samples are shown in Figure 4A. Re-analyses of individual samples are shown in Figure S1B and number of cells in each cluster is shown in Table S2. With increased number of cells and more genes sequenced, further subclassification of epithelial cells became possible. Since the number of clusters identified in this new clustering are different (23) from the first analysis (13), new clusters are named with prefix N (N0-N22). Consistent with our earlier report and a recent report on organoid-derived single cell data, there was inter-individual differences in proportion of cells in each cluster (Nakshatri et al., 2015; Rosenbluth et al., 2020). The UMAP cell embeddings and cell cluster information generated from Seurat analysis were imported into 10X genomics Loupe Browser. By checking various gene expression with the Loupe Browser, we first assigned the subdivided clusters into basal/stem, luminal progenitor, and luminal mature cells. Based on *CD49f* and *EpCAM* expression patterns (Figure 4B), clusters N5-7, N11, N13, N18 and N22 were basal; N3, N4, N9, N14, N16, N19 and N20 were luminal progenitors; and N2, N8, N12 and N17 were mature luminal cells. *ALDH1A3* expression, which has been suggested to identify breast cancer stem cells (Marcato et al., 2011), was expressed mainly in N4, N14 and N16 clusters of luminal progenitor cells (Figure 4C). Among transcription factors that define basal/stem, luminal progenitor and luminal mature cells, as expected, *ELF5* and *EHF* expression was restricted to luminal progenitor cells, whereas *ESR1* and *FOXA1* expression was restricted to luminal mature cell clusters (Figure S2). While *ESR1* expression was wide spread across mature luminal subclusters with cluster 12 displaying stronger signals, the expression of its target gene *PGR* was much more restricted within mature luminal cells suggesting the natural existence of ER+/PR+ and ER+/PR- cells, similar to the features of luminal A and luminal B breast cancers (Sorlie et al., 2001). Expression of best studied ER target gene *GREB1* overlapped with *PGR* expression suggesting ER has cell type-specific targets within the normal breast. Consistent with earlier report (Nolan et al., 2017), *RANK (TNFRSF11A)* expression was restricted to few luminal progenitor cells, whereas its ligand *RANKL (TNFSF11)* expression was observed in progesterone receptor positive mature luminal cells (Figure S2). *EHF* and *ELF5* expression showed strong signals in subclusters N14 and N16, which could be alveolar progenitor cells as these two transcription factors play a major role in alveolar differentiation during pregnancy (Luk et al., 2018). However, expression of *NFIB* and *TP63* did not correlate with prior assessment (Pellacani, 2016) as *NFIB* expression was not restricted to basal/stem cells, whereas *TP63* expression was observed only in cluster 15. Cluster 15 is likely myoepithelial cells, as this cluster expressed higher levels of *ACTA2* and *KRT17*, previously described markers of myoepithelial cells (Nguyen et al., 2018) (Figure S2). Although basal cells are expected to express *KRT14*, we found its expression predominantly in a subpopulation of luminal progenitor cells and in N15 with myoepithelial characteristics (Figure 4D). *KRT18* and *KRT19* expression were found equally in luminal progenitor and mature luminal subclusters (Figure S2).

**Figure 4:**
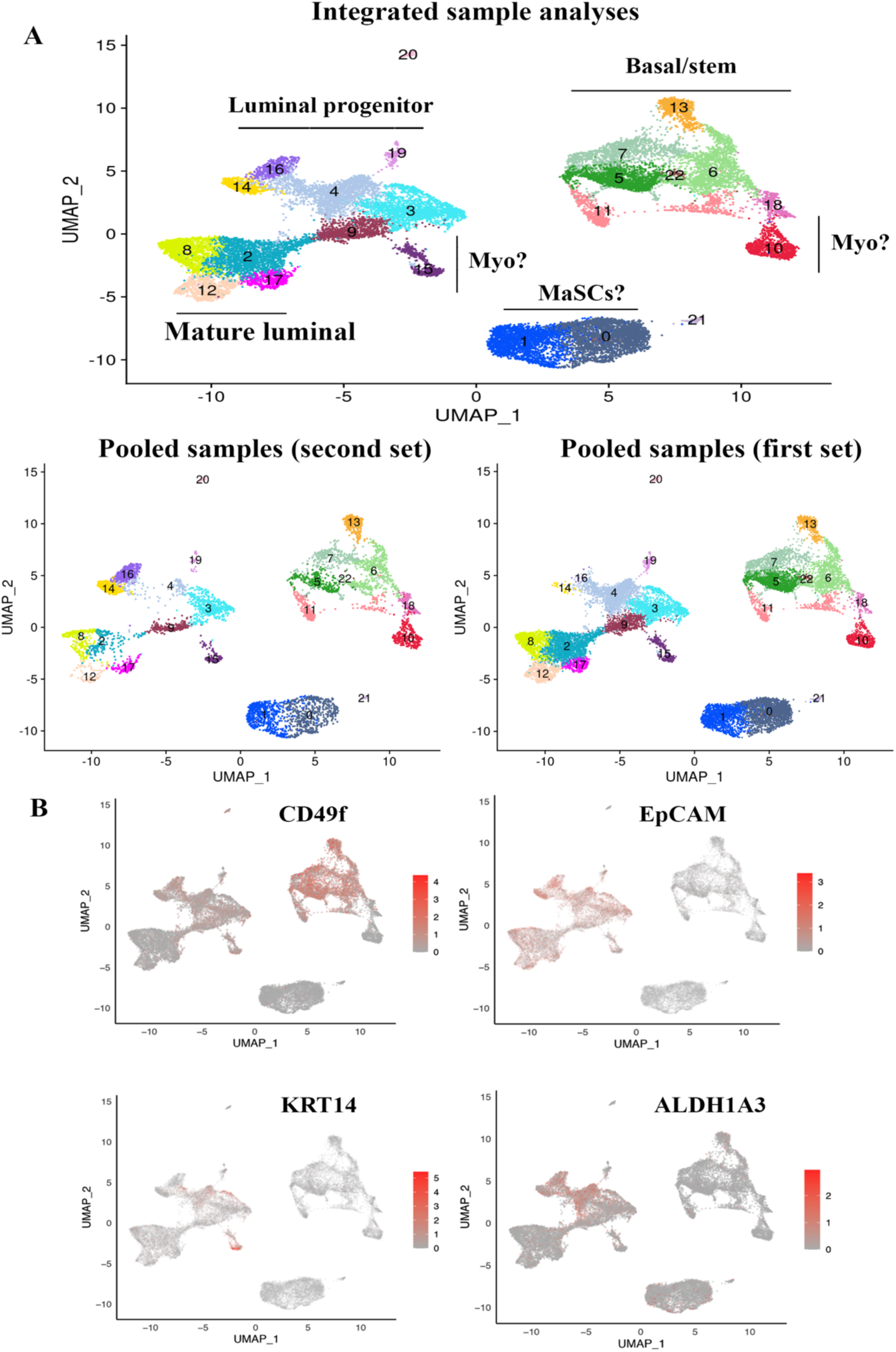
Recharacterization of epithelial cells of the normal breasts with additional samples. A) Combined integrated analyses that included samples in Figure 1, a new sample from an Asian (Chinese), and pooled five new samples. There were 23 clusters of cells, which can be subdivided into three major groups of basal/stem, luminal progenitor, and mature luminal cells. Potential myoepithelial cells (Myo) and mammary stem cells (MaSc) distinct from basal/stem cells are also indicated in the bottom. The bottom panel shows distribution patterns of cell clusters in five samples of the first set and the five pooled samples of the second set. Clusters in individual samples are shown in Figure S1B. Expression patterns of various markers that are used to subclassify clusters are shown Figure S2. B) *CD49f*, *EpCAM*, *ALDH1A3,* and *KRT14* expression in various clusters.

To further document reproducibility, we analyzed a surgical sample from a 33-year old Hispanic BRCA1 carrier and a core biopsy of a healthy Asian (Chinese) woman. BRCA1 sample was analyzed from cryopreserved tissues and we included duplicate samples because of availability of large starting material. One sample was prepared as above involving both enzymatic and mechanical disruption, whereas another sample utilized only digestion with gentle hyaluronidase/collagenase cocktail from Stem Cell technologies. Sample preparation using gentle hyaluronidase/collagenase yielded lower number of basal cells compared to the method that involved both enzymatic and mechanical disruption. Nonetheless, we did not observe any clusters unique to the BRCA1 mutated sample (Figure S1B). The breast tissue from Asian/Chinese donor showed a disproportionately higher number of cells within basal/stem characteristics compared to other samples.

### Gene expression patterns in Clusters 11 and 12 are similar to N19 and N0-N1, respectively, of the new analysis

Since cluster 11 of the first analysis, despite being a minor cluster, was enriched for genes associated with cell cycle and chromosome segregation, we next investigated which among the clusters in the second analysis are enriched for cluster 11 genes. In addition, since N0-N1 clusters grouped farther away from the remaining clusters, we investigated their relationship to the first analyses. A significant number of genes in cluster 11 were also enriched in cluster N19 of the second analysis, whereas cluster 12 genes were enriched in clusters N0 and N1 (Figure 5). Similar to C11, N19 expressed *MKI67, BIRC5* and *PCLAF*. Cluster N19 cells also expressed higher levels of *APOBEC3B*, which is considered a driver of mutations in cancer (Olson et al., 2018). Similar to C12, N0 and N1 expressed PTGDS and IGF1. These two clusters are likely enriched for unique stem cells as these cells expressed higher levels of *ZEB1, EGFR, CD44* and low levels of various keratins (Morel et al., 2017). A fraction of these cells as well as cluster N6 cells among basal/stem cell group were *PROCR+*, another mammary stem cell marker (Wang et al., 2014). Note that none of 23 clusters expressed mesenchymal stem cell markers such as *CD90*, *CD73* and *CD105* (Kfoury and Scadden, 2015).

**Figure 5:**
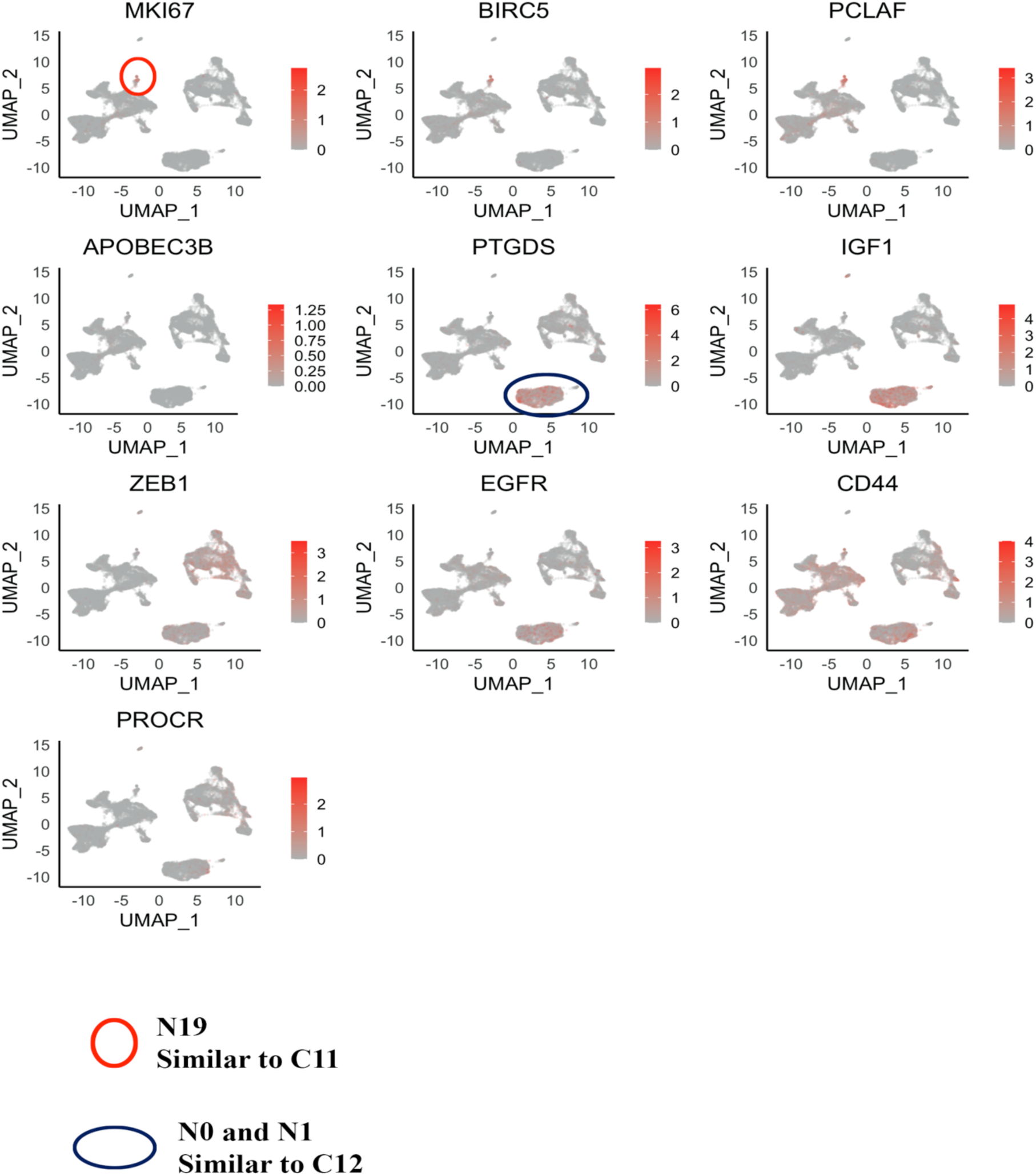
Gene expression in clusters N19 and N0-N1 of Figure 4 overlap with unique genes in C11 and C12, respectively. A) *MKI67, BIRC5*, and *PCLAF*, which are all overexpressed in cluster 11 (Figure 1D), are enriched in N19. B) *PTGDS* and *IGF1*, which are overexpressed in cluster 12 (Figure 1D), are enriched in N0-N1 clusters. This cluster also expresses *ZEB1* and *EGFR*.

### Cluster 11 and cluster 12-specific genes are enriched in breast cancer

Although it is not possible to definitively link cancer to a specific cell type from which it may originate (Gupta et al., 2019), it is likely that genes that define a epithelial cell cluster are enriched in tumors that originate from that cluster of epithelial cells. To determine such relationships, we compared gene scores of each epithelial cluster with METABRIC and TCGA breast cancer gene expression datasets (Cancer Genome Atlas, 2012; Curtis et al., 2012). To increase number of samples per cluster, we used clusters 0-12 of the first analysis instead of second analysis clusters N0-N22. Furthermore, since three clusters among CD49f+/EpCAM- showed modest gene expression differences, we analyzed them together as cluster 5a. In the CD49f-/EpCAM+ population, we combined clusters 1, 3, and 4 as cluster 1a. With both datasets, the highest number of breast cancers mapped to the minor cluster 12. This was followed by cluster 1a and cluster 11 in both datasets (Figure 6A). Consistent with the literature that the majority of breast cancers originate from luminal progenitor cells (Lim et al., 2009), gene expression in the highest number of basal breast cancers overlapped with cluster 11 followed by clusters 12, 2, and 6 (Tables S3 and S4). In METABRIC dataset, every intrinsic subtype of breast cancer was found to be enriched for genes of cluster 12. Survival analyses indicated significant differences in overall survival between breast cancers with overlapping signature of specific normal epithelial clusters in METABRIC but not in TCGA dataset. Note that METABRIC dataset is much larger than TCGA dataset. In the TCGA dataset, we observed tumors enriched for cluster 1a (mature luminal) signature associating with better disease-free survival (DFS) than tumors enriched for cluster 2 (luminal progenitor) signature (Figure 6B). Within luminal progenitor clusters, tumors enriched for cluster 12 signature were associated with better DFS than tumors with signature of cluster 2. Although tumors enriched for signatures of cluster 1a and cluster 12 did not show significant differences in overall survival in the TCGA dataset, tumors with cluster 12 signature were associated with better overall survival than cluster 1a in METABRIC dataset (Figure 6C).

**Figure 6:**
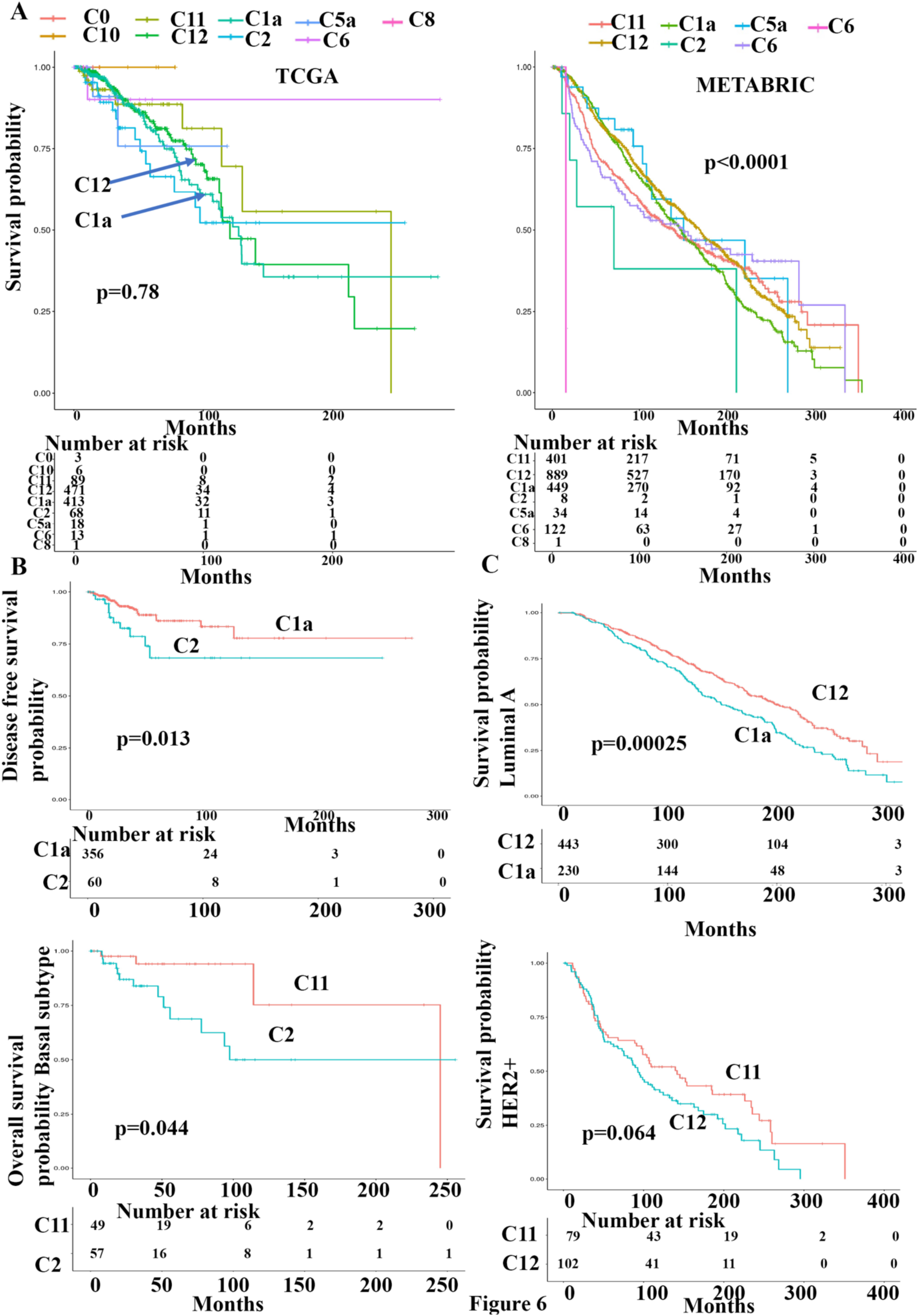
Breast cancer subtype specific expression of cluster-signature genes. Breast cancer gene expression data in TCGA (left) and METABRIC datasets were analyzed for enrichment of cluster-specific genes described in Table S2. Clusters 1, 3 and 4 were combined to create Cluster 1a because of limited differences. Similarly, clusters 5, 7 and 9 were clubbed to create cluster 5a. PAM50 intrinsic subtype classifiers were used to subdivide breast cancers into luminal A, luminal B, HER2, and basal subtypes. Enrichment of cluster-specific genes in these subtypes of breast cancer were further analyzed. Additional data can be found in Figure S3.

We used PAM50 classifier to determine a relationship between intrinsic subtypes and normal breast epithelial clusters. In both datasets, ∼50% of luminal A breast cancers contained cluster 1a signature and remaining were enriched for cluster 12 signature (Figure S3). In the METABRIC dataset, 673 out of 679 Luminal A breast cancers were assigned to cluster 1a and cluster 12 clusters suggesting two distinct cell-of-origin of Luminal A breast cancer. Luminal A breast cancers with cluster 12 signature were associated with better overall survival compared to luminal A with cluster 1a signatures in METABRIC dataset but not with TCGA dataset (Figure 6C and Figure S3). Disproportionately higher percentage of luminal B breast cancers carried cluster 1a signature compared to cluster 12 signature in both datasets (Figure S3). Cluster 11 signature was found in ∼50% luminal B breast cancers of METABRIC dataset (Table S3 and S4). Failure to find similar association between luminal B and cluster 11 in TCGA dataset could be due to low number of tumors with cluster 11 signature in this dataset. HER2+ breast cancers were enriched for cluster 1a, cluster 12 and cluster 11 (only in METABRIC) signatures, whereas basal-like breast cancers were enriched predominantly for cluster 11 signature. With respect to outcome, interactions between cell clusters and oncogenic signals may determine outcome. For example, while luminal B breast cancers with C1a signatures displayed better outcome than luminal B breast cancers with cluster 12 signatures, opposite outcomes were observed if the tumors were HER2+ (Figure S3). With respect to mutation frequency, breast tumors enriched for cluster 11 signature had highest number of mutations per tumor compared to other cluster enriched tumors (Table S4). This table also lists names of genes mutated in more than 10% of tumors.

Basal-like breast cancers with cluster 11 signatures were associated with better overall survival compared to basal-like breast cancers with cluster 2 signature in the TCGA dataset, although sample size is bit small (Figure 6B). Interestingly, Kaplan-Meier curve of basal breast cancers enriched for cluster 2 and cluster 12 signatures showed slopes similar to what is observed with triple negative breast cancer patients with most events occurring within first 100 months (Figures 6 and S3). Cluster 1a or cluster 5a signatures were rarely represented in basal breast cancers.

### TBX3 and PDK4 expression patterns determine subtypes of ER+ breast cancers

We observed expression of *ESR1* in three clusters (1, 3, and 4) of the normal breasts, which are characterized by differential expression of *TBX3, PDK4*, and *GATA3* (Figure 3C). While cluster 4 expressed similar levels of *ESR1*, *TBX3* and *PDK4*, *PDK4* expression was lower in cluster 3. Compared to cluster 4, cluster 1 expressed lower levels of both *TBX3* and *PDK4*. *GATA3* expression was highest in cluster 3 followed by clusters 1 and 4. Since ER-related pioneer factor activity of GATA3 has already been established (Zaret and Carroll, 2011), we focused our studies on TBX3 and PDK4 proteins. Both PDK4 and TBX3 expression are linked to anti-estrogen response (Razavi et al., 2018; Walter et al., 2015). To determine whether there is a relationship between clinical progression and ER/TBX3/PDK4 status, we immunostained 586 breast tumor containing TMA with 15-years of follow up for TBX3 and PDK4. As expected, while TBX3 staining was predominantly nuclear, PDK4 expression was cytoplasmic (Figure 7A). Detailed univariate and multivariate statistical analysis report of TBX3 and PDK4 expression generated in a blinded manner to the statistician, is presented in Star methods file and only statistically significant data are described below.

**Figure 7:**
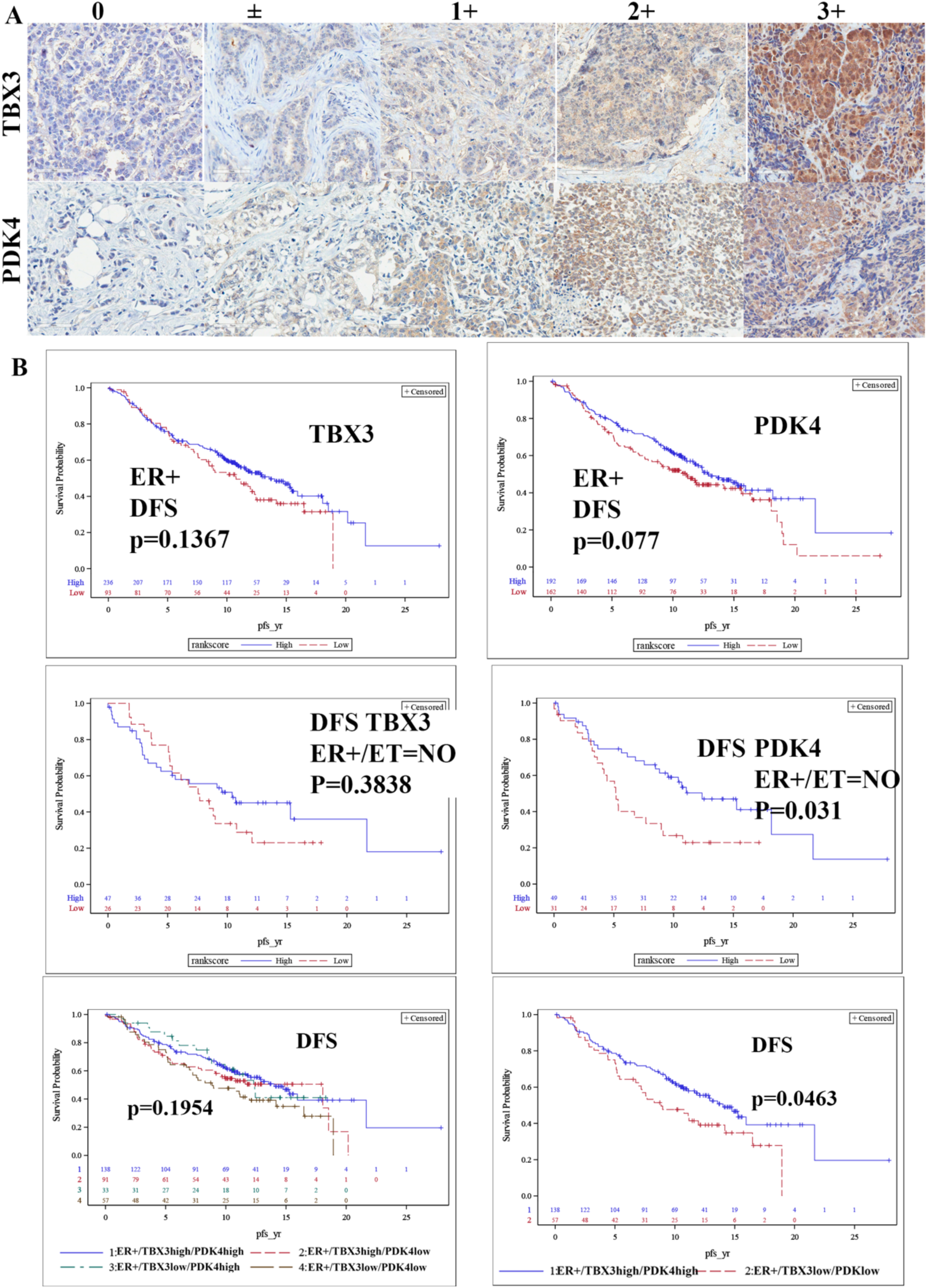
PDK4 and TBX3 enable further classification of ER+ breast cancers. A) Immunohistochemistry of breast TMA for PDK4 and TBX3. B) ER+ breast cancers expressing lower levels of PDK4 compared to tumors with higher PDK4 and not received endocrine therapy were associated with poor disease free survival (DFS). Similarly, ER+ tumors expressing lower levels of both TBX3 and PDK4 compared to tumors expressing higher levels of PDK4 and TBX3 were associated with poor DFS.

PDK4 expression levels correlated with ER+/PR+/HER2-, whereas TBX3 expression levels correlated with tumor grades and stage. Higher grade/stage tumors had higher TBX3 compared to lower grade/stage tumors. In multivariable models treating the PDK4 H-score as dichotomous, H-score category was significant for tumors that are ER+, and patients with ER+ tumors and on endocrine therapy (Table 1). For the results that were significant, the higher PDK4 H-score was correlated with lower survival. In multivariable models treating the H-score as continuous variable, the H-score was significant result in patients who were ER+, on endocrine therapy, were ER+ and on endocrine therapy, and for patients who were ER+/PR+/HER2-. A few Kaplan-Meier curves of disease-free survival analyses are shown in Figure 7B.

**Table 1:**
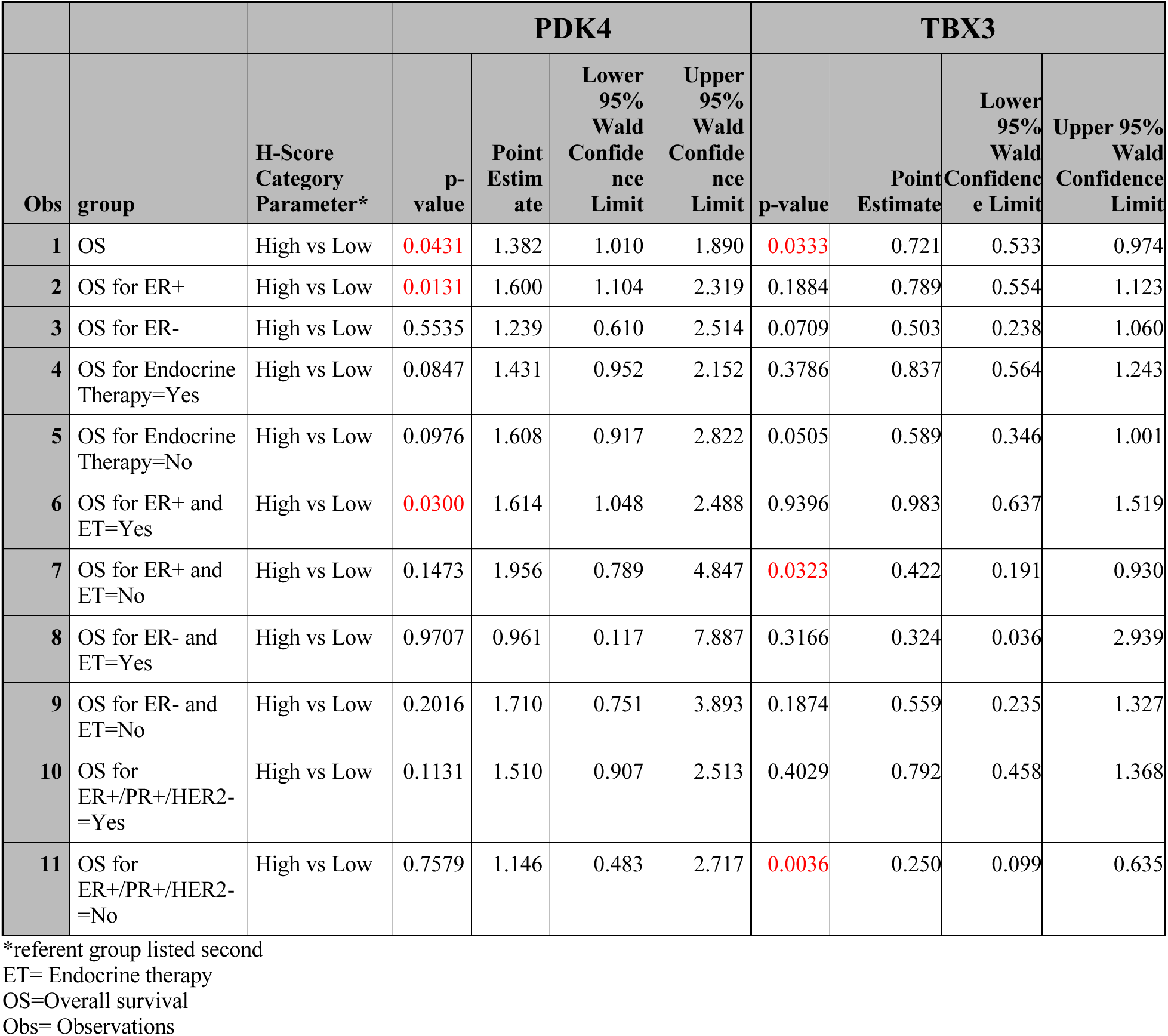
Relationship between PDK4 and TBX3 H-scores and outcome.

In the case of TBX3, in multivariable models treating the H-score as dichotomous, H-score category was significant for patients with tumors that were ER+ and not on endocrine therapy and patients who were not ER+/PR+/HER2- (Table 1). For the results that were significant, the lower TBX3 H-score was correlated with lower survival. In multivariable models treating the H-score as continuous, the H-score was significant in patients who were not ER+/PR+/HER2-.

Since our TMA had more than 300 ER+ cases, we were able to perform subgroup analyses that included all three markers: ER, TBX3 and PDK4. The analyses included ER+/TBX3+/PDK4+, ER+/TBX3+/PDK4_low_, ER+/TBX3_low_/PDK4_high_, and ER+/TBX3_low_/PDK4_low_. Although we did not observe any difference in overall survival between these groups, disease-free survival was shorter for patients with the tumors displaying ER+/TBX3_low_/PDK4_low_ expression patterns compared to ER+/TBX3+/PDK4+ expression patterns (Figure 7B). Among 399 ER+ tumors, 138 displayed ER+/TBX3+/PDK4+ characteristics, whereas 57 showed ER+/TBX3_low_/PDK4_low_ characteristics. These results indicate that ER+ breast cancers can be subclassified into at least four distinct subtypes based on TBX3 and PDK4 expression patterns, potentially representing four different cell-of-origin of ER+ breast cancers.

## Discussion

In this study, we present evidence for the presence of at least 13 different clusters of epithelial cells in the normal breast and we suggest that the majority of breast cancers contain gene expression signatures that overlap with gene expression signatures of two minor clusters of the luminal progenitors and the rest with mature luminal cells. It is possible that cells in these clusters (cluster 11, cluster 12 and cluster 1a that combines clusters 1, 3, and 4) are the cancer-prone population of normal breast epithelial cells. Finding of this study may permit breast cancer classification based on cell-of-origin of tumors. In this respect, a recent study involving 10,000 tumors from 33 cancer types showed that cell-of-origin patterns dominate in the molecular classification of tumors (Hoadley et al., 2018). Although each intrinsic subtype of breast cancer ostensibly has a distinct cell-of-origin in the breast stem-progenitor-mature cell hierarchy (Prat and Perou, 2009), we observed a cluster-enriched signature being represented in more than one intrinsic subtype and an intrinsic subtype being represented in more than one cluster of epithelial cells. Existing technologies do not permit experimental validation of cells-of-origin of tumors but the use of the techniques such as single cell RNA-seq may allow further refinement of cancer classification based on their presumptive cell-of-origin.

### Complexities in breast epithelial cell types; past and the present

Since scRNA-seq technology is still an evolving field requiring constant improvement in technologies starting from source of tissues to dissociation protocols, sequencing techniques, and bioinformatics tools (Lim et al., 2020), it is likely that clusters that we identified here will undergo further refinement in the future. Thus, it is appropriate to compare what has been done in the past to the current data. There has been a limited number of studies that utilized scRNA-seq technology to subclassify breast epithelial cells. Two publications to our knowledge utilized tissues from reduction mammoplasty samples and cells were either purified by flow cytometry or grown under organoid cultures prior to single cell sequencing (Nguyen et al., 2018; Rosenbluth et al., 2020). Although our source of tissue and methodology differed significantly than these studies as we used breast tissues from healthy women and were able to prepare single cell cDNA within two hours of tissue collection, there were several overlapping observations. For example, the L2 luminal differentiated cell cluster described by Nguyen et al., and luminal differentiated cluster 3 in our study are enriched for *ANKRD30A* (Nguyen et al., 2018). Similarly, luminal progenitor cluster L1 in that study and our luminal progenitor subcluster 10 are enriched for the expression of *SLPI* and *ANXA1*. Similar to that study, our luminal mature cell subcluster 4 was enriched for *PIP*. A basal subcluster identified by Nguyen et al., (Nguyen et al., 2018) and our cluster 5, which is basal, both expressed *TCF4*. There were few differences. Nguyen et al (Nguyen et al., 2018) suggested that *ACTA2*, which codes for α-SMA, distinguishes basal/myoepithelial cell from other cell types. Although we observed expected enrichment of *ACTA2* in basal subclusters, it is also expressed in a distinct subcluster of luminal progenitor cells (Cluster 6). This cluster is enriched for *MYLK*, an actin binding protein, regulated by ZEB1/miR-200 feedback loop associated with epithelial to mesenchymal transition (Sundararajan et al., 2015). It is possible that cluster 6 cells correspond to naturally occurring luminal/basal hybrid cells that can trans-differentiate based on environmental cues or to mixed luminal/basal lineage cells described in the mouse mammary gland (Pal et al., 2017).

Two studies have described distinct cell types in the mouse mammary gland during different stages of development (Bach et al., 2017; Pal et al., 2017). Similar to differences in the number of epithelial subgroups identified in two human studies (three by Nguyen et al., and 13 by us), Bach et al identified 15 clusters of mouse mammary epithelial cells through single cell sequencing of sorted EpCAM+ cells (Bach et al., 2017). Pal et al identified seven clusters of epithelial cells (Pal et al., 2017). There is some overlap in genes expressed in specific clusters of the mouse mammary gland and human identified in our study. Similar to our results, Pal et al showed *ACTA2* expression in both basal cells and two small subclusters of luminal cells (Pal et al., 2017). *CXCR14* expression was found in a subset of luminal progenitor/intermediate cells and basal cells. Gene expression in hormone sensing cells that included *ESR1* and *FOXA1* showed similarity in expression between Bach et al., and our studies (Bach et al., 2017). Pal et al identified *SFRP1* as a marker of pre-pubertal mammary epithelial cells, which decreased after puberty (Pal et al., 2017). In our analysis, *SFRP1* expressing cells were luminal progenitor cells. We noted one major difference between mouse and human mammary epithelial cells. In mouse mammary gland, *TSPAN8* expressing cells are considered quiescent stem cells (Fu et al., 2018), but we found *TSPAN8* expression to be restricted to cells in cluster 4, which is a mature luminal subcluster. Thus, few of the stem cell markers may show species-specific variability.

Basal, luminal progenitor and mature luminal cells are defined using cell surface markers CD49f and EpCAM (Visvader and Stingl, 2014). However, these markers are not ideal for in situ estimation of three cell types. A closer look at genes enriched in each cluster and their signature genes revealed that *XBP1* is expressed predominantly in luminal mature cells, whereas *SFRP1* is expressed predominantly in luminal progenitor cells. Multiple genes are enriched in basal cell clusters including *CD96, MECOM, CLDN5,* and *CD93*. These genes can be used in the future for *in situ* estimation of composition of the breast.

### Gene signatures of epithelial clusters and their relevance to breast cancer

Our single cell studies confirmed prior reports of basal breast cancers originating from luminal progenitor cells. The majority of basal breast cancers in both TCGA and METABRIC datasets were enriched for genes found in luminal progenitor clusters 2, 11 and 12. Two minor clusters that we identified, clusters 11 and 12, displayed several unique characteristics. Cluster 11 is enriched for the expression of proliferation-associated genes including *MKI67, CDK1*, and *TK1*, apoptosis regulators (*BIRC5, DEPDC1*). It co-expressed *TOP2A*, *BIRC5* and *NEK2*, which form a functional protein-protein network in luminal A breast cancer (Nuncia-Cantarero et al., 2018). In our second set of analyses, we found this cluster to be enriched for *APOBEC3B* and its elevated expression is associated with higher mutation rate (Olson et al., 2018). Indeed, breast tumors with cluster 11 signatures displayed the highest number of mutations per tumor compared to tumors enriched for other genes of other clusters. In fact, 15 out of 34 genes of this cluster have been shown to be present in a rarest cluster of breast tumor cells with highly malignant phenotype (Gao et al., 2017). Although this cluster represented 1.5% of epithelial cells, ∼20% of breast cancers in TCGA and METABRIC datasets were enriched for genes of this cluster. There is a trend of higher number of cluster 11 cells in young donors, but this observation needs to be confirmed with large number of samples (Table S2). Collectively, these observations reinforces the possibility of these rare subpopulation of cells being the cell-of-origin of significant numbers of breast cancers. Based on the expression of MKI67 as well as pathway analyses, cluster 11 contain rapidly proliferating cells, which may predispose cells for aberrant chromosome segregation and mutations.

Cluster 12 was enriched for *IGF1, MEG3, PTGDS* and *SRPX*. Among these genes, *MEG3* codes for a linker RNA and is dysregulated in multiple cancers (He et al., 2017). Although this cluster appeared minor in our first analyses, when we sequenced additional samples and did combined analyses, the number of cells representing this cluster increased and we were able to identify additional marker genes including *ZEB1* and *TCF4*. Co-expression of *TCF4* and *ZEB1* cells indicate that these cells are similar to a subset of basal cells described by Nguyen et al., but these cells lack the expression of *CD49f* to be considered as basal cells (Nguyen et al., 2018). To a certain extent, these cells are similar to mammary stem-like cells described by Morel et al., and ZEB1+ cells we described previously (Morel et al., 2017; Nakshatri et al., 2019). Because the gene signatures from this cluster are heavily represented in almost all intrinsic subtypes of breast cancer, additional studies are needed to determine the role of these cells in tumorigenesis.

With respect to ER+ breast cancers, these cancers can originate from both luminal mature and luminal progenitor cells as both luminal A and luminal B breast cancers displayed gene signatures enriched in cluster 1a (clusters 1, 3 and 4), and clusters 11 and 12. There appears to be quantitative differences in *ESR1* between these clusters as cluster 1a expressed highest level of *ESR1* followed by clusters 11 and 12 (Table S2). Why ER+ breast cancers with a luminal progenitor gene expression pattern compared to mature luminal gene expression is associated with better outcome is an intriguing question that needs to be explored. Using an independent TMA, we were able to further subclassify mature luminal cell-derived ER+ tumors based on TBX3 and PDK4 expression. While the role of TBX3 in ER activity and its mutations in ER+ lobular carcinomas have been described in the literature (Ciriello et al., 2015; Razavi et al., 2018), there are no studies that functionally linked PDK4 to ER. PDK4 is a cytoplasmic kinase involved in the TCA cycle and induces metabolic changes in transformed cells (Coloff and Brugge, 2017). Whether it also modulates transcription by targeting transcription machinery needs further investigation.

ScRNA-seq as well as single cell protein sequencing have been used to subclassify breast cancers and to identify treatment resistant population. For example, Karaayvaz et al described an aggressive disease-associated gene signature related to glycosphingolipid metabolism by scRNA-seq of six TNBCs (Karaayvaz et al., 2018). However, pathway analyses of our cluster enriched genes did not identify a normal breast epithelial cluster enriched for this pathway. Similarly, signatures derived through scRNA-seq of a chemoresistant subpopulation of TNBCs did not show overlap with any of our normal cell clusters (Kim et al., 2018). Genes in the breast cancer-specific RNA signature that detect circulating tumor cells did not show overlap with any specific cluster, but genes like *PIP, CXCL14*, and *SFRP2*, which are markers of circulating tumor cells are enriched in clusters 4, 6 and 12, respectively (Kwan et al., 2018). Thus, genomic aberrations and transcription programming rather than cell-of-origin may have given rise to drug resistant and metastatic subpopulations of tumor cells.

A recent single cell proteomics observed expression of EMT-associated proteins in triple negative breast cancers as well as luminal A and luminal B breast cancers (Wagner et al., 2019). Cluster 12 (and its equivalent in the second analyses N0-N1) naturally expressed higher levels of EMT-associated genes and signature of this cluster were enriched in all intrinsic subtypes. Thus, expression of EMT-associated genes may not be an acquired phenomenon in all breast cancers.

From a basic research point of view, results presented here provides an opportunity to determine whether a gene is truly differentially expressed in tumors compared to normal, as the expression pattern of a specific gene in the tumor could be a reflection of its cell-of-origin. Using single cell RT-PCR of tumor adjacent normal and tumor cells from the same individual, we had previously demonstrated that elevated expression of few genes in tumor can be attributed to cell-of-origin of tumor instead due to tumor-specific genomic aberration (Anjanappa et al., 2017). Resources created here, which will be made available to researchers and can be mined for expression patterns of specific genes in various epithelial clusters of the normal breasts using the tool such as Loupe Browser of 10X Genomics. For example, genes such as *MUCL1*, *PIP*, *TP63* are expressed in a specific subpopulation of normal breast epithelial cells and their overexpression in tumors may reflect cell-of-origin of tumors instead of tumor-specific aberrant expression. This approach would also allow streamlining of drug discovery efforts by focusing on targets that are truly differentially expressed in tumors due to genomic aberrations.

## ACKNOWLEDGMENTS

We thank the countless number of women who donated normal and malignant breast tissues for research. We also thank the volunteers who facilitated this tissue collection. Special thanks to members of the Komen Tissue Bank including Ms. Jill Henry, Alison Hughes, Pam Rockey, Julia Rose von Arx, Rana German and Dr. Natascia Marino as well as the IU Simon Cancer Center tissue procurement facility for providing tissues and related data. **Funding:** The Catherine Peachy Fund of the Heroes Foundation family (HN), Breast Cancer Research Foundation (HN), Chan-Zuckerberg Initiative Human Atlas Project (HN, AS, YL), Susan G. Komen for the Cure (AS), and Vera Bradley Foundation for Breast Cancer Research (IUSM). Walther Cancer Institute provided support to Cancer Bioinformatics Core.

## AUTHOR CONTRIBUTIONS

Conception and design: HN

Development of methodology: PBN, PCM, XX, LS, JW, GS, HG, YL, HN

Acquisition of data: PBN, PCM, AC, GS.

Analysis and interpretation of data: PBN, AMS, HG, YL, LS, JW, HN

Writing, review and/or revision of the manuscript: HG, LS, AMS, HN

Administrative, technical, or material support: AMS, HN

Study supervision: HN

## SUPPLEMENTAL INFORMATION

Supplemental information includes 4 figures and four tables and can be found online.

## STAR * METHODS

| **REAGENT or RESOURCE** | **SOURCE** | **IDENTIFIER** |
| --- | --- | --- |
| **ANTIBODIES** |  |  |
| PDK4 | Abcam | Ab71240 |
| TBX3 | Abcam | Ab99302 |
| **REAGENTS** |  |  |
| COLLAGENASE/HYALURANIDASE | Stem cell technologies |  |
| Cryopreservation media | LONZA | 12-132A |
| ROCK Inhibitor Y-27632 | TOCRIS | 1254 |
| Gentle Collagenase/Hyaluronidase | StemCell technologies | 07919 |
| Tumor dissociation kit (human) | Miltenyi Biotech | 130-095-929 |
| Red cell lysis buffer | Miltenyi Biotech | 130-094-183 |
| Debris removal kit | Miltenyi Biotech | 130-109-398 |
| Chromium Single cell 3'reagents | 10X Genomics | CG00052 Rev B or CG000183 Rev C |
| Bioanalyzer HSDNA CHIP | G2943CA | Agilent |
| **SOFTWARES AND ALGORITHMS** |  |  |
| CellRanger 2.1.0 or 3.0.2 | 10X Genomics | http://support.10xgenomics.com |
| Loupe Browser | 10X Genomics | [https://support.10xgenomics.com/single-cell-gene-](https://support.10xgenomics.com/single-cell-gene) |
|  |  | expression/software/visualization/la |
|  |  | test/installation |
| SAS Version 9.4 |  | www.sas.com |

### CONTACT FOR REAGENT AND RESOURCE SHARING

Further information and requests for reagents should be directed to and will be fulfilled by the corresponding author: Harikrishna Nakshatri

### EXPERIMENTAL MODEL AND SUBJECT DETAILS

#### Normal breast tissues

All breast tissues from healthy women were collected by the Komen Normal Tissue Bank with informed consent and with the approval from the institutional review board. Standard operating procedure for tissue collection is described on the Komen Tissue Bank website. Per standard operating procedures, the normal breast biopsies were always collected from the upper outer quadrant of the breasts. Within an average of six minutes from the time of biopsy, tissues were either placed in a growth media and transported to the lab for immediate single cell sequencing or cryopreserved for single cell sequencing at a later date. Our cryopreservation protocol has been described previously (Nakshatri et al., 2015). Briefly, tissue was minced and placed in one ml of 50% growth media and 50% Lonzo freezing media with 2 micromolar ROCK inhibitor. Vials with tissues were placed in CoolCell Containers (Nalgene) and placed in a −80°C freezer overnight and then in liquid nitrogen. Tissue specimens were thawed rapidly at 37°C and then washed extensively in growth media prior to dissociation. Tissue specimens represented women of different race, age, parity, menstrual phase, and BMI (Table S1).

#### Tissue dissociation procedure, cDNA library preparation and sequencing

We used the human tumor dissociation kit from Miltenyi Biotech (130-095-929) to generate single cells from tissue specimens. Red blood lysis buffer (130-094-183) and debris removal solution (130-109-398) were used as needed to improve purity of single cells. Viability and single cell status were determined via trypan blue staining and phase contrast microscopy. Samples with 80% or more viability were utilized for the subsequent steps. Cells were suspended at ∼100-800 cells/µl depending on sample and subjected to cDNA library generation using 10 X Genomics V2 (initial study) or V3 (second set of samples) Chromium Single Cell 3’ Reagents (CG00052 Rev B and CG000183 Rev C, respectively). We used HSDNA Chips on the Bioanalyzer from Agilent technologies (G2943CA) to quantify cDNA. cDNA was amplified using Chromium TM Single cell Library kit v2 or v3. The resulting libraries were sequenced on Illumina NovaSeq 6000 to a read depth of ∼50,000 reads per cell. 26 bp of cell barcode and UMI sequences, and 91 bp RNA reads were generated for the libraries made with the V2 kit; and 28 bp plus 91 bp paired-end for the libraries with the V3 kit.

#### Analysis of scRNA-seq sequence data

CellRanger 2.1.0 or 3.0.2 (http://support.10xgenomics.com/) was utilized to process the raw sequence data generated. Briefly, CellRanger used bcl2fastq (https://support.illumina.com/) to demultiplex raw base sequence calls generated from the sequencer into sample-specific FASTQ files. The FASTQ files were then aligned to the human reference genome GRCh38 with RNA-seq aligner STAR. The aligned reads were traced back to individual cells and the gene expression level of individual genes were quantified based on the number of UMIs (unique molecular indices) detected in each cell.

The filtered gene-cell barcode matrices generated with CellRanger were used for further analysis with the R package Seurat version 2.3.1 and development version 3.0.0.9000 with R studio version 1.1.453 and R version 3.5.1 (Butler et al., 2018; Stuart et al., 2019). Quality control (QC) of the data was implemented as the first step in our analysis. We first filtered out genes that were detected in less than five cells and cells with less than 200 genes. To further exclude low-quality cells in downstream analysis we used the function isOutlier from R package scater together with visual inspection of the distributions of number of genes, UMIs, and mitochondrial gene content (McCarthy et al., 2017). Cells with extremely high or low number of detected genes/UMIs were excluded. In addition, cells with high percentage of mitochondrial reads were also filtered out. After removing likely doublets/multiplets and low-quality cells, the gene expression levels for each cell were normalized with the NormalizeData function in Seurat. To reduce variations sourced from different number of UMIs and mitochondrial gene expression, we used the ScaleData function to linearly regress out these variations. Highly variable genes were subsequently identified.

To integrate the single cell data from individual donor samples, functions FindIntegrationAnchors and IntegrateData from Seurat v3 were implemented. The integrated data was then scaled and PCA was performed. Clusters were identified with the Seurat functions FindNeighbors and FindClusters. The FindConservedMarkers function was subsequently used to identify canonical cell type marker genes. Cell cluster identities were manually defined with the cluster-specific marker genes or known marker genes. The cell clusters were visualized using the Uniform Manifold Approximation and Projection (UMAP) plots. To help interactively explore various gene expression pattern across cell clusters, the UMAP cell embeddings and cell cluster information generated from Seurat analysis were imported into 10X genomics Loupe Browser (https://support.10xgenomics.com/single-cell-gene-expression/software/visualization/latest/installation). R packages ggplot2 and Seurat FeaturePlot were used to generate feature plots to visualize specific gene expression across clusters. R package ComplexHeatmap was used to generate the heatmaps (Gu et al., 2016; Wickham, 2016).

#### TCGA and METABRIC dataset analyses

1741 genes in all 13 clusters of the first analyses are considered as signatures. Due to higher similarity of gene expression among clusters 1, 3, and 4, they are merged as one cluster 1a. Similarly, clusters 5, 7, and 9 are merged as one cluster 5a. Centroids of the nine resulting merged clusters were generated for Prediction Analysis of Microarray (PAM) algorithm using the 1741 genes (Parker et al., 2009; Tibshirani et al., 2002). Expression, clinical, and mutation data of METABRIC and TCGA BRCA were retrieved from cBioportal (Bhandari et al., 2019; Cerami et al., 2012; Ellrott et al., 2018; Gao et al., 2013; Gao et al., 2018; Hoadley et al., 2018; Liu et al., 2018; Pereira et al., 2016; Sanchez-Vega et al., 2018; Taylor et al., 2018). Expression data are median centered and applied to the PAM classifier based on the 1741 genes using Spearman’s rank correlation as distance. Each sample in METABRIC and TCGA BRCA data was assigned to one of the nine clusters. Relationship of each sample’s cluster membership with intrinsic subtypes and mutation were analyzed. Survival analysis of METABRIC and TCGA BRCA samples between clusters were analyzed using R package survminer v0.4.6 in R (Kassambara, 2019; Team, 2020; Wickham, 2016).

#### Breast cancer TMA and immunostaining for TBX3 and PDK4

The breast cancer TMA with ∼15-years of follow up has been described previously (Perkins et al., 2015). All tissue samples were collected following a detailed IRB approved protocol, informed patient consent, and HIPAA compliance protocol. Tissues were fixed overnight at room temperature in 10% NBF. A pathologist (GES) utilized light microscopy (Leica) to evaluate the staining in each tissue core (range from 0 to +3) to make sure there was no over staining and/or extensive background staining. The slides were imaged using the Aperio Scanscope CS. Computer-assisted morphometric analysis of digital images was performed using the Aperio Image Analysis software that came with the Aperio Whole Slide Digital Imaging System. The Positive Pixel Count algorithm was used to quantify the amount of a specific stain present in a scanned slide image. A range of color (range of hues and saturation) and three intensity ranges (weak, positive, and strong) were masked and evaluated. The algorithm counted the number and intensity-sum in each intensity range, along with three additional quantities: average intensity, ratio of strong/total number, and average intensity of weak positive pixels.

The algorithm was applied to an image by using the TMA Lab algorithm. This program allowed us to select each core, specify the input parameters, run the algorithm, and view/save the algorithm results. When using the Image Scope program, a pseudo-color markup image is also shown as an algorithm result. The H score was calculated using the Aperio TMA software algorithm. Formula is:

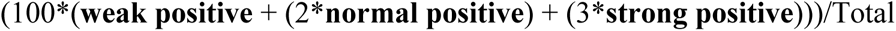

Approximately 80 to 90 breast biopsies in each of the 14 breast TMA immunostain were evaluated with TBX3 and PDK4 antibodies. Anti-PDK4 (ab71240) and anti-TBX3 (ab99302) antibodies were obtained from Abcam. The normal tissue controls (TMA orientation cores) were normal liver, cecum, kidney, spleen, tonsil, and heart.

With TBX3, immunostaining was seen in both the cytoplasm and nucleus in most tumor cells, and within few stromal cells in a few cases. In cases with inflammation, several of the lymphocytes (subset) were strongly stained. Staining patterns in tumor cells ranged from weak to moderate to strong. The two cores from the same patient in the arrays were often similar depending on the amount of fat and /or stroma in the core. Little to no background staining was seen in the other tissues in the core (vascular endothelial cells, smooth muscle cells, adipocytes, and fibroblasts).

With PDK4, immunostaining was seen in the cytoplasm of most tumor cells and in some cases, that of a few stroma cells. Lymphocyte staining was seen only in cases with inflammation. The tumor cells were weak to moderate to strong in staining with two cores from the same patient. This was consistent in all arrays with minimal background staining in the other tissues in the core (vascular endothelial cells, smooth muscle cells, adipocytes, and fibroblasts).

#### Statistical analyses of TMA data

For subjects with multiple tumor samples available, we included only the sample with the highest PDK4 or TBX3 H-score. Wilcoxon Rank Sum and Kruskal-Wallis tests were used to determine if PDK4 or TBX3 H-scores correlated with other tumor markers. Cox proportional hazards regression models were used to determine whether H-scores and other variables were related to overall and disease-free survival either univariately or in multivariable models. In these analyses, TBX3 H-scores were divided into low and high categories at the score of 27.91721 for overall survival (time from surgery to death or censoring) and disease-free survival (time from surgery to first recurrence or censoring, excluding patients with M1 stage at surgery). PDK4 H-scores were divided into low and high categories at the score of 19.41508 for overall survival (time from surgery to death or censoring) and 34.05692 for disease-free survival (time from surgery to first recurrence or censoring, excluding patients with distant metastatic diseases at surgery). These cutoff values were determined by using the maximum chi-square value for all score values between the 25th and 75th percentile as described previously (Perkins et al., 2015). PDK4 high/low and TBX3 high/low was included in all multivariable models. As a double check on the direction of the hazard ratio and as a more powerful test if the H-score effect was truly linear, we also fit multivariable models with the H-score as continuous.

We conducted subgroup analyses on overall survival using the ER-positive subgroup, endocrine therapy group, ER-positive on endocrine therapy, ER-negative, and ER+/PR+/HER2-. First, log-rank tests were done with the dichotomous H-score variable. Second, multivariable models with the H-score as dichotomous and then continuous were fit similar as was done in the main analyses. Analyses were conducted using SAS Version 9.4. An α level of 5% was used to determine statistical significance.

Individual marker statistical reports are below.

### Statistical Methods: PDK4

For subjects with multiple tumor samples available, we included only the sample with the highest PDK4 H-score. Wilcoxon Rank Sum and Kruskal-Wallis tests were used to determine if PDK4 H-scores were correlated with other tumor markers. Cox proportional hazards regression models were used to determine whether PDK4 H-scores and other variables were related to overall and disease-free survival either univariately or in multivariable models. In these analyses, PDK4 H-scores were divided into low and high categories at the score of 19.41508 for overall survival (time from surgery to death or censoring) and 34.05692 for disease-free survival (time from surgery to first recurrence or censoring, excluding patients with M1 stage at surgery). These cutoff values were determined by using the maximum chi-square value for all score values between the 25th and 75th percentile (http://www.pharmasug.org/proceedings/2012/SP/PharmaSUG-2012-SP12.pdf). PDK4 high/low was included in all multivariable models. As a double check on the direction of the hazard ratio and as a more powerful test if the H-score effect was truly linear, we also fit multivariable models with the H-score as continuous.

We conducted subgroup analyses on overall survival using the ER-positive subgroup, endocrine therapy group, ER-positive on endocrine therapy, ER-negative, and ER+/PR+/HER2-. First, log-rank tests were done with the dichotomous H-score variable. Second, multivariable models with the H-score as dichotomous and then continuous were fit similar as was done in the main analyses.

Analyses were conducted using SAS Version 9.4. An α level of 5% was used to determine statistical significance.

### Results

Of the 586 patients who were part of the TMAs available to be read (TMA1-14) for PDK4, 497 patients (85%) had PDK4 values available/readable. Clinical parameters of the subjects included in the TMAs studied are summarized in Table 1 with grouping for patients with or without PDK4 values. Age, Tumor Grade, Tumor Stage, Recurrences, and Disease Free Survival were significantly different between the two groups (Age higher in missing group, Grade 1 had more missing, T1 had more missing, and Fewer Recurrences in missing group, percentagewise with the DFS having a higher median in the missing group). Since the numbers for the missing PDK4 values were so low, these significant differences were not investigated further.

### Correlation of PDK4 H-score with other disease markers

We compared PDK4 H-score expression with ER, PR, HER-2/neu, Nodal stage, Tumor Stage or Grade. PDK4 levels were correlated with ER+/PR+/HER- (Table 2). Being ER+/PR+/HER- was significantly less than not being ER+/PR+/HER-.

### Overall Survival Analysis

#### Univariate

In univariate analyses, variables significantly related to overall survival in the Cox proportional hazards regression models were PR Status, HER-2 Status, ER+/PR+/HER2- Status, Tumor Grade, Tumor Stage, and Nodal Stage (Table 3a). PR-negative, having HER2-, not having ER+/PR+/HER2-, Higher Tumor Grade, Higher Tumor Stage, and Nodal Stage-positive were correlated with lower survival. PDK4 H-score was not related to overall survival (log rank test p-value 0.1165).

#### Multivariable

In the multivariate analysis (Tables 4aa and 4ab), models were run both with and without HER2- since there are a number of patients with missing results. ER+/PR+/HER2- was not included in the model since it would be correlated with the other variables. In the multivariate model with HER2- (Table 4aa), PR Status, and Nodal Stage were found to be significant. PR-negative and Nodal Stage-positive were correlated with lower survival.

In the multivariate model without ER+/PR+/HER2- (Table 4aa), PR Status, Tumor Grade, Tumor Stage, Nodal Stage, and the categorical PDK4 H-Score were found to be significant. PR-negative, Higher Tumor Grade, Higher Tumor Stage, and Nodal Stage-positive, and higher PDK4 H-score were correlated with lower survival.

#### Subgroup analysis

For unadjusted tests (i.e. univariately), the results were not significant for any of the subgroups (log rank test p-value 0.084 for ER-positive, 0.860 for ER-negative, 0.073 for patients on endocrine therapy, 0.464 for patients not on endocrine therapy, 0.053 for patients who were ER+ and on Endocrine Therapy, and 0.653 for ER-positive and not on endocrine therapy).

In multivariable models treating the H-score as dichotomous, H-score category was significant for ER-positive patients, and patients who were ER-positive and on endocrine therapy (Table 5a). For the results that were significant, the higher PDK4 H-score was correlated with lower survival. In multivariable models treating the H-score as continuous, the H-score was significant result in patients who were ER+, on Endocrine Therapy, were ER+ AND on Endocrine Therapy, and for patients who were ER+/PR+/HER2- (Table 6a). For the results that were significant, the higher PDK4 H-scores were associated with lower overall survival.

In addition, Kaplan-Meier plots are provided for the univariate analyses using the categorical PDK4 H-score for overall and the subgroup analyses.

### Disease Free Survival Analysis

#### Univariate

In univariate analyses, variables significantly related to disease free survival in the Cox proportional hazards regression models were PR Status, Tumor Grade, Tumor Stage, and Nodal Stage (Table 3a). PR-negative, Higher Tumor Grade, Higher Tumor Stage, and Nodal Stage-positive were correlated with lower disease free survival. PDK4 H-score was not related to disease free survival (log rank test p-value 0.0544).

#### Multivariable

In the multivariate analysis (Table 4b), PR Status, Tumor Grade, Tumor Stage, and Nodal Stage were found to be significant. PR-negative, Higher Tumor Grade, Higher Tumor Stage, and Nodal Stage-positive were correlated with lower disease free survival.

#### Subgroup analysis

For unadjusted tests (i.e. univariately), the results were significant for patients not on endocrine therapy (log rank test p-value=0.039), and patients who were ER-positive and also not on endocrine therapy (log rank test p-value=0.013). Other analyses were not statistically significant (log rank test p-value 0.078 for ER-positive, 0.675 for ER-negative, 0.849 for patients on endocrine therapy, and 0.766 for ER-positive and on endocrine therapy). For the significant results, the lower PDK4 H-score was correlated to lower disease free survival.

In multivariable models treating the H-score as dichotomous, H-score category was significant for patients who were ER+ AND not on Endocrine Therapy (Table 5b) where the patients with lower PDK4 H-score were correlated to lower disease free survival. In multivariable models treating the H-score as continuous, H-score was significant for patients who were on Endocrine Therapy, and patients who were ER-positive and also on Endocrine Therapy (Table 6b). For the significant results, the higher PDK4 H-score was correlated to lower disease free survival.

In addition, Kaplan-Meier plots are provided for the univariate analyses using the categorical PDK4 H-score for overall and the subgroup analyses.

**Table 1.** Description of the patients and characteristics of their tumors (n=586)

| **Characteristic** | **Have PDK4**<br>**(N=497)** | **Missing PDK4**<br>**(N=89)** | **p-value\*** |
| --- | --- | --- | --- |
| Age at Diagnosis, y |  |  | 0.0035 |
| Mean (SD) | 59.2 (15.08) | 64.2 (13.04) |  |
| Median | 58.15 | 66.14 |  |
| Range | 5.92-94.75 | 35.49-88.33 |  |
| Race, n(%) |  |  | 0.0927 |
| White | 385 (77.46%) | 77 (86.52%) |  |
| African American | 108 (21.73%) | 11 (12.36%) |  |
| Asian | 2 ( 0.40%) | 1 ( 1.12%) |  |
| Other | 1 ( 0.20%) | 0 |  |
| Unknown | 1 ( 0.20%) | 0 |  |
| PR Status, n(%) |  |  | 0.1818 |
| Negative | 158 (31.79%) | 23 (25.84%) |  |
| Positive | 295 (59.36%) | 61 (68.54%) |  |
| Unknown/Not Done | 44 ( 8.85%) | 5 ( 5.62%) |  |
| ER Status, n(%) |  |  | 0.2985 |
| Negative | 107 (21.53%) | 15 (16.85%) |  |
| Positive | 364 (73.24%) | 70 (78.65%) |  |
| Unknown/Not Done | 26 ( 5.23%) | 4 ( 4.50%) |  |
| HER-2/neu, n(%) |  |  | 0.3361 |
| Negative | 228 (45.88%) | 46 (51.69%) |  |
| Equivocal | 4 ( 0.80%) | 0 |  |
| Positive | 47 ( 9.46%) | 5 ( 5.62%) |  |
| Unknown/Not Done | 218 (43.86%) | 38 (42.70%) |  |
| Tumor Grade, n(%) |  |  | 0.0147 |
| I | 131 (26.36%) | 36 (40.45%) |  |
| II | 211 (42.45%) | 32 (35.96%) |  |
| III | 126 (25.35%) | 13 (14.61%) |  |
| IV | 1 ( 0.20%) | 0 |  |
| Unknown | 27 ( 5.43%) | 7 ( 7.87%) |  |
| Not Applicable | 1 ( 0.20%) | 1 ( 1.12%) |  |
| T Stage, *n*(%) |  |  | 0.0389 |
| T0 | 1 ( 0.20%) | 1 ( 1.12%) |  |
| T1 | 269 (54.12%) | 61 (68.54%) |  |
| T2 | 181 (36.42%) | 20 (22.47%) |  |
| T3 | 25 ( 5.03%) | 4 ( 4.49%) |  |
| T4 | 15 ( 3.02%) | 3 ( 3.37%) |  |
| TX/Unknown | 6 ( 1.21%) | 0 |  |
| N Stage, *n*(%) |  |  | 0.1192 |
| N0 | 295 (59.36%) | 62 (69.66%) |  |
| N1 | 126 (25.35%) | 18 (20.22%) |  |
| N2 | 29 ( 5.84%) | 1 ( 1.12%) |  |
| N3 | 10 ( 2.01%) | 2 ( 2.25%) |  |
| NX/Unknown | 36 ( 7.25%) | 6 ( 6.74%) |  |
| Not Applicable | 1 ( 0.20%) | 0 |  |
| M Stage, *n*(%) |  |  | 0.3296 |
| M0 | 255 (51.31%) | 55 (61.80%) |  |
| M1 | 17 ( 3.42%) | 1 ( 1.12%) |  |
| MX/Unknown | 223 (44.86%) | 33 (37.08%) |  |
| Not Applicable | 2 ( 0.40%) | 0 |  |
| Endocrine Therapy, *n*(%) |  |  | 0.5009 |
| Yes | 297 (59.76%) | 58 (65.17%) |  |
| No | 181 (36.42%) | 30 (33.71%) |  |
| Unknown/Missing | 19 ( 3.82%) | 1 ( 1.12%) |  |
| Death, *n*(%) |  |  | 0.0960 |
| Yes | 243 (48.89%) | 35 (39.33%) |  |
| No | 254 (51.11%) | 54 (60.67%) |  |
| Recurrence, *n*(%) |  |  | 0.0253 |
| Yes | 153 (30.78%) | 17 (19.10%) |  |
| No | 344 (69.22%) | 72 (80.90%) |  |
| TMA |  |  | 0.2725 |
| TMA-1 | 40 ( 8.05%) | 3 ( 3.37%) |  |
| TMA-2 | 30 ( 6.04%) | 8 ( 8.99%) |  |
| TMA-3 | 34 ( 6.84%) | 7 ( 7.87%) |  |
| TMA-4 | 32 ( 6.44%) | 7 ( 7.87%) |  |
| TMA-5 | 36 ( 7.24%) | 6 ( 6.74%) |  |
| TMA-6 | 31 ( 6.24%) | 7 ( 7.87%) |  |
| TMA-7 | 31 ( 6.24%) | 12 (13.48%) |  |
| TMA-8 | 38 ( 7.65%) | 4 ( 4.49%) |  |
| TMA-9 | 38 ( 7.65%) | 7 ( 7.87%) |  |
| TMA-10 | 38 ( 7.65%) | 3 ( 3.37%) |  |
| TMA-11 | 40 ( 8.05%) | 3 ( 3.37%) |  |
| TMA-12 | 36 ( 7.24%) | 9 (10.11%) |  |
| TMA-13 | 35 ( 7.04%) | 7 ( 7.87%) |  |
| TMA-14 | 38 ( 7.65%) | 6 ( 6.74%) |  |
| Follow-up, y, median<br>(range) | 12.9 (0.04-27.9) | 12.5 (0.29-19.9) | 0.1558 |
| Overall survival, y,<br>median (95% CI) | 12.8 (11.6, 16.5) | 15.4 (11.8, NE) | 0.1061 |
| Disease-free survival, y,<br>median (95% CI)** | 11.8 (9.9, 13.7) | 15.4 (11.2, NE) | 0.0352 |

\*from t-test, Chi-square, Fisher's Exact Test, or Log-Rank Test
\*\*M1 patients excluded from analysis
NE=Not Estimable

**Table 2.** Bivariate analysis of PDK4 H-score with other tumor markers

| Variable | PDK4 H-score Median (25<sup>th</sup> percentile, 75<sup>th</sup> percentile) |  |  |  |  |  | p-value* |
| --- | --- | --- | --- | --- | --- | --- | --- |
|  | n | values | n | Values | n | values |  |
|  | Negative |  | Positive |  |  |  |  |
| ER | 107 | 41.9 (22.7, 66.4) | 364 | 36.5 (17.7, 61.6) |  |  | 0.1001 |
| PR | 158 | 41.5 (19.4, 65.1) | 295 | 37.2 (18.5, 62.3) |  |  | 0.2937 |
| HER-2/neu | 228 | 32.6 (14.6, 54.6) | 47 | 38.7 (18.4, 61.9) |  |  | 0.2218 |
| ER+/PR+/HER- | 73 | 41.4 (18.9, 63.9) | 196 | 30.6 (14.4, 51.5) |  |  | 0.0114 |
| Nodal Status | 295 | 39.1 (20.0, 61.9) | 165 | 37.0 (17.0, 61.8) |  |  | 0.3870 |
|  | Grade 1 |  | Grade 2 |  | Grade 3 |  |  |
| Tumor Grade | 131 | 35.7 (17.2, 58.5) | 211 | 39.1 (20.0, 60.9) | 127 | 46.3 (19.9, 70.2) | 0.1399 |
|  | T0/1 |  | T2 |  | T3/4 |  |  |
| Tumor Stage | 270 | 36.6 (19.5, 63.1) | 181 | 40.6 (19.4, 61.9) | 40 | 40.7 (19.5, 60.9) | 0.9795 |

\*from Wilcoxon Rank Sum test for Hormone Receptor Status and Kruskal-Wallis test for Tumor Grade and Tumor Stage

**Table 3a.** Univariate analysis of other tumor markers for Overall Survival

| **Variable** | **n** | **Comparison\*** | **HR (95% CI)** | **p-value\*\*** |
| --- | --- | --- | --- | --- |
| ER | 556 | ER- vs ER+ | 1.05 (0.78, 1.41) | 0.7643 |
| PR | 537 | PR- vs PR+ | 1.40 (1.08, 1.82) | 0.0112 |
| HER-2/neu | 326 | HER-2/neu- vs HER-2/neu+ | 0.61 (0.40, 0.94) | 0.0269 |
| ER+/PR+/HER- | 320 | No vs Yes | 1.54 (1.08, 1.82) | 0.0420 |
| Tumor Grade | 550 | Grade 1 vs Grade 2 | 0.90 (0.66, 1.23) | <0.0001 |
|  |  | Grade 1 vs Grade 3 | 0.44 (0.32, 0.61) |  |
|  |  | Grade 2 vs Grade 3 | 0.49 (0.37, 0.66) |  |
| T Stage | 580 | T0/1 vs T2 | 0.69 (0.54, 0.89) | <0.0001 |
|  |  | T0/1 vs T3/4 | 0.31 (0.22, 0.45) |  |
|  |  | T2 vs T3/4 | 0.45 (0.31, 0.66) |  |
| N Stage | 543 | N+ vs N0 | 1.86 (1.44, 2.39) | <0.0001 |

\*referent group listed second
\*\*from Wald Chi-square test

**Table 3b.** Univariate analysis of other tumor markers for Disease Free Survival

| **Variable** | **n** | **Comparison\*** | **HR (95% CI)** | **p-value\*\*** |
| --- | --- | --- | --- | --- |
| ER | 538 | ER- vs ER+ | 1.07 (0.80, 1.44) | 0.6395 |
| PR | 520 | PR- vs PR+ | 1.31 (1.01, 1.69) | 0.0399 |
| HER-2/neu | 315 | HER-2/neu- vs HER-2/neu+ | 0.71 (0.46, 1.10) | 0.1266 |
| ER+/PR+/HER- | 309 | No vs Yes | 1.24 (0.85, 1.82) | 0.2735 |
| Tumor Grade | 534 | Grade 1 vs Grade 2 | 0.85 (0.64, 1.14) | <0.0001 |
|  |  | Grade 1 vs Grade 3 | 0.45 (0.33, 0.62) |  |
|  |  | Grade 2 vs Grade 3 | 0.53 (0.39, 0.70) |  |
| T Stage | 562 | T0/1 vs T2 | 0.64 (0.50, 0.81) | <0.0001 |
|  |  | T0/1 vs T3/4 | 0.36 (0.25, 0.54) |  |
|  |  | T2 vs T3/4 | 0.57 (0.38, 0.86) |  |
| N Stage | 527 | N+ vs N0 | 1.79 (1.40, 2.29) | <0.0001 |

\*referent group listed second
\*\*from Wald Chi-square test

**Table 4aa.** Multivariable analysis for Overall Survival including HER-2 (N=274)

| Variable | Comparison* | HR | Lower<br>95% CI | Upper<br>95% CI | p-value** |
| --- | --- | --- | --- | --- | --- |
| PR | PR- vs PR+ | 1.690 | 1.094 | 2.611 | 0.0006 |
|  | PR- vs Unknown | 0.394 | 0.182 | 0.852 |  |
|  | PR+ vs Unknown | 0.233 | 0.106 | 0.511 |  |
| HER-2/neu | HER-2/neu- vs HER-2/neu+ | 1.056 | 0.623 | 1.792 | 0.8393 |
| Tumor Grade | Grade 1 vs Grade 2 | 0.996 | 0.588 | 1.689 | 0.0594 |
|  | Grade 1 vs Grade 3 | 0.727 | 0.409 | 1.291 |  |
|  | Grade 1 vs Unknown | 0.382 | 0.168 | 0.868 |  |
|  | Grade 2 vs Grade 3 | 0.730 | 0.447 | 1.191 |  |
|  | Grade 2 vs Unknown | 0.384 | 0.179 | 0.823 |  |
|  | Grade 3 vs Unknown | 0.526 | 0.233 | 1.186 |  |
| T Stage | T0/1 vs T2 | 0.807 | 0.530 | 1.228 | 0.7074 |
|  | T0/1 vs T3/4 | 0.867 | 0.438 | 1.718 |  |
|  | T0/1 vs TX | 0.566 | 0.128 | 2.497 |  |
|  | T2 vs T3/4 | 1.075 | 0.547 | 2.112 |  |
|  | T2 vs TX | 0.702 | 0.159 | 3.101 |  |
|  | T3/4 vs TX | 0.653 | 0.133 | 3.214 |  |
| N Stage | N+ vs N0 | 1.916 | 1.262 | 2.909 | <0.0001 |
|  | N+ vs NX | 0.476 | 0.231 | 0.981 |  |
|  | N0 vs NX | 0.249 | 0.121 | 0.510 |  |
| PDK4 H-score<br>Category | High vs Low | 1.473 | 0.960 | 2.258 | 0.0761 |

\*referent group listed second
\*\*from Wald Chi-square test

**Table 4ab.**
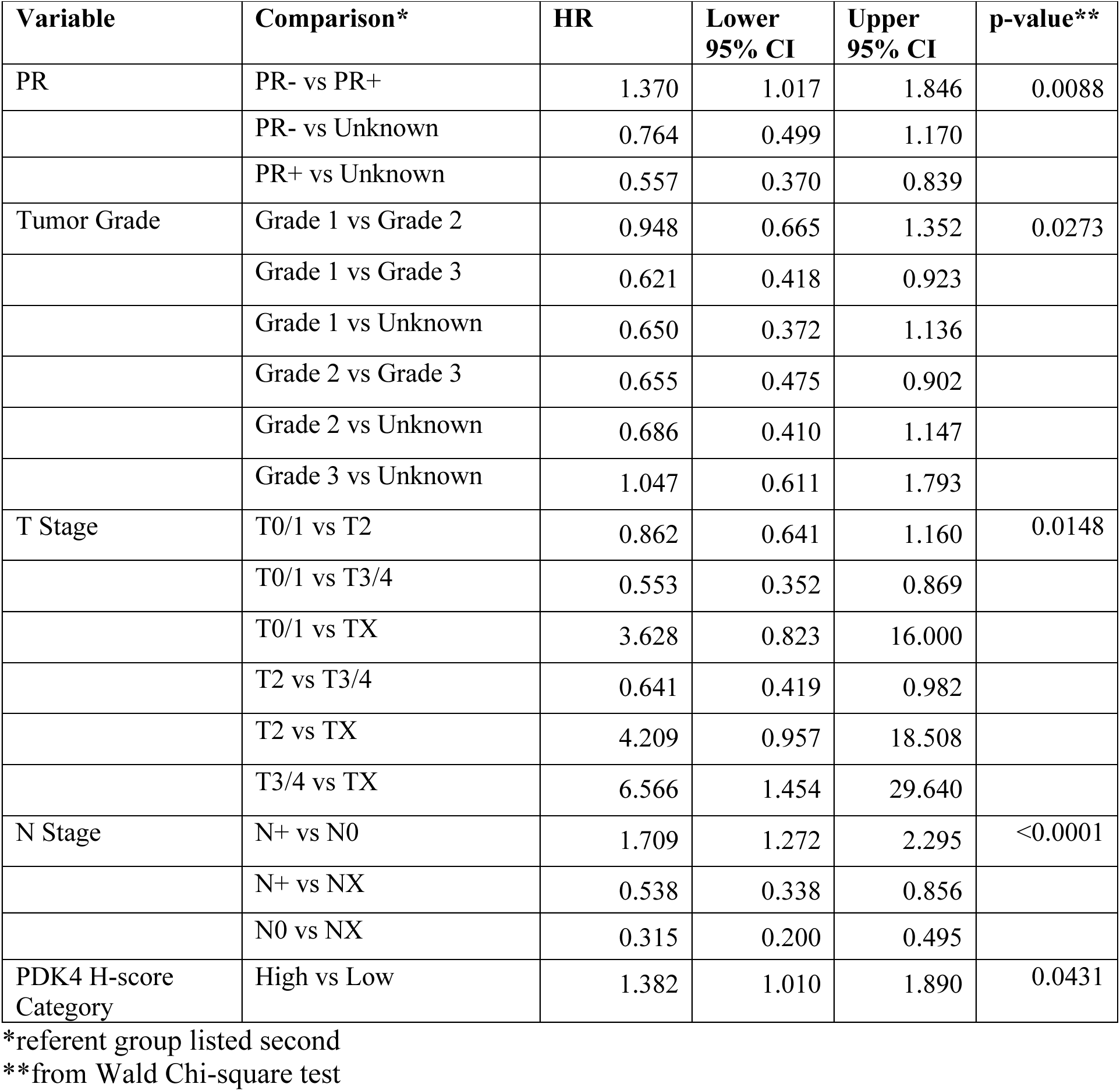
Multivariable analysis for Overall Survival not including HER-2 (N=494)

**Table 4b.** Multivariable analysis for Disease Free Survival (n=477)

| **Variable** | **Comparison\*** | **HR** | **Lower**<br>**95% CI** | **Upper**<br>**95% CI** | **p-value\*\*** |
| --- | --- | --- | --- | --- | --- |
| PR | PR- vs PR+ | 1.260 | 0.940 | 1.689 | 0.0401 |
|  | PR- vs Unknown | 0.771 | 0.503 | 1.183 |  |
|  | PR+ vs Unknown | 0.612 | 0.408 | 0.919 |  |
| Tumor Grade | Grade 1 vs Grade 2 | 0.871 | 0.623 | 1.217 | 0.0356 |
|  | Grade 1 vs Grade 3 | 0.586 | 0.398 | 0.864 |  |
|  | Grade 1 vs Unknown | 0.754 | 0.433 | 1.313 |  |
|  | Grade 2 vs Grade 3 | 0.674 | 0.489 | 0.927 |  |
|  | Grade 2 vs Unknown | 0.865 | 0.515 | 1.455 |  |
|  | Grade 3 vs Unknown | 1.285 | 0.739 | 2.234 |  |
| T Stage | T0/1 vs T2 | 0.735 | 0.555 | 0.974 | 0.0351 |
|  | T0/1 vs T3/4 | 0.596 | 0.372 | 0.955 |  |
|  | T0/1 vs TX | 1.877 | 0.534 | 6.594 |  |
|  | T2 vs T3/4 | 0.812 | 0.516 | 1.277 |  |
|  | T2 vs TX | 2.554 | 0.730 | 8.933 |  |
|  | T3/4 vs TX | 3.146 | 0.859 | 11.520 |  |
| N Stage | N+ vs N0 | 1.550 | 1.164 | 2.063 | 0.0001 |
|  | N+ vs NX | 0.640 | 0.398 | 1.030 |  |
|  | N0 vs NX | 0.413 | 0.259 | 0.659 |  |
| PDK4 H-score<br>Category | High vs Low | 0.879 | 0.680 | 1.136 | 0.3241 |

\*referent group listed second
\*\*from Wald Chi-square test

**Table 5a:** Overall Survival - PROC PHREG with H score as dichotomous variable in multivariable models

| **Obs** | **Group** | **H-Score**<br>**Category**<br>**Parameter\*** | **p-value** | **Point**<br>**Estimate** | **Lower**<br>**95%**<br>**Wald**<br>**Confidenc**<br>**e Limit** | **Upper**<br>**95%**<br>**Wald**<br>**Confidenc**<br>**e Limit** |
| --- | --- | --- | --- | --- | --- | --- |
| **1** | OS | High vs Low | 0.0431 | 1.382 | 1.010 | 1.890 |
| **2** | OS for ER+ | High vs Low | 0.0131 | 1.600 | 1.104 | 2.319 |
| **3** | OS for ER- | High vs Low | 0.5535 | 1.239 | 0.610 | 2.514 |
| **4** | OS for Endocrine<br>Therapy=Yes | High vs Low | 0.0847 | 1.431 | 0.952 | 2.152 |
| **5** | OS for Endocrine Therapy=No | High vs Low | 0.0976 | 1.608 | 0.917 | 2.822 |
| **6** | OS for ER+ and ET=Yes | High vs Low | 0.0300 | 1.614 | 1.048 | 2.488 |
| **7** | OS for ER+ and ET=No | High vs Low | 0.1473 | 1.956 | 0.789 | 4.847 |
| **8** | OS for ER- and ET=Yes | High vs Low | 0.9707 | 0.961 | 0.117 | 7.887 |
| **9** | OS for ER- and ET=No | High vs Low | 0.2016 | 1.710 | 0.751 | 3.893 |
| **10** | OS for ER+/PR+/HER2-=Yes | High vs Low | 0.1131 | 1.510 | 0.907 | 2.513 |
| **11** | OS for ER+/PR+/HER2-=No | High vs Low | 0.7579 | 1.146 | 0.483 | 2.717 |

\*referent group listed second

**Table 5b:** Disease Free Survival - PROC PHREG with H score as dichotomous variable in multivariable models

| **Obs** | **Group** | **H-Score**<br>**Category**<br>**Parameter\*** | **p-value** | **Point**<br>**Estimate** | **Lower**<br>**95%**<br>**Wald**<br>**Confidence**<br>**Limit** | **Upper**<br>**95%**<br>**Wald**<br>**Confidence**<br>**Limit** |
| --- | --- | --- | --- | --- | --- | --- |
| **1** | DFS | High vs Low | 0.3241 | 0.879 | 0.680 | 1.136 |
| **2** | DFS for ER+ | High vs Low | 0.5038 | 0.902 | 0.666 | 1.221 |
| **3** | DFS for ER- | High vs Low | 0.9336 | 0.975 | 0.537 | 1.772 |
| **4** | DFS for Endocrine<br>Therapy=Yes | High vs Low | 0.4082 | 1.151 | 0.824 | 1.608 |
| **5** | DFS for Endocrine<br>Therapy=No | High vs Low | 0.2109 | 0.754 | 0.484 | 1.174 |
| **6** | DFS for ER+ and ET=Yes | High vs Low | 0.3474 | 1.190 | 0.828 | 1.711 |
| **7** | DFS for ER+ and ET=No | High vs Low | 0.0448 | 0.474 | 0.228 | 0.983 |
| **8** | DFS for ER- and ET=Yes | High vs Low | 0.6785 | 1.355 | 0.322 | 5.710 |
| **9** | DFS for ER- and ET=No | High vs Low | 0.6392 | 1.179 | 0.592 | 2.350 |
| **10** | DFS for ER+/PR+/HER2-=Yes | High vs Low | 0.2106 | 0.750 | 0.478 | 1.177 |
| **11** | DFS for ER+/PR+/HER2-=No | High vs Low | 0.4555 | 1.393 | 0.584 | 3.323 |

\*referent group listed second

**Table 6a:** Overall Survival - PROC PHREG with H score as continuous variable in multivariable models

| **Obs** | **Group** | **Parameter** | **p-value** | **Point**<br>**Estimate** | **Lower**<br>**95%**<br>**Wald**<br>**Confidence**<br>**Limit** | **Upper**<br>**95%**<br>**Wald**<br>**Confidence**<br>**Limit** |
| --- | --- | --- | --- | --- | --- | --- |
| **1** | OS | score | 0.1082 | 1.003 | 0.999 | 1.007 |
| **2** | OS for ER+ | score | 0.0043 | 1.006 | 1.002 | 1.011 |
| **3** | OS for ER- | score | 0.4747 | 0.997 | 0.988 | 1.005 |
| **4** | OS for Endocrine<br>Therapy=Yes | score | 0.0008 | 1.009 | 1.004 | 1.014 |
| **5** | OS for Endocrine<br>Therapy=No | score | 0.9615 | 1.000 | 0.995 | 1.006 |
| **6** | OS for ER+ and ET=Yes | score | 0.0007 | 1.009 | 1.004 | 1.015 |
| **7** | OS for ER+ and ET=No | score | 0.2803 | 1.005 | 0.996 | 1.013 |
| **8** | OS for ER- and ET=Yes | score | 0.3364 | 1.019 | 0.981 | 1.058 |
| **9** | OS for ER- and ET=No | score | 0.3776 | 0.996 | 0.986 | 1.005 |
| **10** | OS for ER+/PR+/HER2-<br>=Yes | score | 0.0333 | 1.008 | 1.001 | 1.016 |
| **11** | OS for ER+/PR+/HER2-<br>=No | score | 0.5765 | 0.996 | 0.983 | 1.010 |

**Table 6b:** Disease Free Survival - PROC PHREG with H score as continuous variable in multivariable models

| **Obs** | **Group** | **Parameter** | **p-value** | **Point Estimate** | **Lower 95% Wald Confidence Limit** | **Upper 95% Wald Confidence Limit** |
| --- | --- | --- | --- | --- | --- | --- |
| **1** | DFS | score | 0.4423 | 1.001 | 0.998 | 1.005 |
| **2** | DFS for ER+ | score | 0.2105 | 1.003 | 0.998 | 1.007 |
| **3** | DFS for ER- | score | 0.6455 | 0.998 | 0.991 | 1.006 |
| **4** | DFS for Endocrine Therapy=Yes | score | 0.0103 | 1.007 | 1.002 | 1.012 |
| **5** | DFS for Endocrine Therapy=No | score | 0.7727 | 0.999 | 0.994 | 1.005 |
| **6** | DFS for ER+ and ET=Yes | score | 0.0237 | 1.007 | 1.001 | 1.012 |
| **7** | DFS for ER+ and ET=No | score | 0.8985 | 0.999 | 0.991 | 1.008 |
| **8** | DFS for ER- and ET=Yes | score | 0.2757 | 1.015 | 0.988 | 1.043 |
| **9** | DFS for ER- and ET=No | score | 0.7149 | 0.998 | 0.990 | 1.007 |
| **10** | DFS for ER+/PR+/HER2-=Yes | score | 0.6246 | 1.002 | 0.994 | 1.010 |
| **11** | DFS for ER+/PR+/HER2-=No | score | 0.7799 | 1.002 | 0.989 | 1.015 |

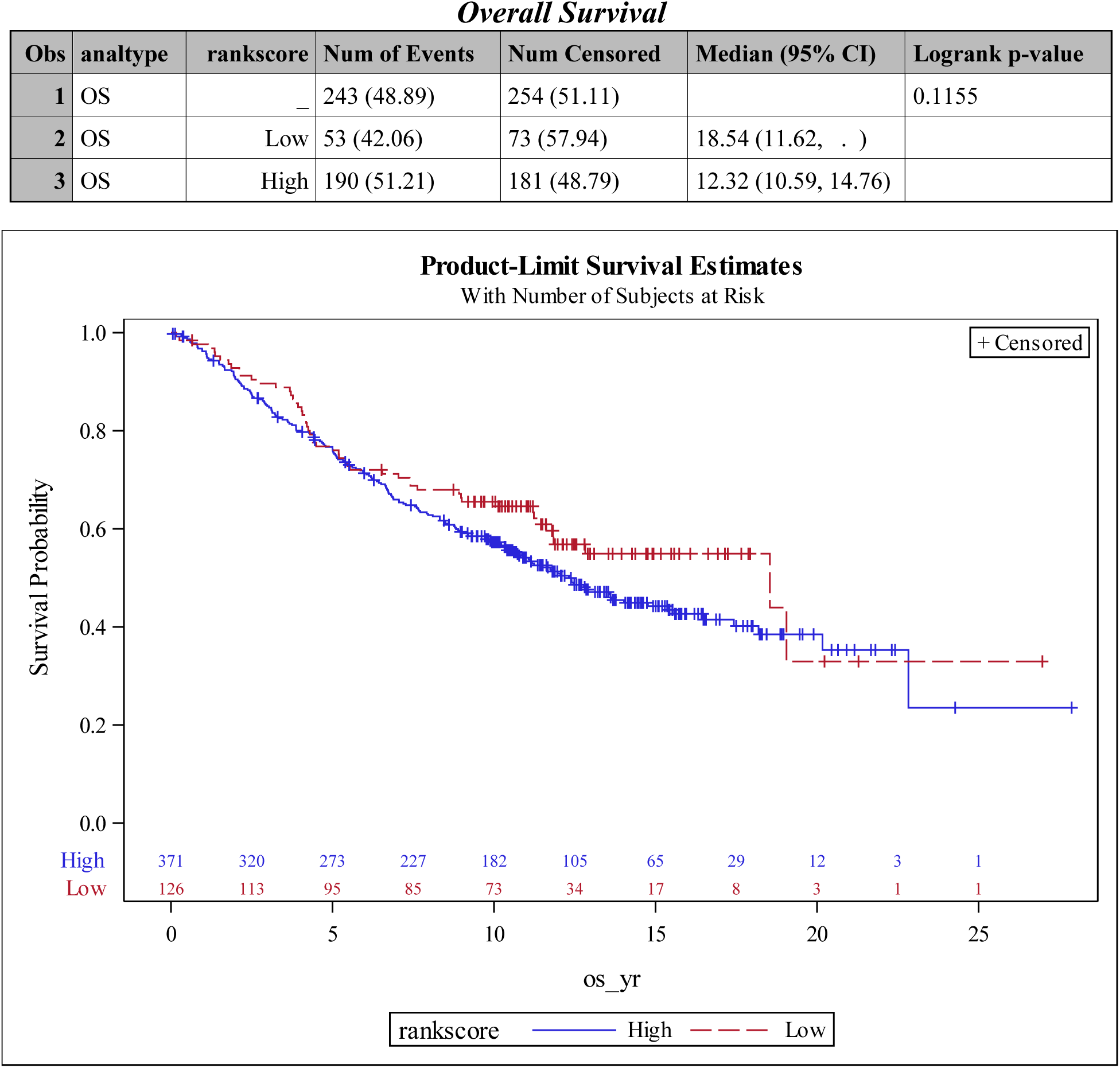
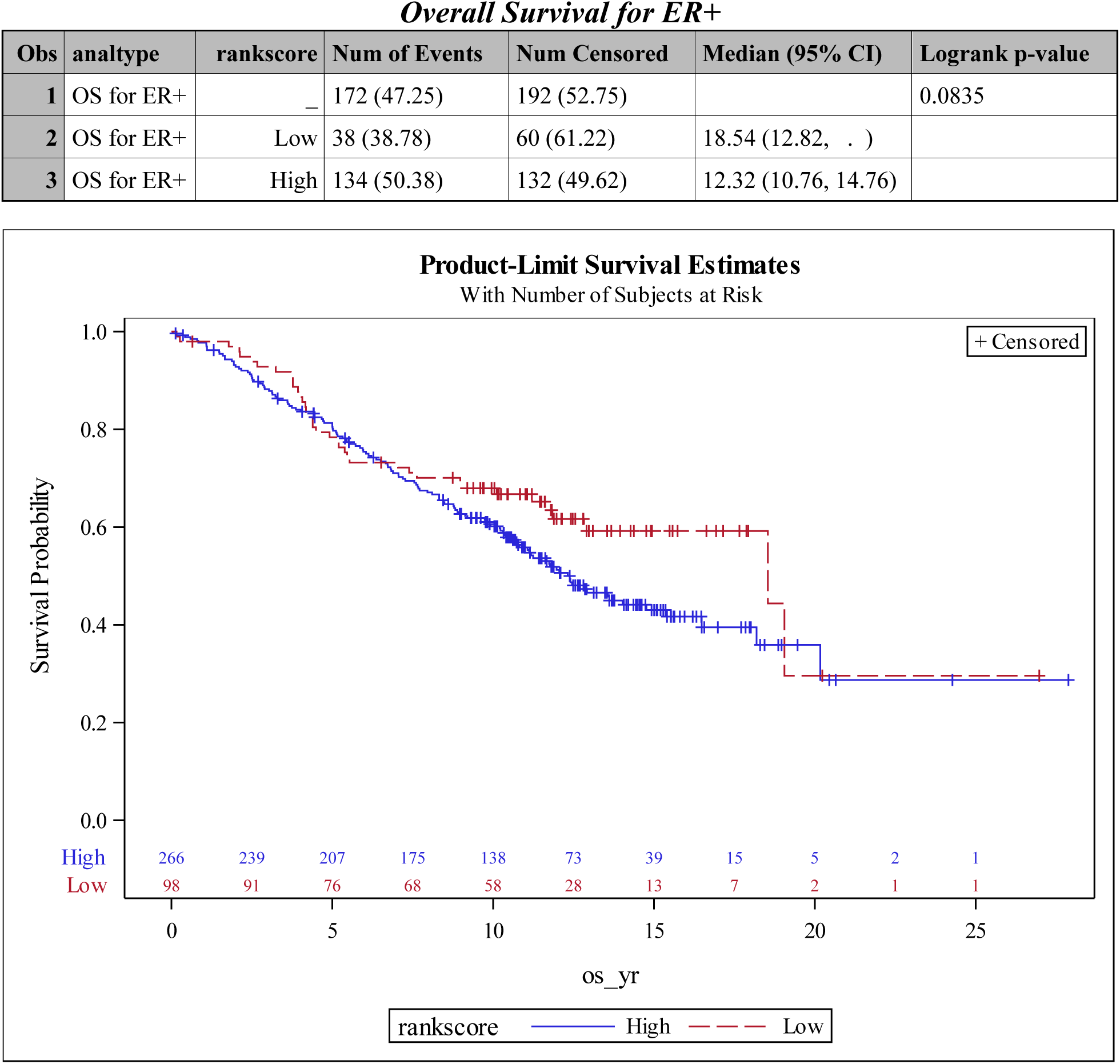
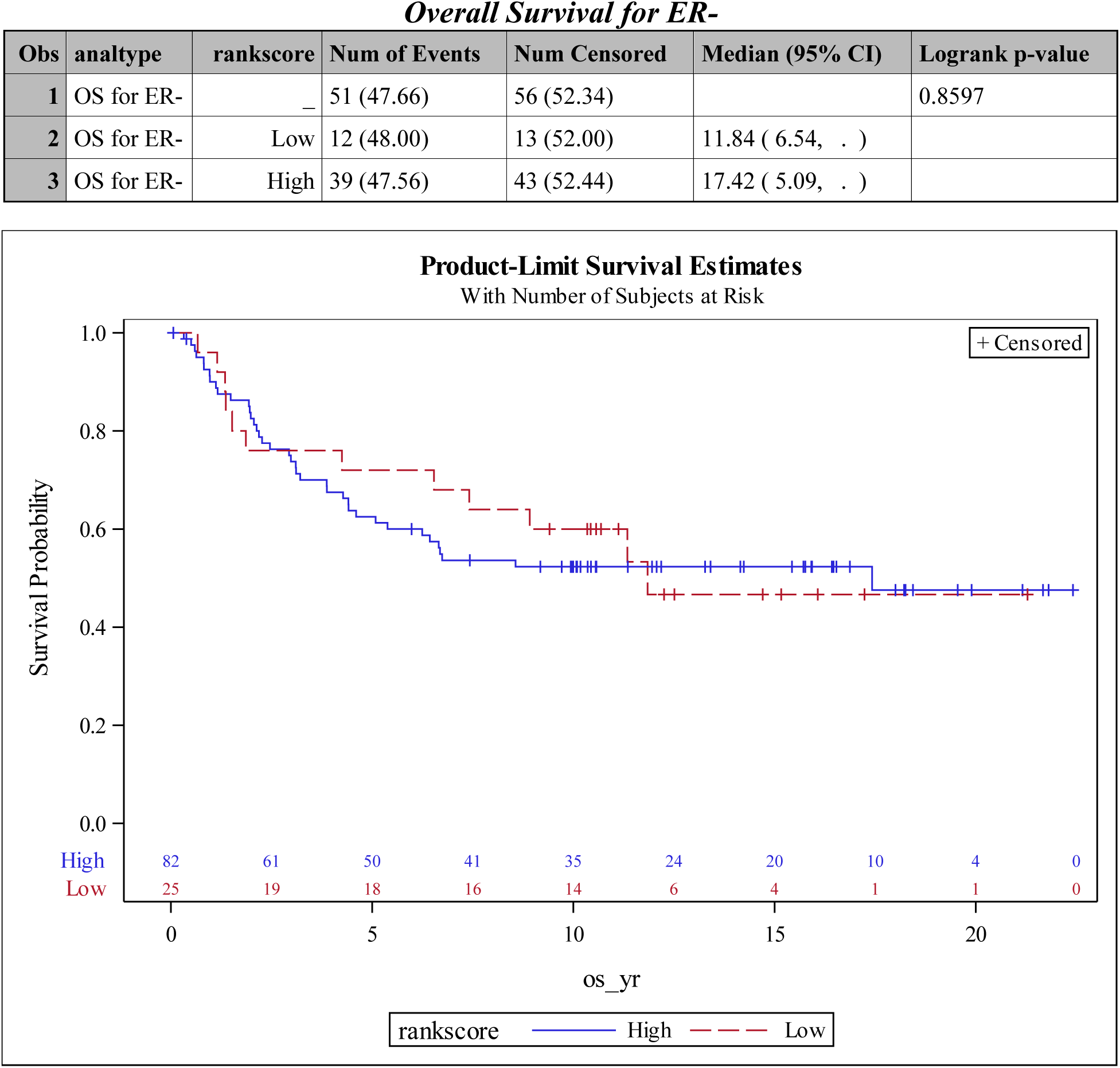
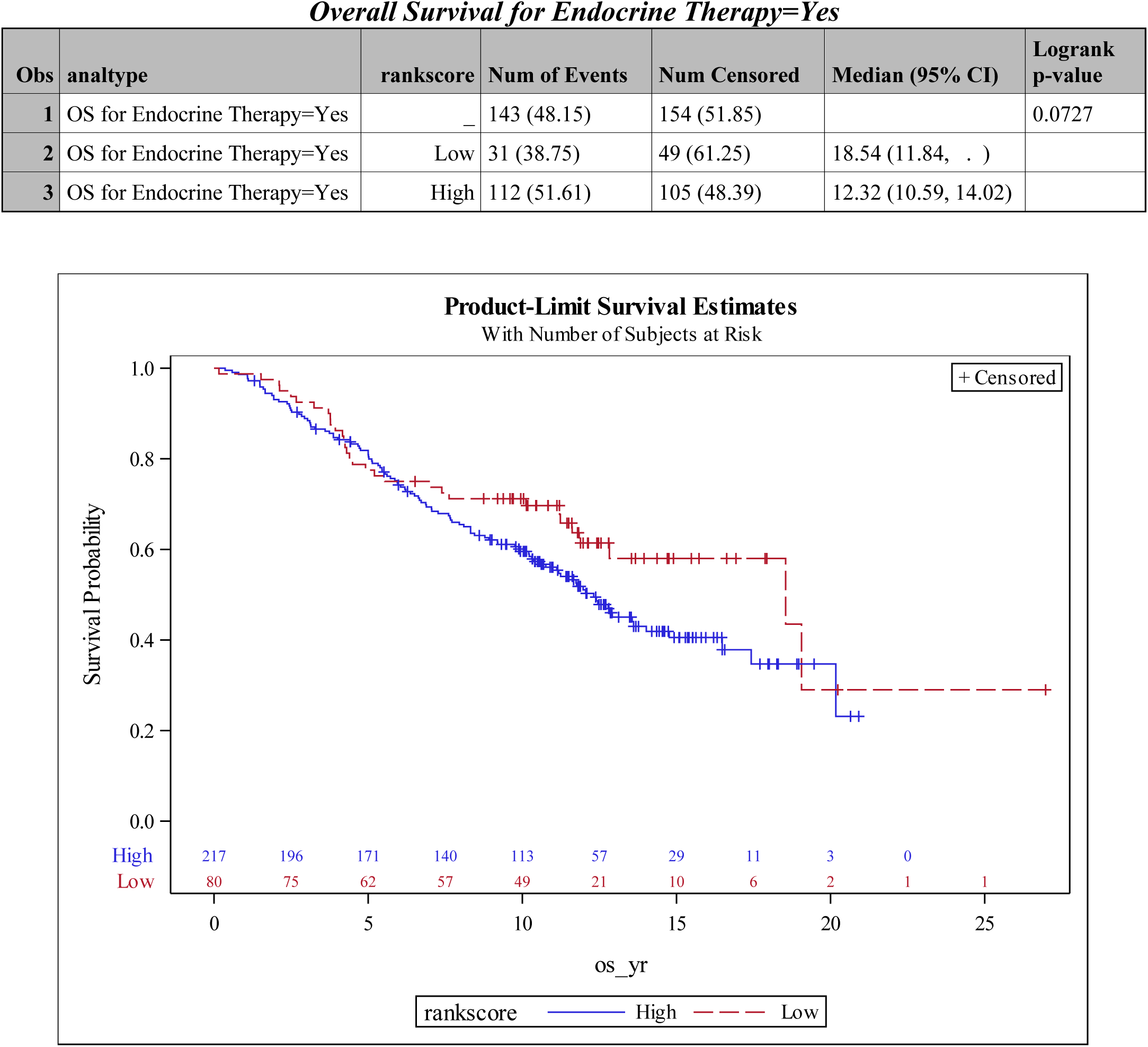
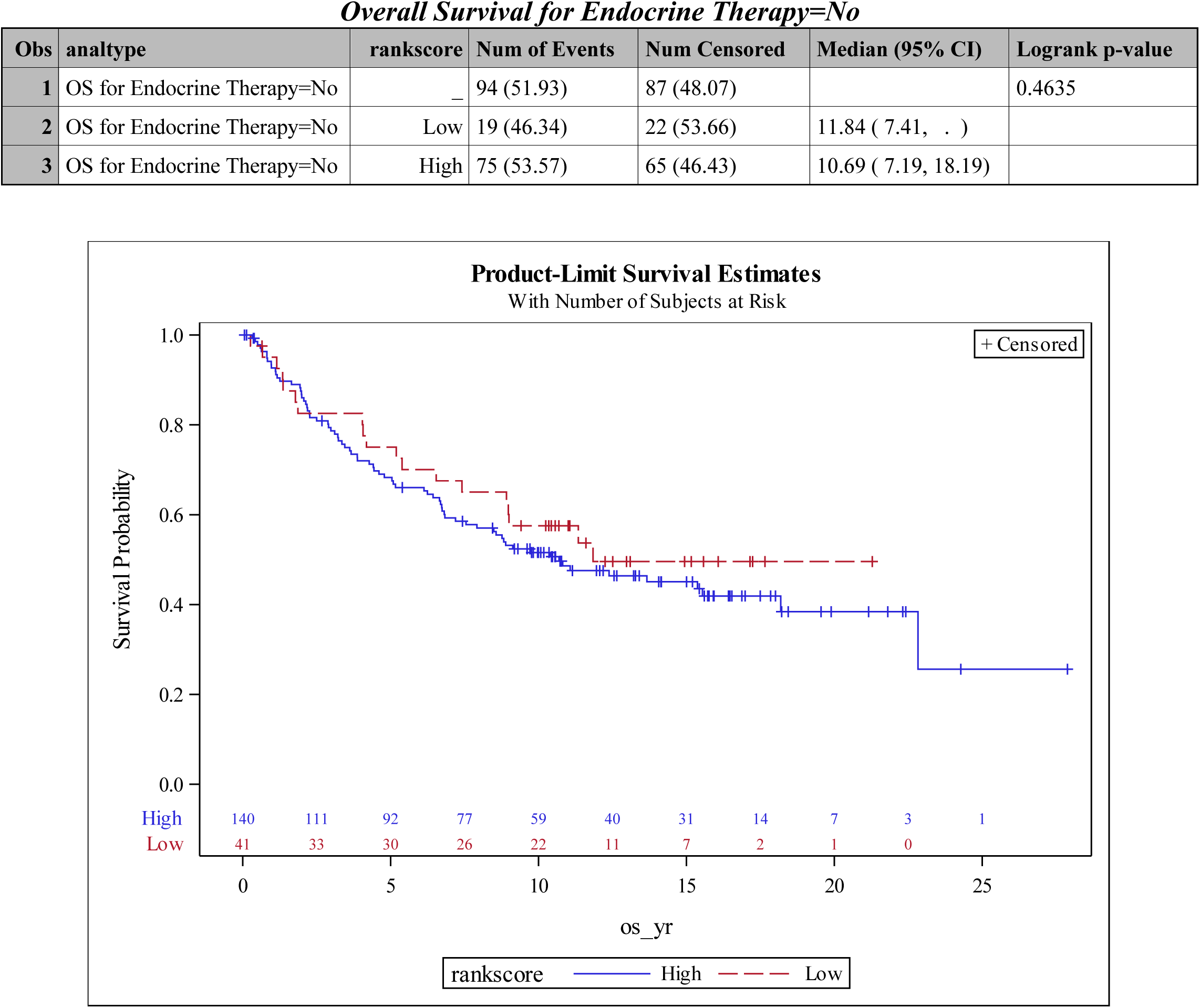
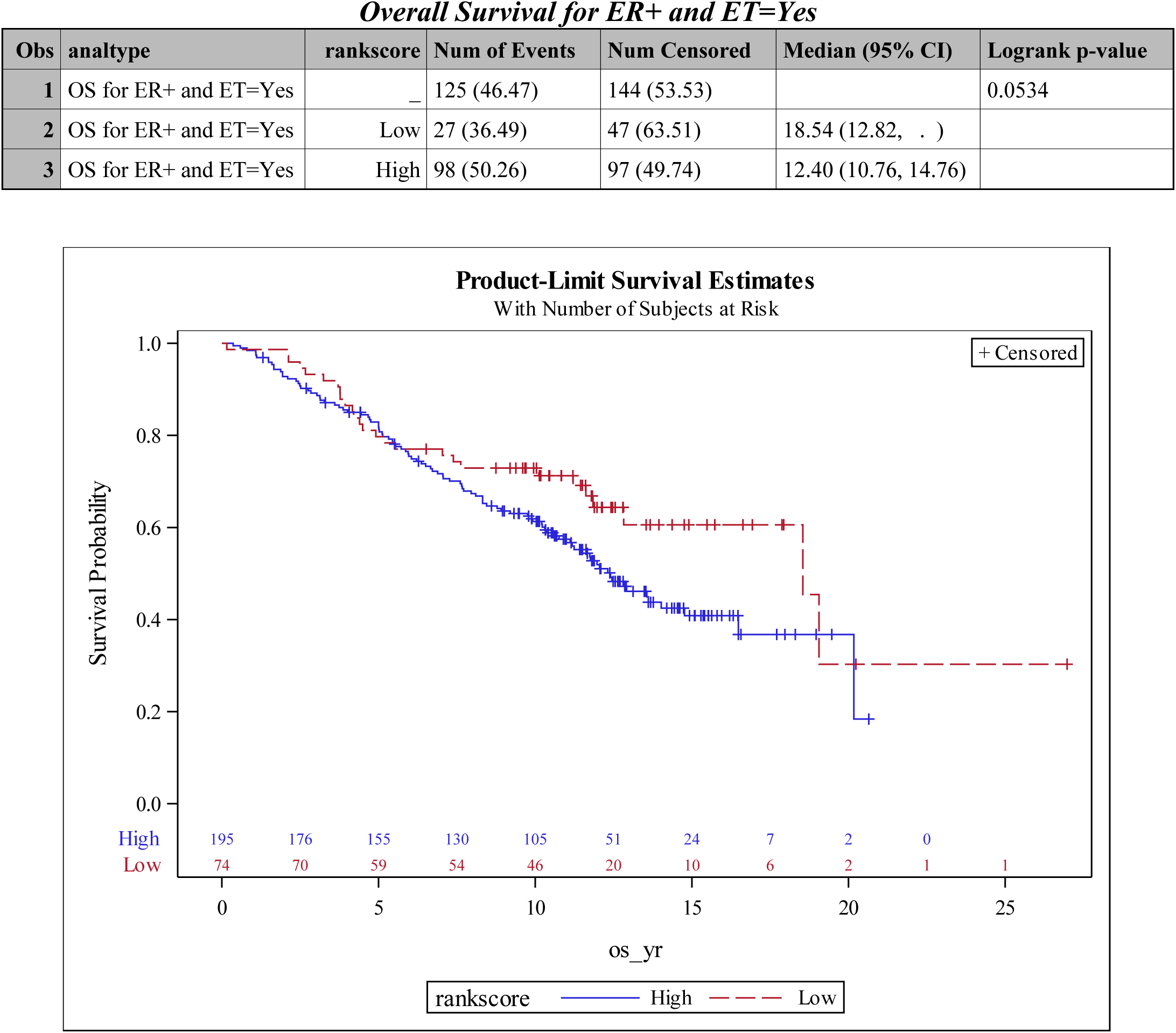
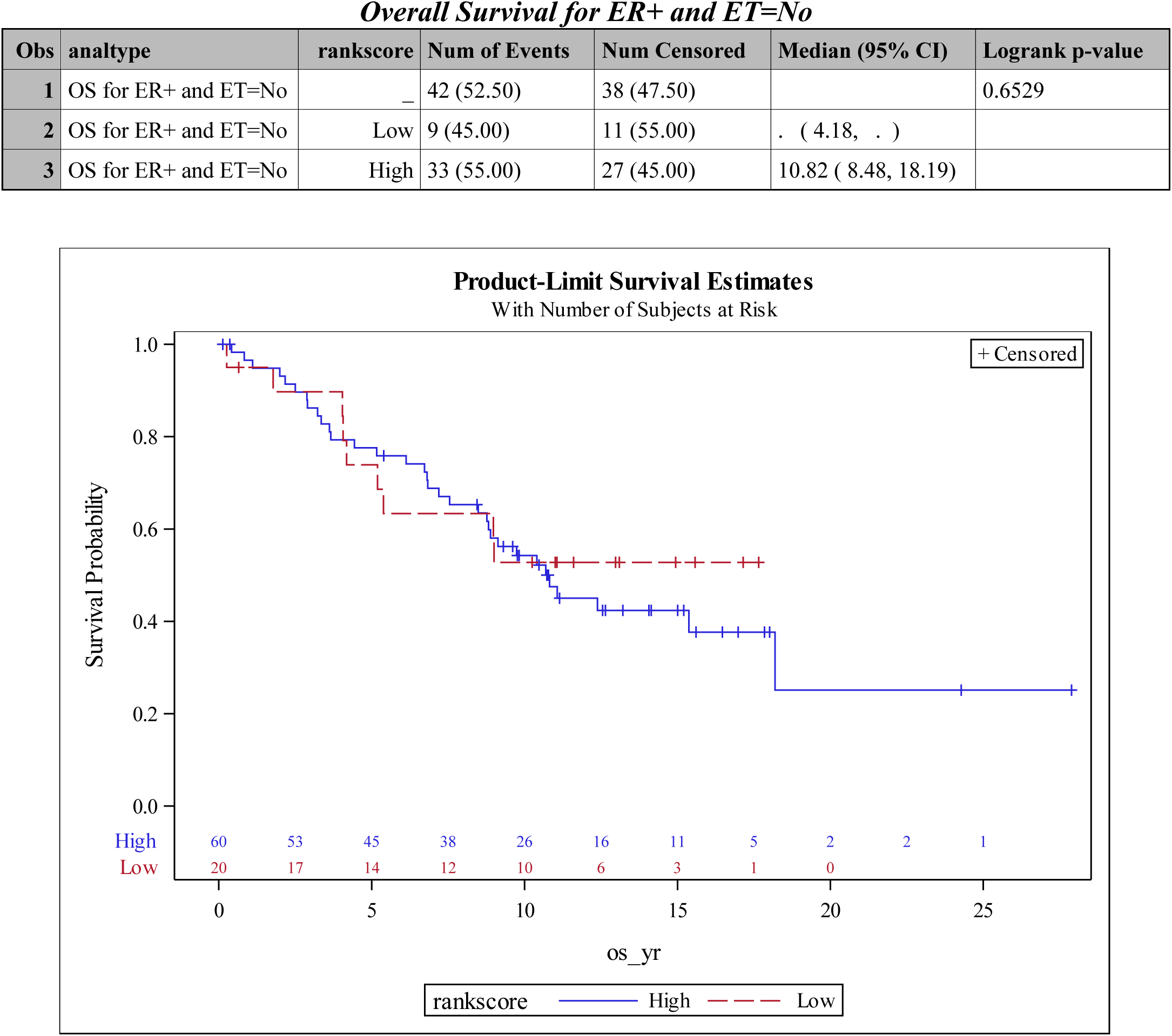
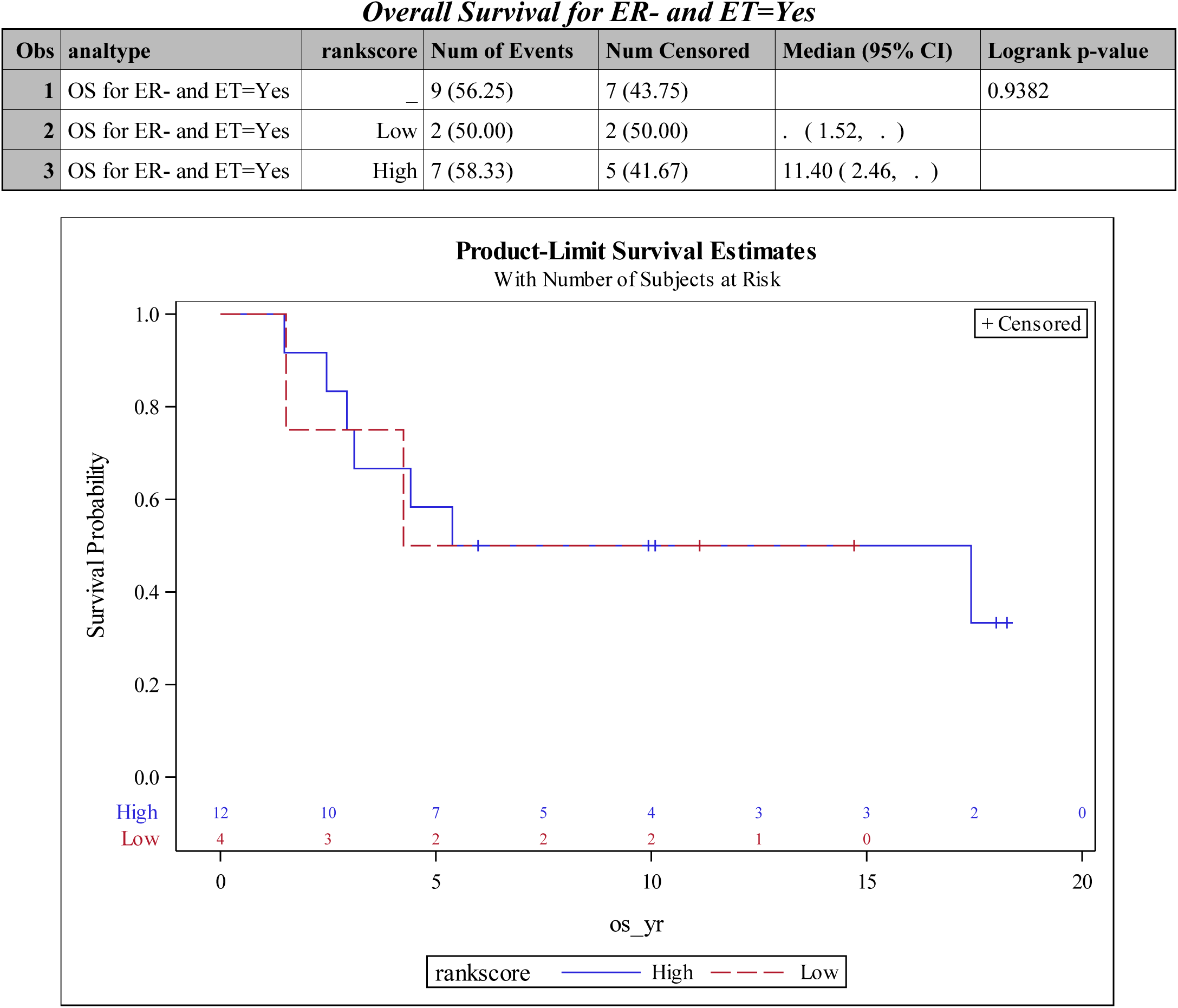
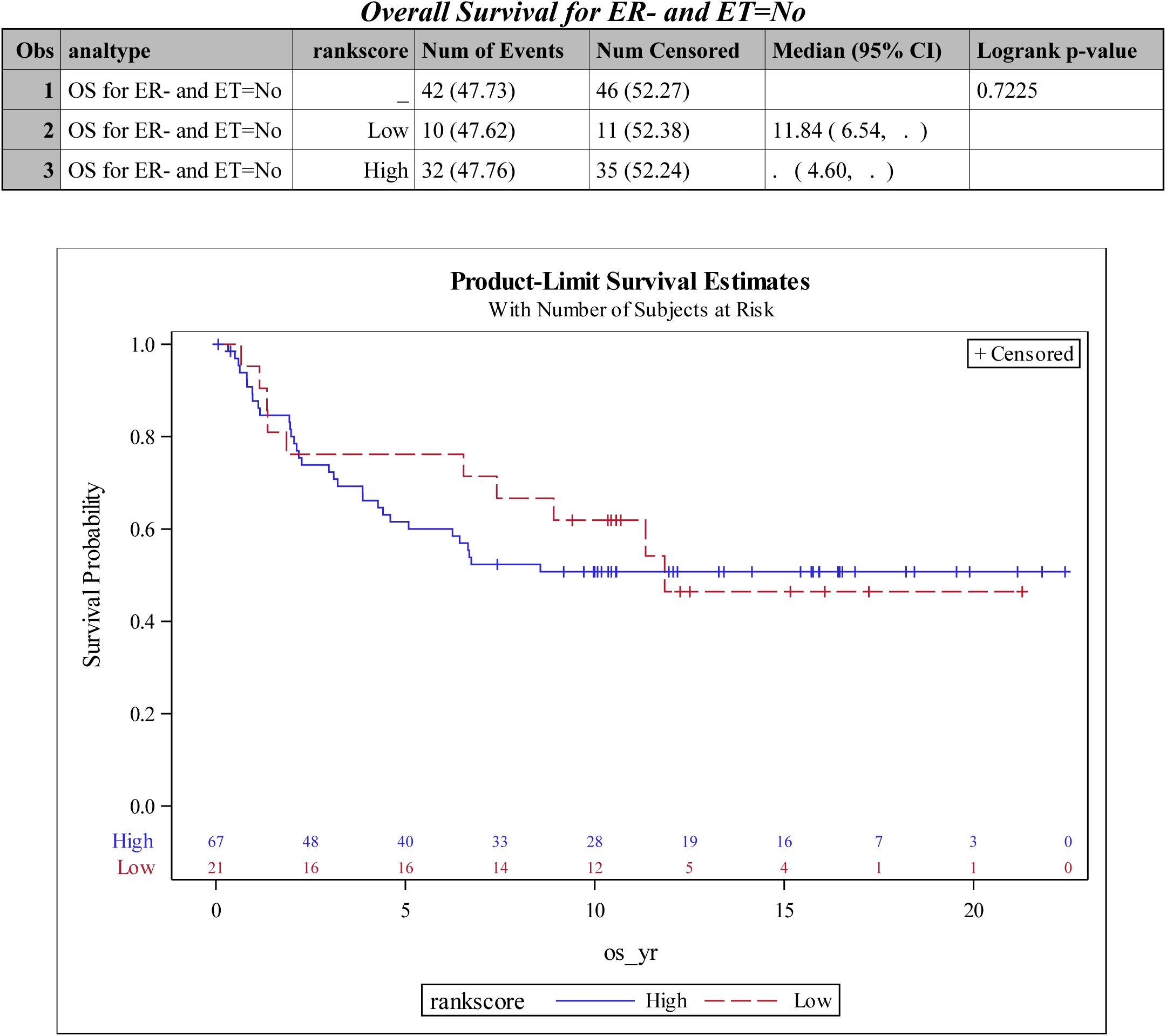
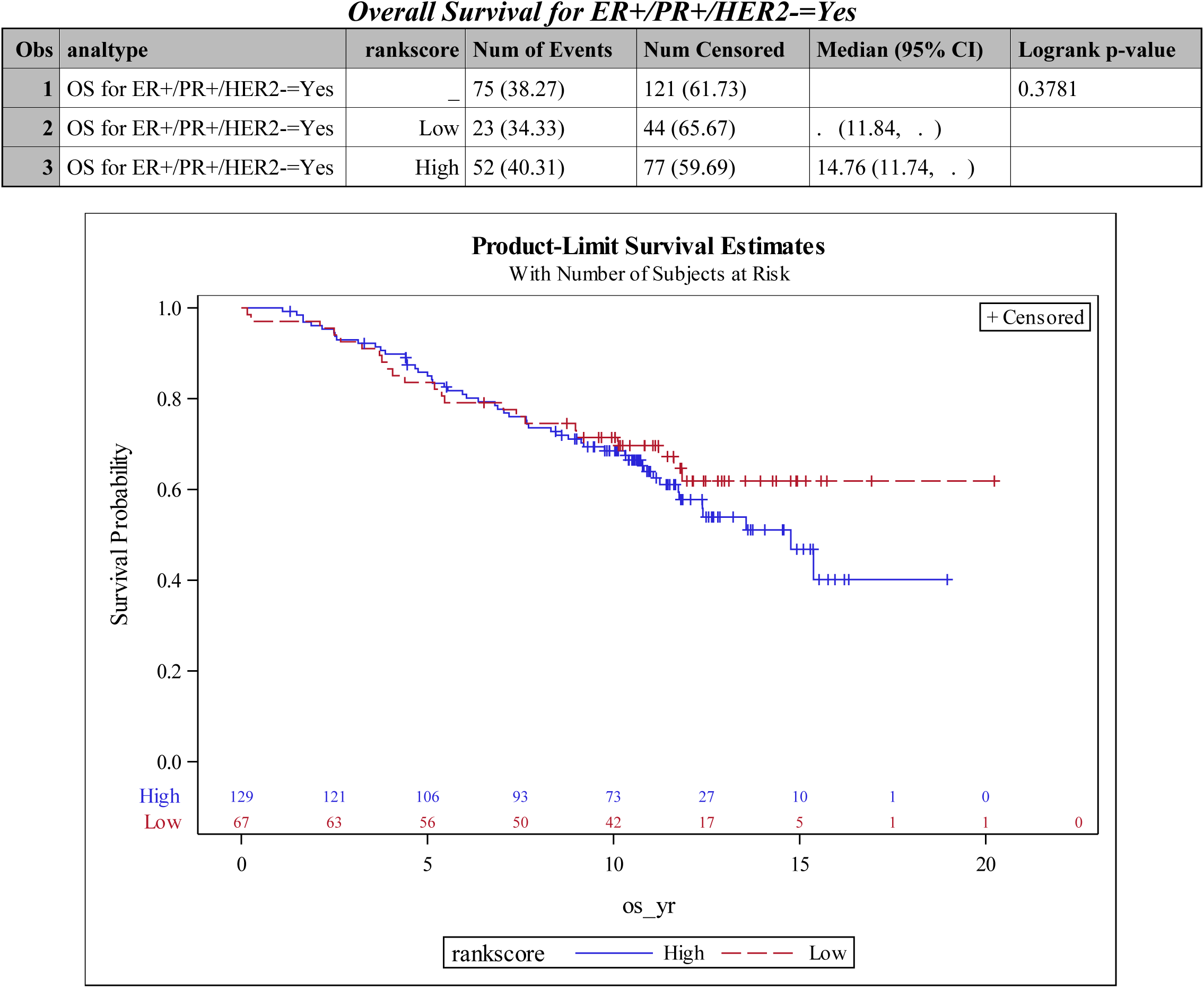
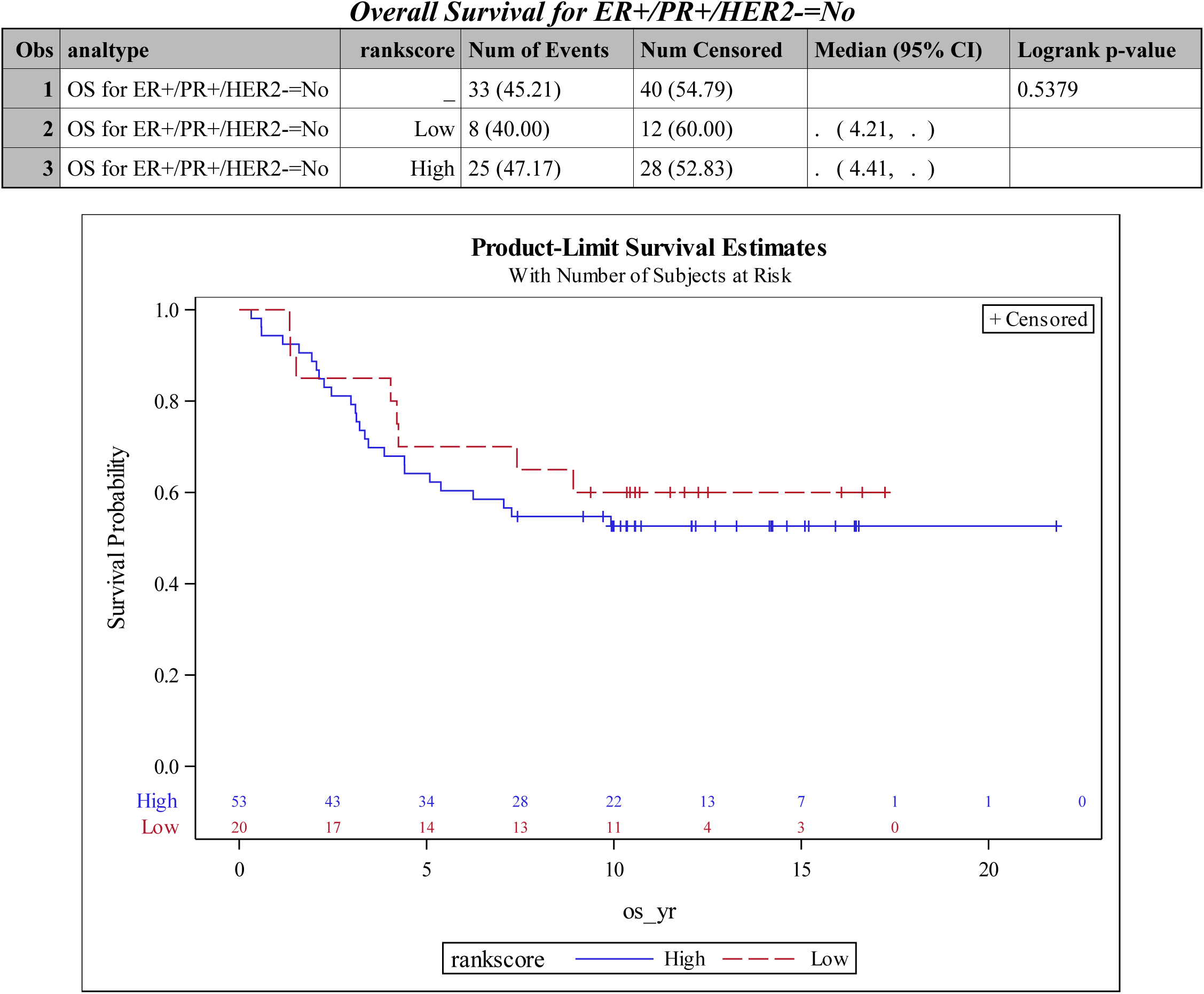
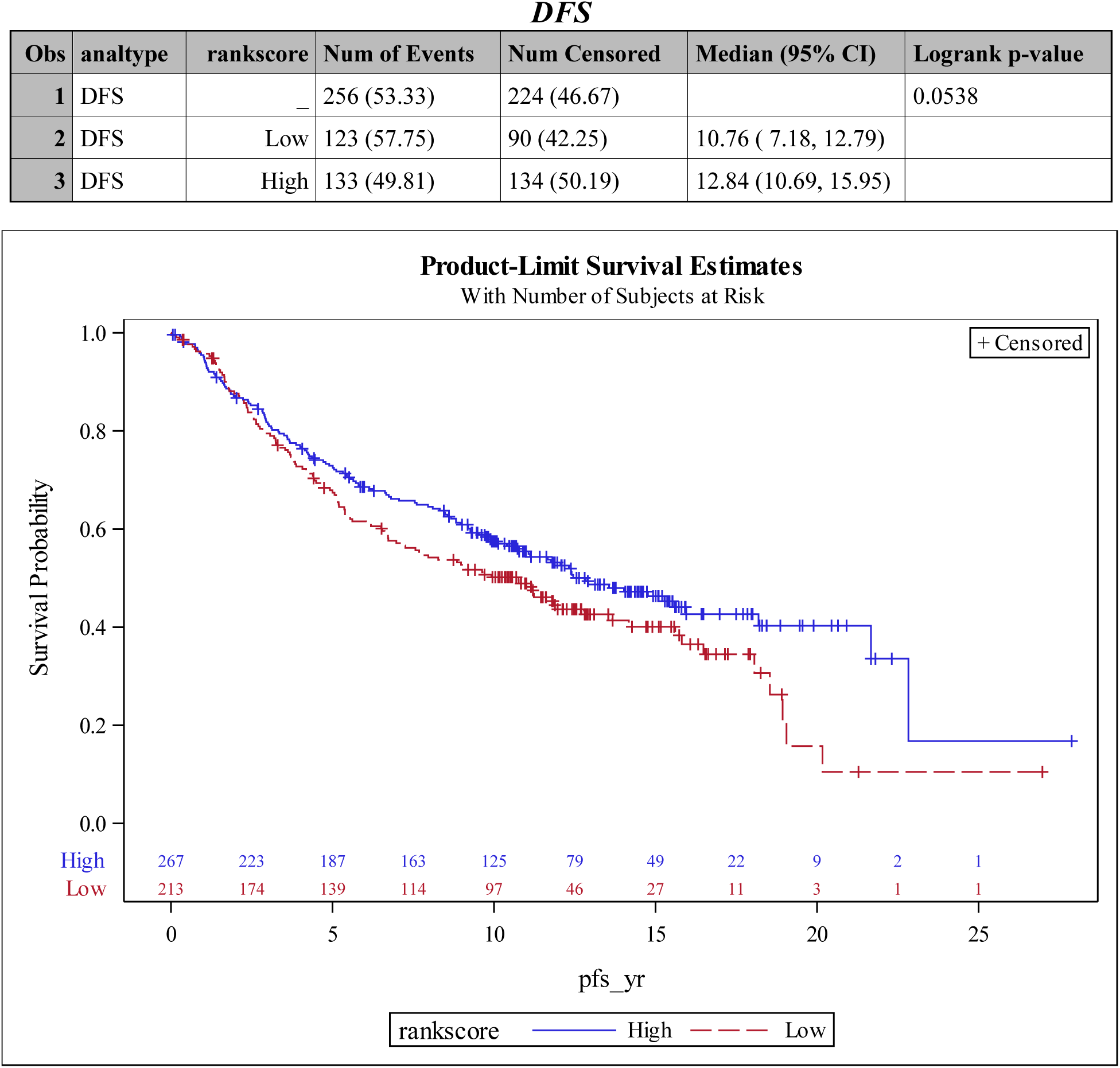
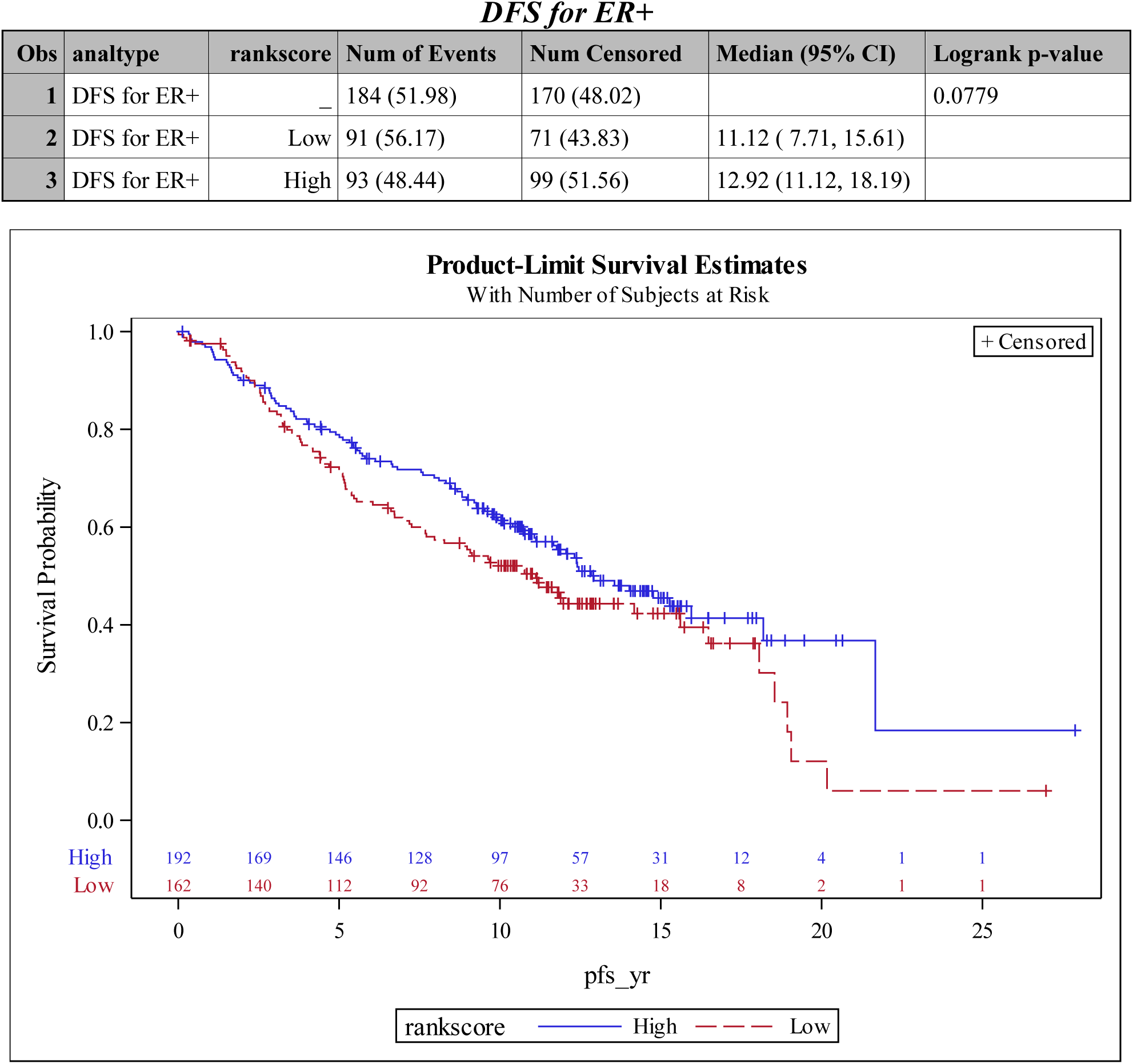
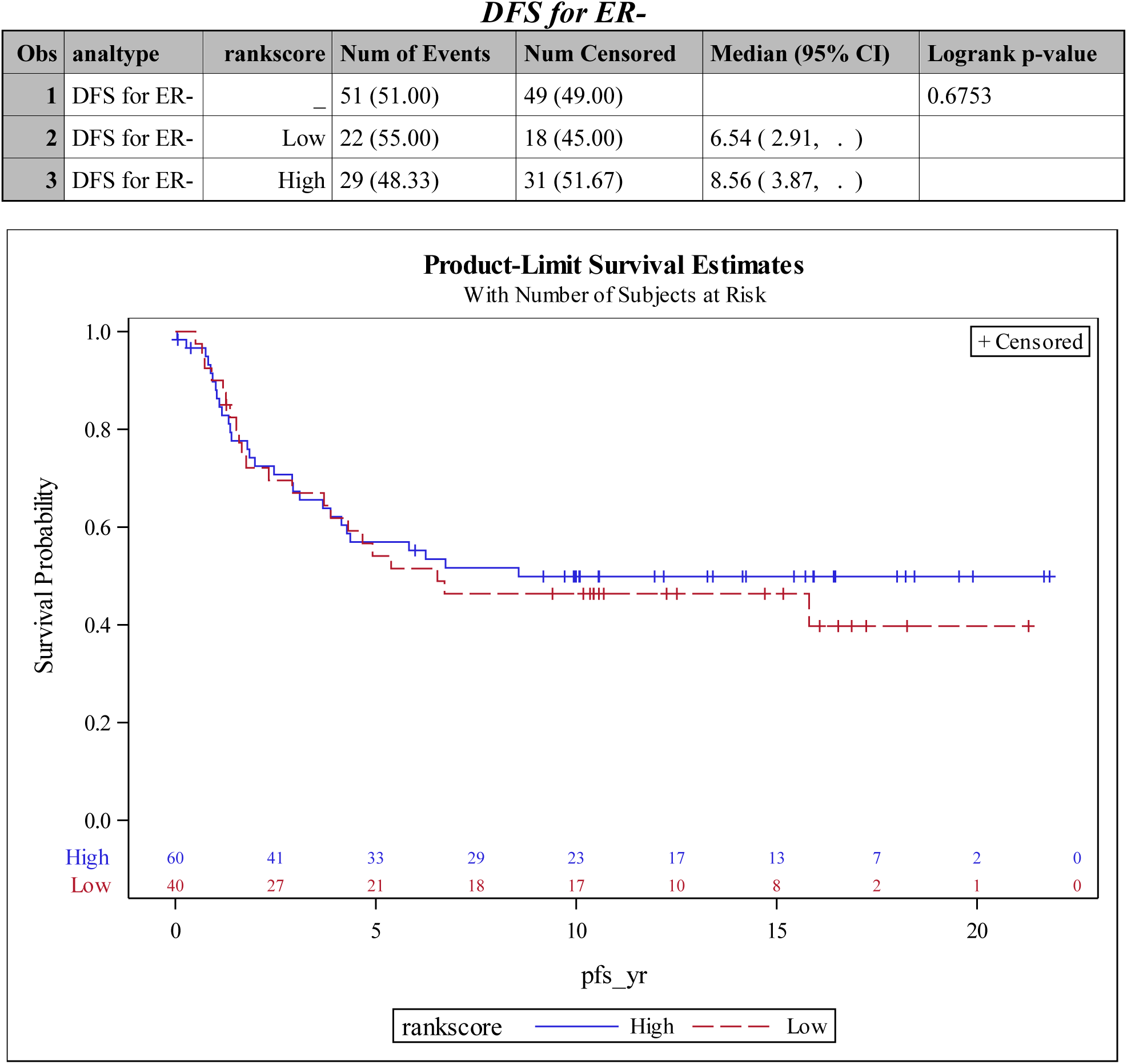
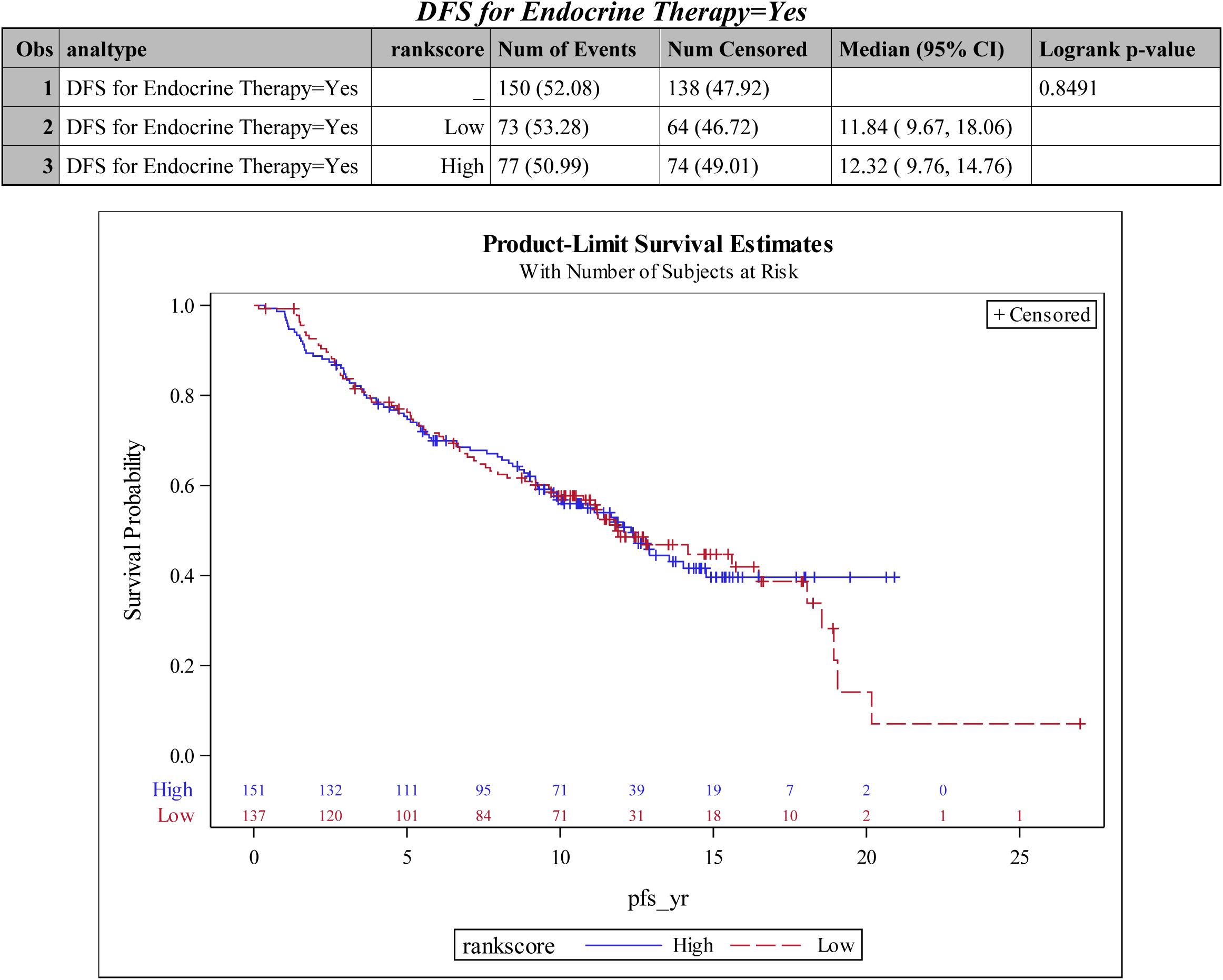
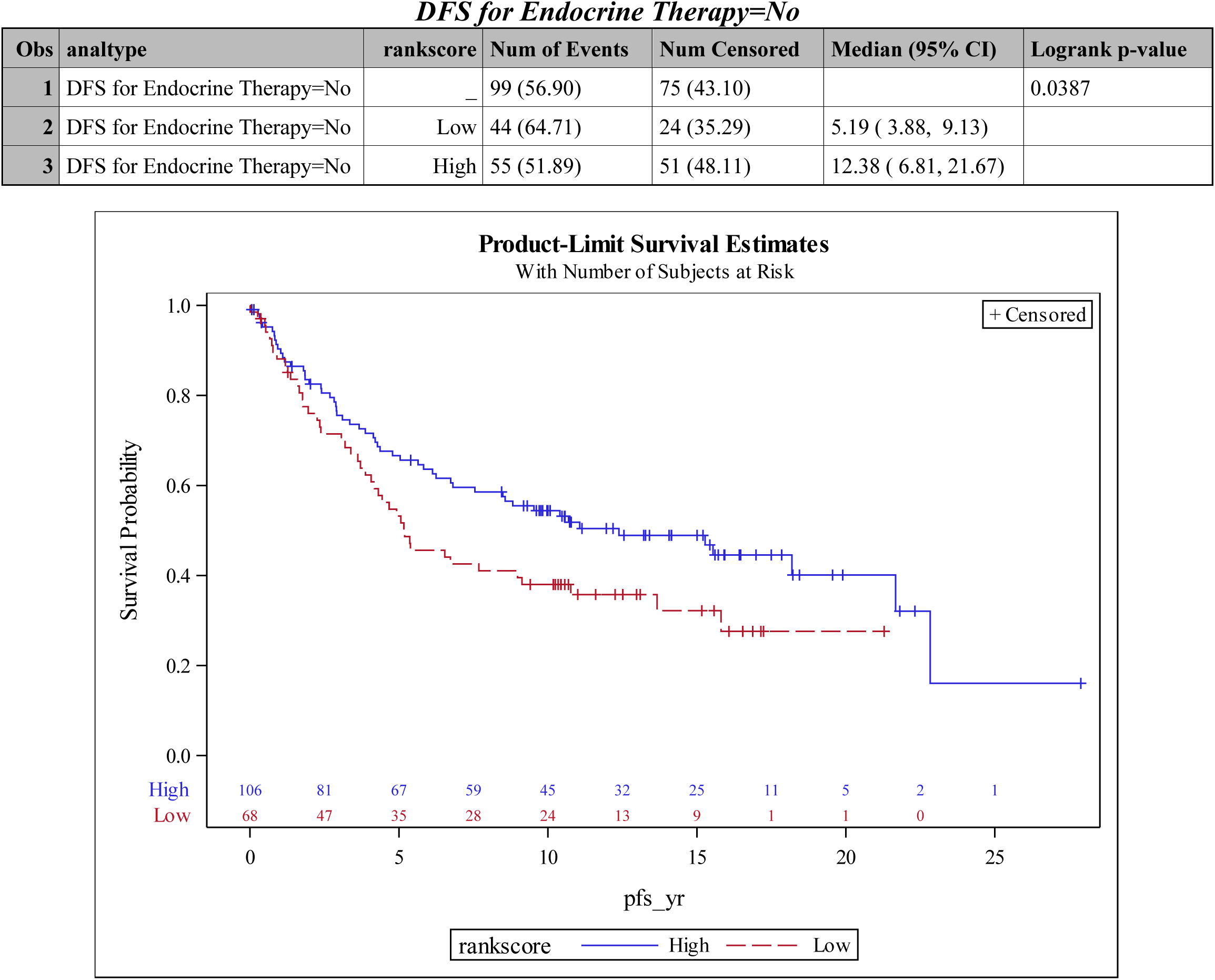
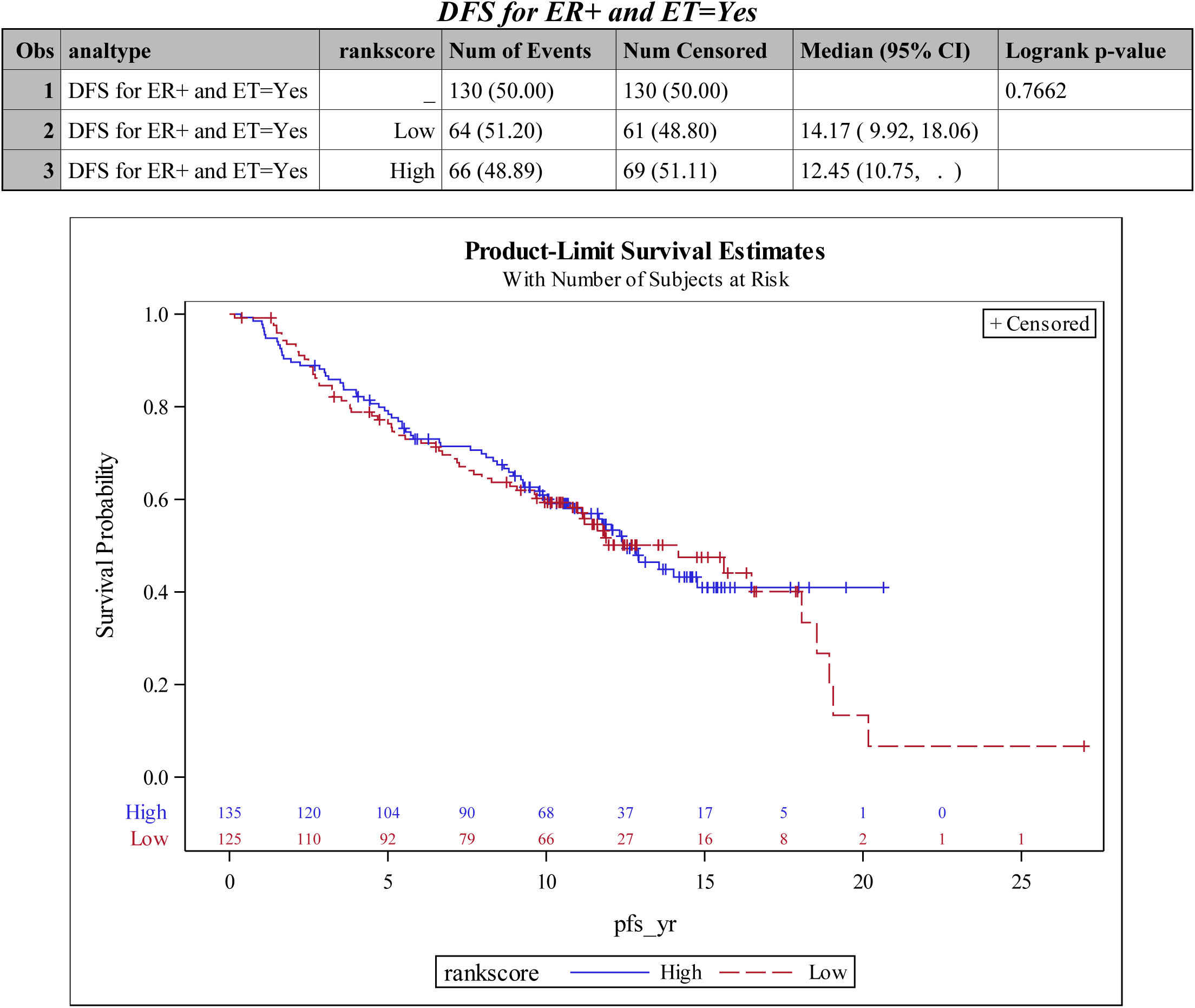
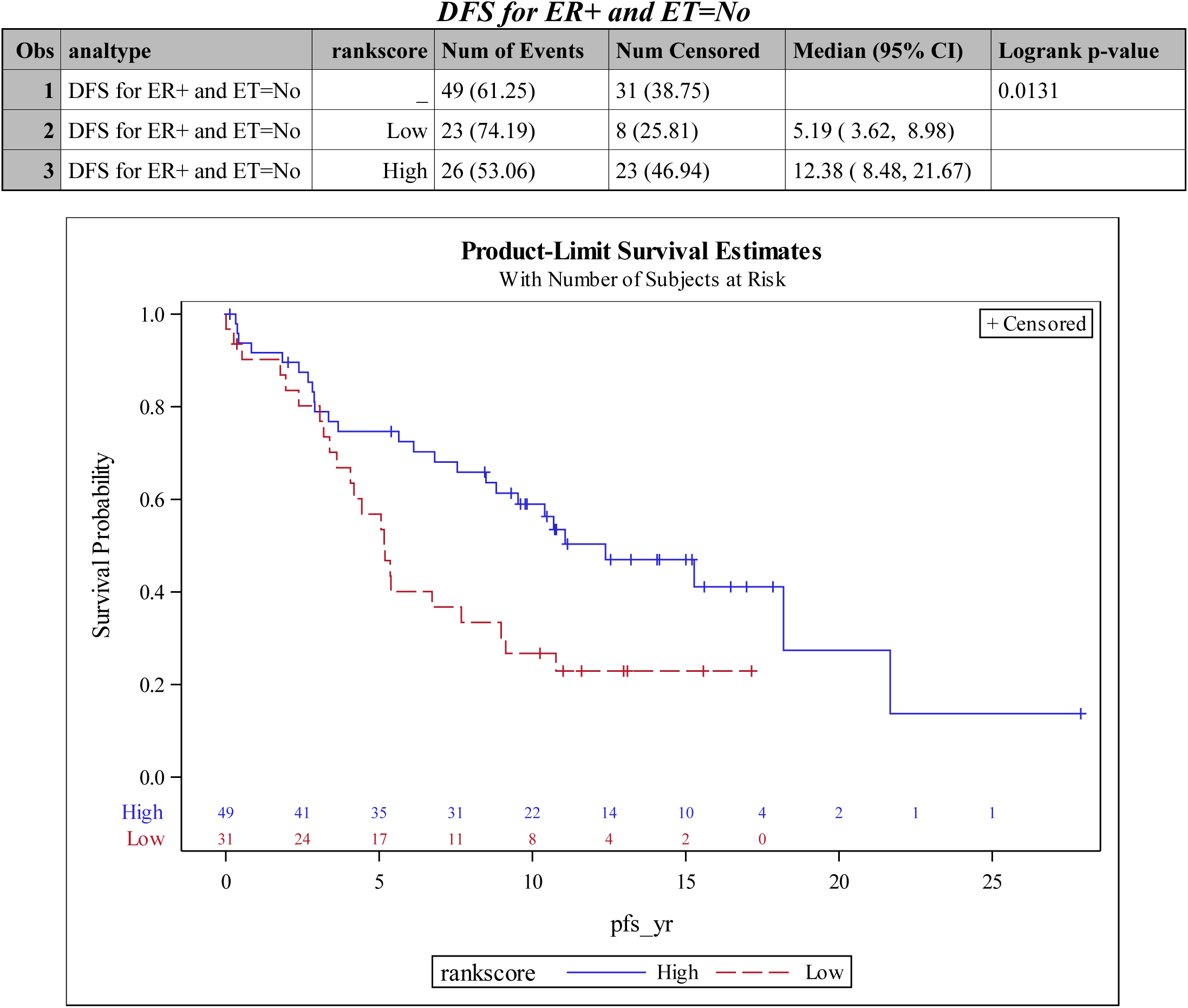
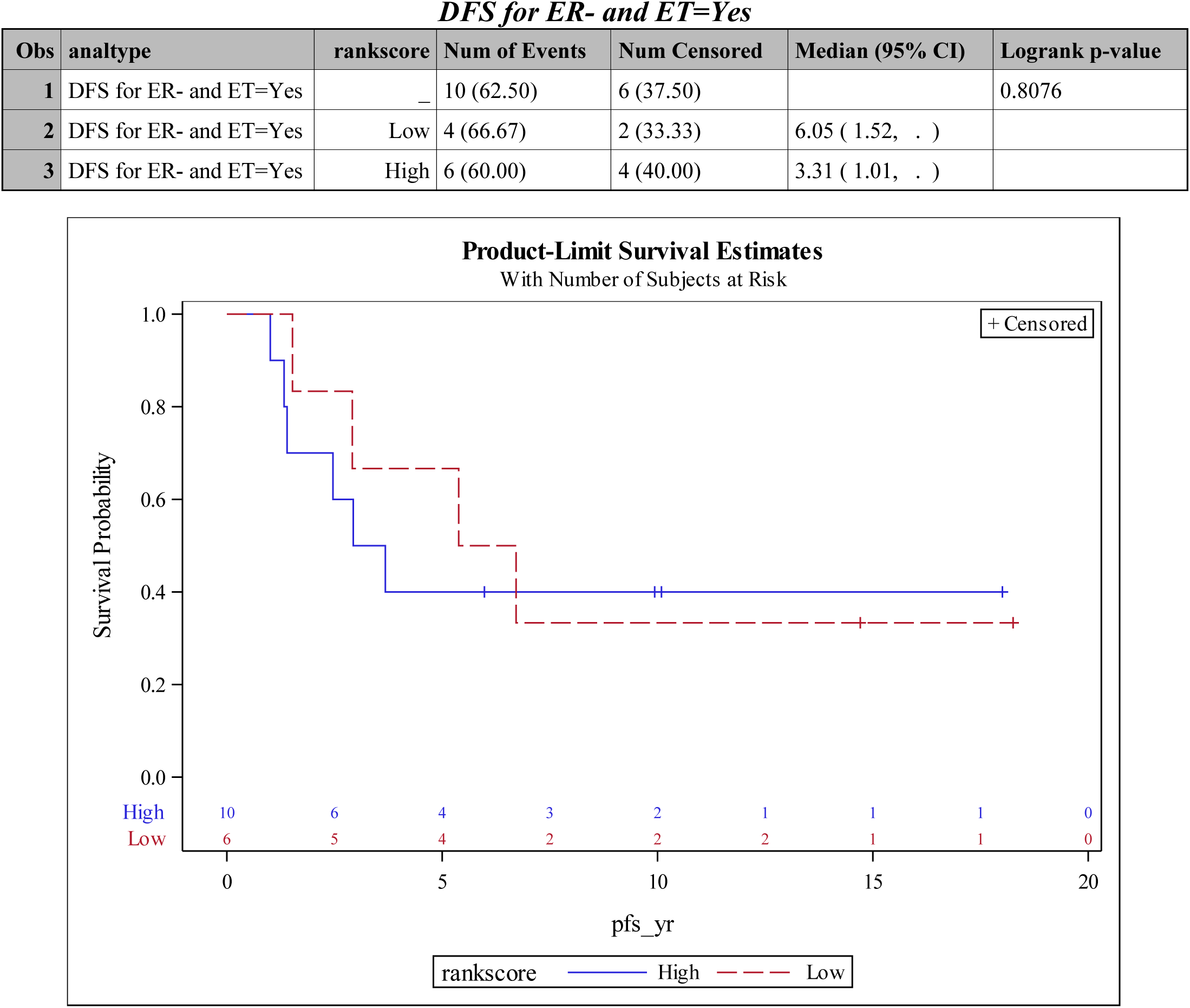
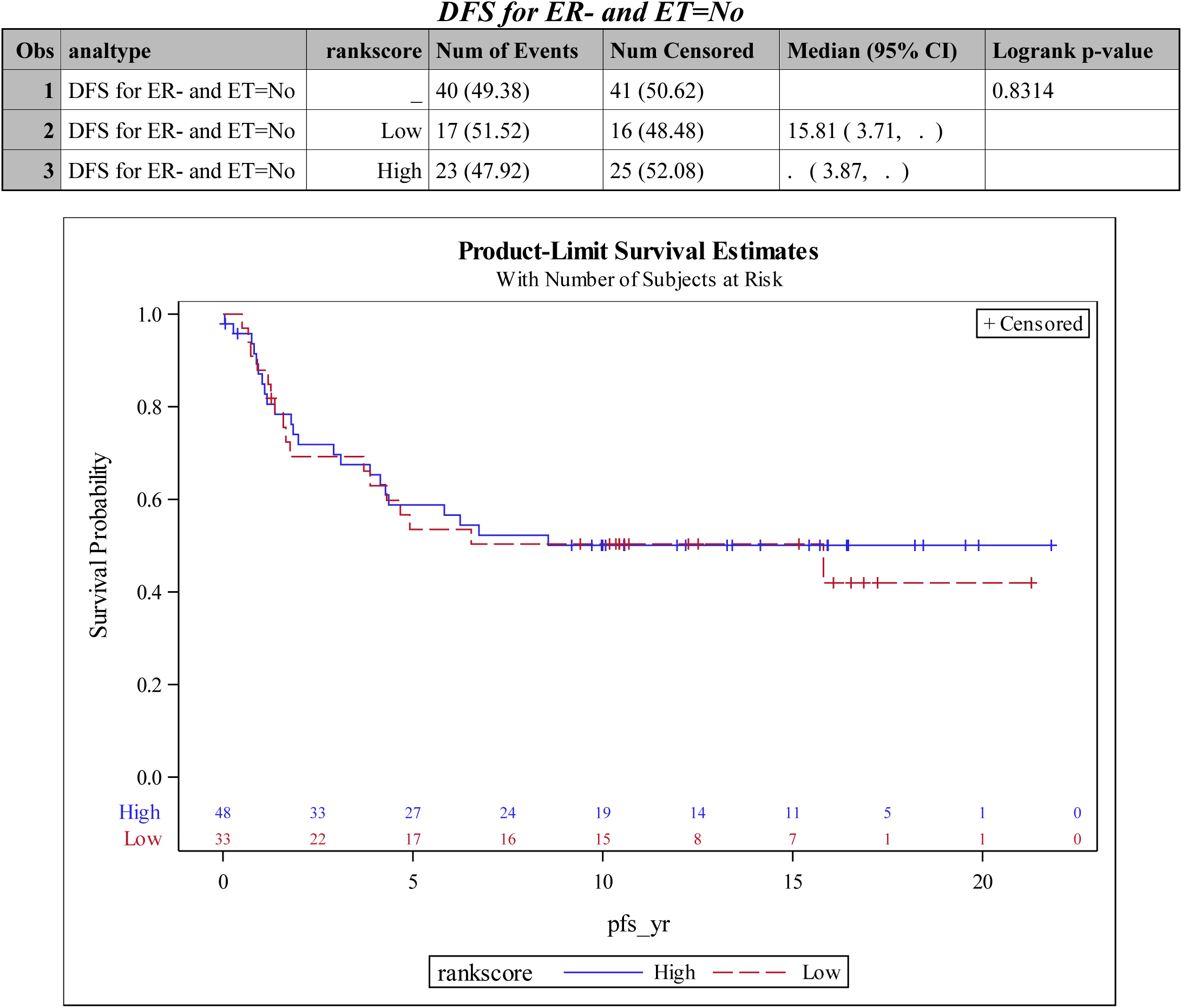
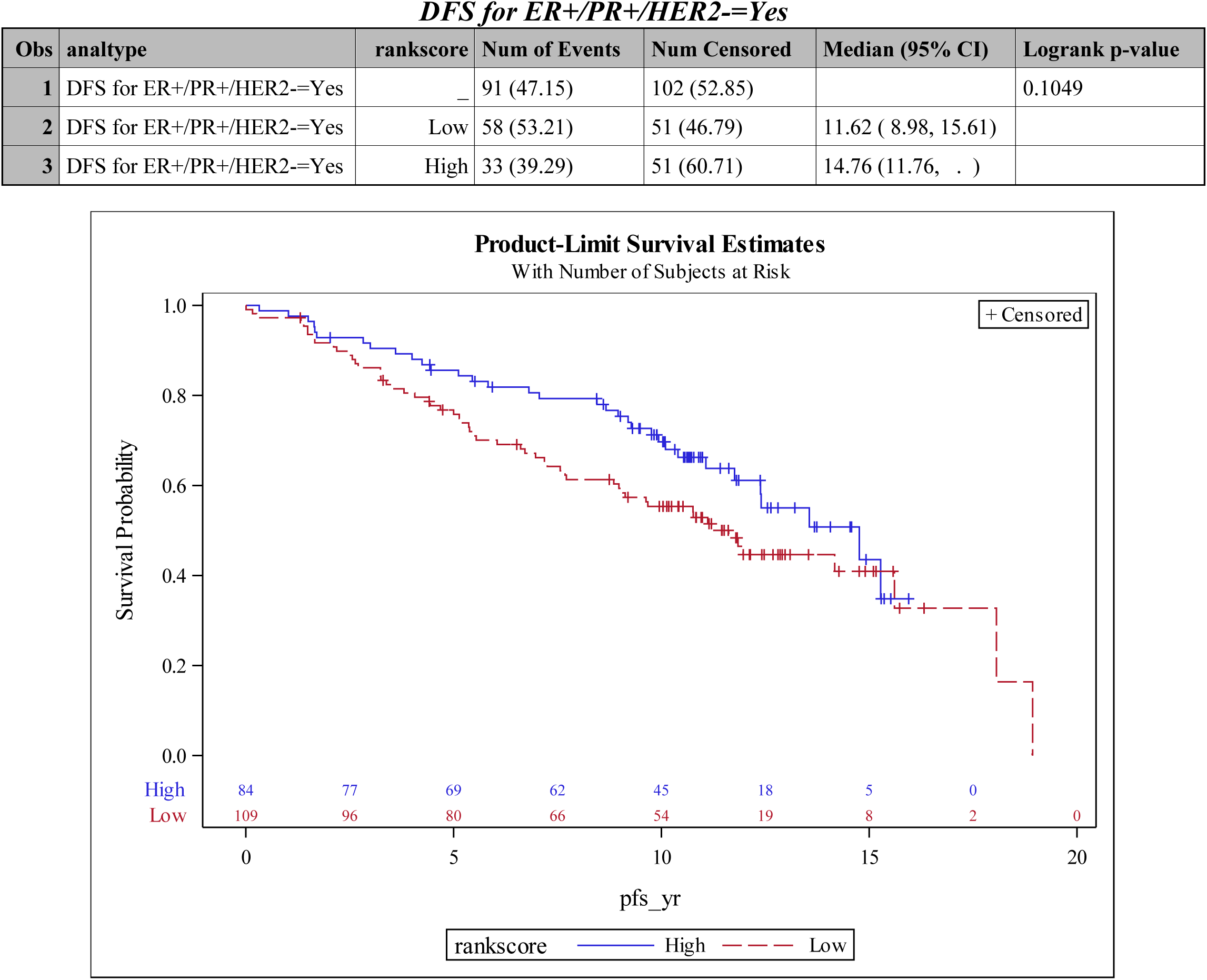
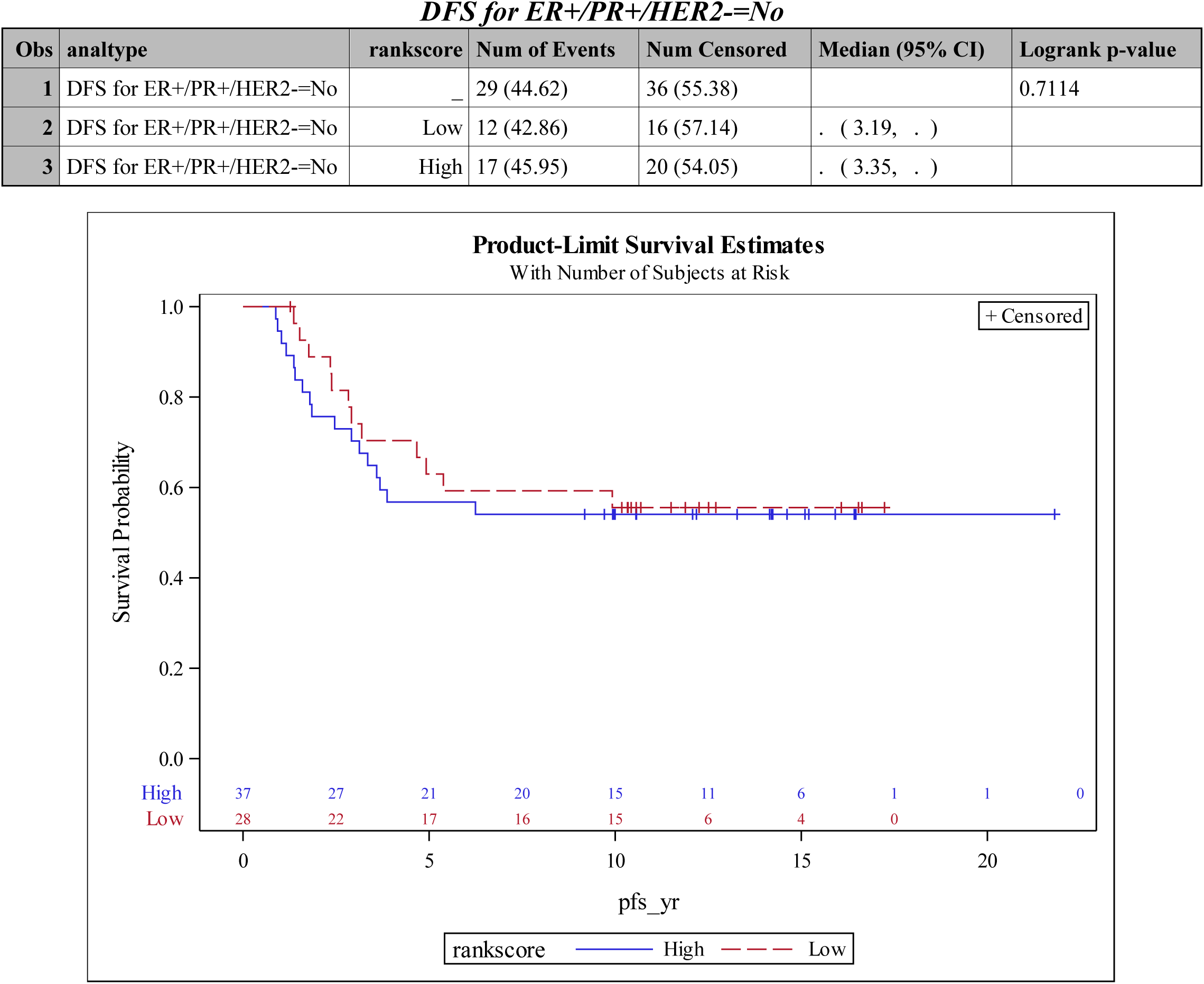
Summary of results for Overall Survival and DFS analysis – Univariate analyses on the PDK4 H-score category.

### Statistical Methods: TBX3

For subjects with multiple tumor samples available, we included only the sample with the highest TBX3 H-score. Wilcoxon Rank Sum and Kruskal-Wallis tests were used to determine if TBX3 H-scores were correlated with other tumor markers. Cox proportional hazards regression models were used to determine whether TBX3 H-scores and other variables were related to overall and disease-free survival either univariately or in multivariable models. In these analyses, TBX3 H-scores were divided into low and high categories at the score of 27.91721 for overall survival (time from surgery to death or censoring) and disease-free survival (time from surgery to first recurrence or censoring, excluding patients with M1 stage at surgery). These cutoff values were determined by using the maximum chi-square value for all score values between the 25th and 75th percentile (http://www.pharmasug.org/proceedings/2012/SP/PharmaSUG-2012-SP12.pdf). TBX3 high/low was included in all multivariable models. As a double check on the direction of the hazard ratio and as a more powerful test if the H-score effect was truly linear, we also fit multivariable models with the H-score as continuous.

We conducted subgroup analyses on overall survival using the ER-positive subgroup, endocrine therapy group, ER-positive on endocrine therapy, ER-negative, and ER+/PR+/HER2-. First, log-rank tests were done with the dichotomous H-score variable. Second, multivariable models with the H-score as dichotomous and then continuous were fit similar as was done in the main analyses.

Analyses were conducted using SAS Version 9.4. An α level of 5% was used to determine statistical significance.

### Results

Of the 557 patients who were part of the TMAs available to be read (TMA1-14) for TBX3, 457 patients (82%) had TBX3 values available/readable. Clinical parameters of the subjects included in the TMAs studied are summarized in Table 1 with grouping for patients with or without TBX3 values. Age, Tumor Stage, Recurrences, and TMA number were significantly different between the two groups (Age higher in missing group, T1 had more missing, Fewer Recurrences in missing group, and TMA4 and TMA6 had more missing, percentagewise). Since the numbers for the missing TBX3 values were so low, these significant differences were not investigated further.

### Correlation of TBX3 H-score with other disease markers

We compared TBX3 H-score expression with ER, PR, HER-2/neu, Nodal stage, Tumor Stage or Grade. TBX3 levels were correlated with Tumor Stage and Grade (Table 2). Tumor Stage 0/1 was significantly less than Stage 2 and 3, and Tumor Grade was significantly different among all grades with Grade 3 having the higher results.

### Overall Survival Analysis

#### Univariate

In univariate analyses, variables significantly related to overall survival in the Cox proportional hazards regression models were Tumor Grade, Tumor Stage, and Nodal Stage (Table 3a). Higher Tumor Grade, Higher Tumor Stage, and Nodal Stage-positive were correlated with lower survival. TBX3 H-score was not related to overall survival (log rank test p-value 0.2443).

#### Multivariable

In the multivariate analysis (Table 4a), Tumor Grade, Tumor Stage, Nodal Stage and TBX3 H-score were found to be significant. Higher Tumor Grade, Higher Tumor Stage, Nodal Stage-positive and lower TBX3 H-score were correlated with lower survival.

#### Subgroup analysis

For unadjusted tests (i.e. univariately), the results were significant for patients not having ER+/PR+/HER2- (log rank test p-value=0.035). Other analyses were not statistically significant (log rank test p-value 0.416 for ER-positive, 0.747 for ER-negative, 0.984 for patients on endocrine therapy, 0.289 for patients not on endocrine therapy, 0.557 for ER-positive and on endocrine therapy, and 0.057 for ER-positive and not on endocrine therapy). For the significant result, the lower TBX3 H-score was correlated to lower overall survival.

In multivariable models treating the H-score as dichotomous, H-score category was significant for patients who were ER-positive and not on endocrine therapy and patients who were not ER+/PR+/HER2- (Table 5a). For the results that were significant, the lower TBX3 H-score was correlated with lower survival. In multivariable models treating the H-score as continuous, the H-score was significant result in patients who were not ER+/PR+/HER2- (Table 6a).

In addition, Kaplan-Meier plots are provided for the univariate analyses using the categorical TBX3 H-score for overall and the subgroup analyses.

### Disease Free Survival Analysis

#### Univariate

In univariate analyses, variables significantly related to disease free survival in the Cox proportional hazards regression models were Tumor Grade, Tumor Stage, and Nodal Stage (Table 3a). Higher Tumor Grade, Higher Tumor Stage, and Nodal Stage-positive were correlated with lower disease free survival. TBX3 H-score was not related to disease free survival (log rank test p-value 0.1557).

#### Multivariable

In the multivariate analysis (Table 4b), Tumor Grade, Tumor Stage, Nodal Stage and TBX3 H-score were found to be significant. Higher Tumor Grade, Higher Tumor Stage, Nodal Stage-positive and lower TBX3 H-score were correlated with lower disease free survival.

#### Subgroup analysis

For unadjusted tests (i.e. univariately), the results were not statistically significant for any subgroup (log rank test p-value 0.137 for ER-positive, 0.610 for ER-negative, 0.385 for patients on endocrine therapy, 0.675 for patients not on endocrine therapy, 0.522 for ER-positive and on endocrine therapy and 0.384 for ER-positive and not on endocrine therapy).

In multivariable models treating the H-score as dichotomous, H-score category was significant for patients who were ER+ and patients who were not ER+/PR+/HER2- (Table 5b) where the patients with lower TBX3 H-score were correlated to lower disease free survival. In multivariable models treating the H-score as continuous, the H-score was significant result in patients who were not ER+/PR+/HER2- (Table 6b).

In addition, Kaplan-Meier plots are provided for the univariate analyses using the categorical TBX3 H-score for overall and the subgroup analyses.

**Table 1.** Description of the patients and characteristics of their tumors (n=557)

| **Characteristic** | **Have TBX3**<br>**(N=457)** | **Missing TBX3**<br>**(N=100)** | **p-value\*** |
| --- | --- | --- | --- |
| Age at Diagnosis, y |  |  | 0.0066 |
| Mean (SD) | 59.0 (15.01) | 63.5 (13.51) |  |
| Median | 58.12 | 64.30 |  |
| Range | 5.92-94.75 | 34.49-88.55 |  |
| Race, n(%) |  |  | 0.1647 |
| White | 351 (76.81%) | 85 (85.00%) |  |
| African American | 102 (22.32%) | 14 (14.00%) |  |
| Asian | 2 ( 0.44%) | 1 ( 1.00%) |  |
| Other | 1 ( 0.22%) | 0 |  |
| Unknown | 1 ( 0.22%) | 0 |  |
| PR Status, n(%) |  |  | 0.1986 |
| Negative | 147 (32.17%) | 27 (27.00%) |  |
| Positive | 273 (59.74%) | 69 (69.00%) |  |
| Unknown/Not Done | 37 ( 8.10%) | 4 ( 4.00%) |  |
| ER Status, n(%) |  |  | 0.2268 |
| Negative | 100 (21.88%) | 17 (17.00%) |  |
| Positive | 336 (73.52%) | 81 (81.00%) |  |
| Unknown/Not Done | 21 ( 4.60%) | 2 ( 2.00%) |  |
| HER-2/neu, n(%) |  |  | 1.0000 |
| Negative | 214 (46.83%) | 54 (54.00%) |  |
| Equivocal | 3 ( 0.66%) | 0 |  |
| Positive | 41 ( 8.97%) | 10 (10.00%) |  |
| Unknown/Not Done | 199 (43.54%) | 36 (36.00%) |  |
| Tumor Grade, n(%) |  |  | 0.3696 |
| I | 126 (27.57%) | 34 (34.00%) |  |
| II | 194 (42.45%) | 40 (40.00%) |  |
| III | 109 (23.85%) | 17 (17.00%) |  |
| IV | 1 ( 0.22%) | 0 |  |
| Unknown | 26 ( 5.69%) | 8 ( 8.00%) |  |
| Not Applicable | 1 ( 0.22%) | 1 ( 1.00%) |  |
| T Stage, *n*(%) |  |  | 0.0018 |
| T0 | 1 ( 0.22%) | 1 ( 1.00%) |  |
| T1 | 239 (52.30%) | 69 (69.00%) |  |
| T2 | 177 (38.73%) | 20 (20.00%) |  |
| T3 | 21 ( 4.60%) | 7 ( 7.00%) |  |
| T4 | 14 ( 3.06%) | 3 ( 3.00%) |  |
| TX/Unknown | 5 ( 1.09%) | 0 |  |
| N Stage, *n*(%) |  |  | 0.6023 |
| N0 | 274 (59.96%) | 64 (64.00%) |  |
| N1 | 117 (25.60%) | 23 (23.00%) |  |
| N2 | 26 ( 5.69%) | 3 ( 3.00%) |  |
| N3 | 8 ( 1.75%) | 1 ( 1.00%) |  |
| NX/Unknown | 31 ( 6.78%) | 9 ( 9.00%) |  |
| Not Applicable | 1 ( 0.22%) | 0 |  |
| M Stage, *n*(%) |  |  | 0.7417 |
| M0 | 228 (49.89%) | 62 (62.00%) |  |
| M1 | 12 ( 2.63%) | 2 ( 2.00%) |  |
| MX/Unknown | 211 (46.17%) | 36 (36.00%) |  |
| Not Applicable | 6 ( 1.31%) | 0 |  |
| Endocrine Therapy, *n*(%) |  |  | 0.4102 |
| Yes | 272 (59.52%) | 66 (66.00%) |  |
| No | 165 (36.11%) | 33 (33.00%) |  |
| Unknown/Missing | 20 ( 4.38%) | 1 ( 1.00%) |  |
| Death, *n*(%) |  |  | 0.4868 |
| Yes | 214 (46.83%) | 43 (43.00%) |  |
| No | 243 (53.17%) | 57 (57.00%) |  |
| Recurrence, *n*(%) |  |  | 0.0445 |
| Yes | 137 (29.98%) | 20 (20.00%) |  |
| No | 320 (70.02%) | 80 (80.00%) |  |
| TMA |  |  | 0.0414 |
| TMA-1 | 40 ( 8.75%) | 3 ( 3.00%) |  |
| TMA-2 | 33 ( 7.22%) | 10 (10.00%) |  |
| TMA-3 | 0 | 0 |  |
| TMA-4 | 29 ( 6.35%) | 13 (13.00%) |  |
| TMA-5 | 32 ( 7.00%) | 8 ( 8.00%) |  |
| TMA-6 | 27 ( 5.91%) | 10 (10.00%) |  |
| TMA-7 | 33 ( 7.22%) | 10 (10.00%) |  |
| TMA-8 | 40 ( 8.75%) | 4 ( 4.00%) |  |
| TMA-9 | 38 ( 8.32%) | 8 ( 8.00%) |  |
| TMA-10 | 38 ( 8.32%) | 3 ( 3.00%) |  |
| TMA-11 | 41 ( 8.97%) | 4 ( 4.00%) |  |
| TMA-12 | 35 ( 7.66%) | 11 (11.00%) |  |
| TMA-13 | 34 ( 7.44%) | 9 ( 9.00%) |  |
| TMA-14 | 37 ( 8.10%) | 7 ( 7.00%) |  |
| Follow-up, y, median<br>(range) | 12.9 (0.04-27.9) | 12.5 (0.3-19.2) | 0.4455 |
| Overall survival, y,<br>median (95% CI) | 13.7 (12.0, 18.5) | 18.2 (11.2, 19.1) | 0.4395 |
| Disease-free survival, y,<br>median (95% CI)** | 12.1 (10.7, 14.8) | 15.6 (11.2, 19.1) | 0.1321 |

\*from t-test, Chi-square, Fisher's Exact Test, or Log-Rank Test; Missing values not included.
\*\*M1 patients excluded from analysis

**Table 2.** Bivariate analysis of TBX3 H-score with other tumor markers

| Variable | TBX3 H-score Median (25<sup>th</sup> percentile, 75<sup>th</sup> percentile) |  |  |  |  |  | p-value* |
| --- | --- | --- | --- | --- | --- | --- | --- |
|  | n | Values | n | values | n | values |  |
|  | Negative |  | Positive |  |  |  |  |
| ER | 100 | 50.49 (24.17, 95.21) | 336 | 46.30 (24.69, 80.31) |  |  | 0.3715 |
| PR | 147 | 46.37 (23.03, 85.93) | 273 | 46.19 (24.50, 80.09) |  |  | 0.6406 |
| HER-2/neu | 214 | 47.66 (25.35, 87.92) | 14 | 54.31 (32.72, 78.01) |  |  | 0.8488 |
| ER+/PR+/HER- | 66 | 53.91 (26.64, 91.04) | 184 | 47.37 (25.70, 86.28) |  |  | 0.7202 |
| Nodal Status | 274 | 49.66 (23.83, 82.14) | 151 | 45.46 (28.27, 82.79) |  |  | 0.9714 |
|  | Grade 1 |  | Grade 2 |  | Grade 3 |  |  |
| Tumor Grade | 126 | 39.10 (16.86, 61.25) | 194 | 46.11 (25.38, 78.28) | 110 | 72.81 (39.98, 113.79) | <.0001** |
|  | T0/1 |  | T2 |  | T3/4 |  |  |
| Tumor Stage | 240 | 45.58 (21.70, 76.81) | 177 | 49.30 (28.27, 87.66) | 35 | 60.94 (36.19, 101.64) | 0.0063*** |

\*from Wilcoxon Rank Sum test for Hormone Receptor Status and Kruskal-Wallis test for Tumor Grade and Tumor Stage
\*\*the difference between Grade 1 and 2 was significant (p=0.0068), Grade 1 and 3 (p<0.0001), and between Grade 2 and 3 (p<0.0001).
\*\*\*the difference between Stage 0/1 and 2 was significant (p=0.0238), and Stage 0/1 and 3/4 (p=0.0069).

**Table 3a.** Univariate analysis of other tumor markers for Overall Survival

| **Variable** | **n** | **Comparison\*** | **HR (95% CI)** | **p-value\*\*** |
| --- | --- | --- | --- | --- |
| ER | 534 | ER- vs ER+ | 0.96 ( 0.70, 1.31) | 0.7757 |
| PR | 516 | PR- vs PR+ | 1.23 ( 0.94, 1.62) | 0.1263 |
| HER-2/neu | 319 | HER-2/neu- vs HER-2/neu+ | 0.66 ( 0.42, 1.02) | 0.0594 |
| ER+/PR+/HER- | 314 | No vs Yes | 1.36 (0.93, 2.00) | 0.1161 |
| Tumor Grade | 521 | Grade 1 vs Grade 2 | 0.91 ( 0.66, 1.25) | <.0001 |
|  |  | Grade 1 vs Grade 3 | 0.49 ( 0.35, 0.69) |  |
|  |  | Grade 2 vs Grade 3 | 0.54 ( 0.40, 0.73) |  |
| T Stage | 552 | T0/1 vs T2 | 0.70 ( 0.53, 0.91) | <.0001 |
|  |  | T0/1 vs T3/4 | 0.34 ( 0.23, 0.50) |  |
|  |  | T2 vs T3/4 | 0.49 ( 0.33, 0.72) |  |
| N Stage | 516 | N+ vs N0 | 1.78 ( 1.37, 2.31) | <.0001 |

\*referent group listed second
\*\*from Wald Chi-square test

**Table 3b.** Univariate analysis of other tumor markers for Disease Free Survival

| **Variable** | **n** | **Comparison\*** | **HR (95% CI)** | **p-value\*\*** |
| --- | --- | --- | --- | --- |
| ER | 520 | ER- vs ER+ | 1.01 ( 0.75, 1.37) | 0.9330 |
| PR | 503 | PR- vs PR+ | 1.20 ( 0.92, 1.56) | 0.1758 |
| HER-2/neu | 311 | HER-2/neu- vs HER-2/neu+ | 0.74 ( 0.48, 1.15) | 0.1856 |
| ER+/PR+/HER- | 306 | No vs Yes | 1.15 (0.79, 1.68) | 0.4743 |
| Tumor Grade | 509 | Grade 1 vs Grade 2 | 0.86 ( 0.63, 1.16) | <.0001 |
|  |  | Grade 1 vs Grade 3 | 0.49 ( 0.35, 0.68) |  |
|  |  | Grade 2 vs Grade 3 | 0.57 ( 0.42, 0.77) |  |
| T Stage | 538 | T0/1 vs T2 | 0.62 ( 0.48, 0.80) | <.0001 |
|  |  | T0/1 vs T3/4 | 0.40 ( 0.26, 0.60) |  |
|  |  | T2 vs T3/4 | 0.64 ( 0.42, 0.97) |  |
| N Stage | 504 | N+ vs N0 | 1.68 ( 1.30, 2.17) | <.0001 |

\*referent group listed second
\*\*from Wald Chi-square test

**Table 4a.** Multivariable analysis for Overall Survival (N=456)

| Variable | Comparison* | HR | Lower<br>95% CI | Upper<br>95% CI | p-value** |
| --- | --- | --- | --- | --- | --- |
| Tumor Grade | Grade 1 vs Grade 2 | 0.881 | 0.610 | 1.273 | 0.0263 |
|  | Grade 1 vs Grade 3 | 0.574 | 0.380 | 0.866 |  |
|  | Grade 1 vs Unknown | 0.665 | 0.377 | 1.174 |  |
|  | Grade 2 vs Grade 3 | 0.651 | 0.463 | 0.915 |  |
|  | Grade 2 vs Unknown | 0.755 | 0.441 | 1.291 |  |
|  | Grade 3 vs Unknown | 1.160 | 0.657 | 2.047 |  |
| T Stage | T0/1 vs T2 | 0.765 | 0.558 | 1.050 | 0.0093 |
|  | T0/1 vs T3/4 | 0.449 | 0.272 | 0.741 |  |
|  | T0/1 vs TX | 2.128 | 0.492 | 9.215 |  |
|  | T2 vs T3/4 | 0.586 | 0.371 | 0.928 |  |
|  | T2 vs TX | 2.782 | 0.645 | 12.006 |  |
|  | T3/4 vs TX | 4.743 | 1.042 | 21.593 |  |
| N Stage | N+ vs N0 | 1.658 | 1.211 | 2.269 | <.0001 |
|  | N+ vs NX | 0.495 | 0.297 | 0.825 |  |
|  | N0 vs NX | 0.299 | 0.184 | 0.485 |  |
| TBX3 H-score<br>Category | High vs Low | 0.721 | 0.533 | 0.974 | 0.0333 |

\*referent group listed second
\*\*from Wald Chi-square test

**Table 4b.** Multivariable analysis for Disease Free Survival (n=444)

| **Variable** | **Comparison\*** | **HR** | **Lower**<br>**95% CI** | **Upper**<br>**95% CI** | **p-value\*\*** |
| --- | --- | --- | --- | --- | --- |
| Tumor Grade | Grade 1 vs Grade 2 | 0.840 | 0.594 | 1.188 | 0.0129 |
|  | Grade 1 vs Grade 3 | 0.539 | 0.362 | 0.801 |  |
|  | Grade 1 vs Unknown | 0.705 | 0.398 | 1.246 |  |
|  | Grade 2 vs Grade 3 | 0.641 | 0.460 | 0.895 |  |
|  | Grade 2 vs Unknown | 0.839 | 0.487 | 1.445 |  |
|  | Grade 3 vs Unknown | 1.308 | 0.730 | 2.343 |  |
| T Stage | T0/1 vs T2 | 0.660 | 0.489 | 0.890 | 0.0203 |
|  | T0/1 vs T3/4 | 0.543 | 0.323 | 0.916 |  |
|  | T0/1 vs TX | 1.312 | 0.380 | 4.534 |  |
|  | T2 vs T3/4 | 0.824 | 0.507 | 1.338 |  |
|  | T2 vs TX | 1.988 | 0.579 | 6.831 |  |
|  | T3/4 vs TX | 2.414 | 0.648 | 8.986 |  |
| N Stage | N+ vs N0 | 1.577 | 1.169 | 2.126 | 0.0002 |
|  | N+ vs NX | 0.657 | 0.387 | 1.115 |  |
|  | N0 vs NX | 0.417 | 0.252 | 0.690 |  |
| TBX3 H-score<br>Category | High vs Low | 0.689 | 0.516 | 0.922 | 0.0121 |

\*referent group listed second
\*\*from Wald Chi-square test

**Table 5a:** Overall Survival - PROC PHREG with H score as dichotomous variable in multivariable models

| **Obs** | **Group** | **H-Score**<br>**Category**<br>**Parameter\*** | **p-value** | **Point**<br>**Estimate** | **Lower**<br>**95%**<br>**Wald**<br>**Confidenc**<br>**e Limit** | **Upper**<br>**95%**<br>**Wald**<br>**Confidenc**<br>**e Limit** |
| --- | --- | --- | --- | --- | --- | --- |
| **1** | OS | High vs Low | 0.0333 | 0.721 | 0.533 | 0.974 |
| **2** | OS for ER+ | High vs Low | 0.1884 | 0.789 | 0.554 | 1.123 |
| **3** | OS for ER- | High vs Low | 0.0709 | 0.503 | 0.238 | 1.060 |
| **4** | OS for Endocrine<br>Therapy=Yes | High vs Low | 0.3786 | 0.837 | 0.564 | 1.243 |
| **5** | OS for Endocrine Therapy=No | High vs Low | 0.0505 | 0.589 | 0.346 | 1.001 |
| **6** | OS for ER+ and ET=Yes | High vs Low | 0.9396 | 0.983 | 0.637 | 1.519 |
| **7** | OS for ER+ and ET=No | High vs Low | 0.0323 | 0.422 | 0.191 | 0.930 |
| **8** | OS for ER- and ET=Yes | High vs Low | 0.3166 | 0.324 | 0.036 | 2.939 |
| **9** | OS for ER- and ET=No | High vs Low | 0.1874 | 0.559 | 0.235 | 1.327 |
| **10** | OS for ER+/PR+/HER2-=Yes | High vs Low | 0.4029 | 0.792 | 0.458 | 1.368 |
| **11** | OS for ER+/PR+/HER2-=No | High vs Low | 0.0036 | 0.250 | 0.099 | 0.635 |

\*referent group listed second

**Table 5b:**
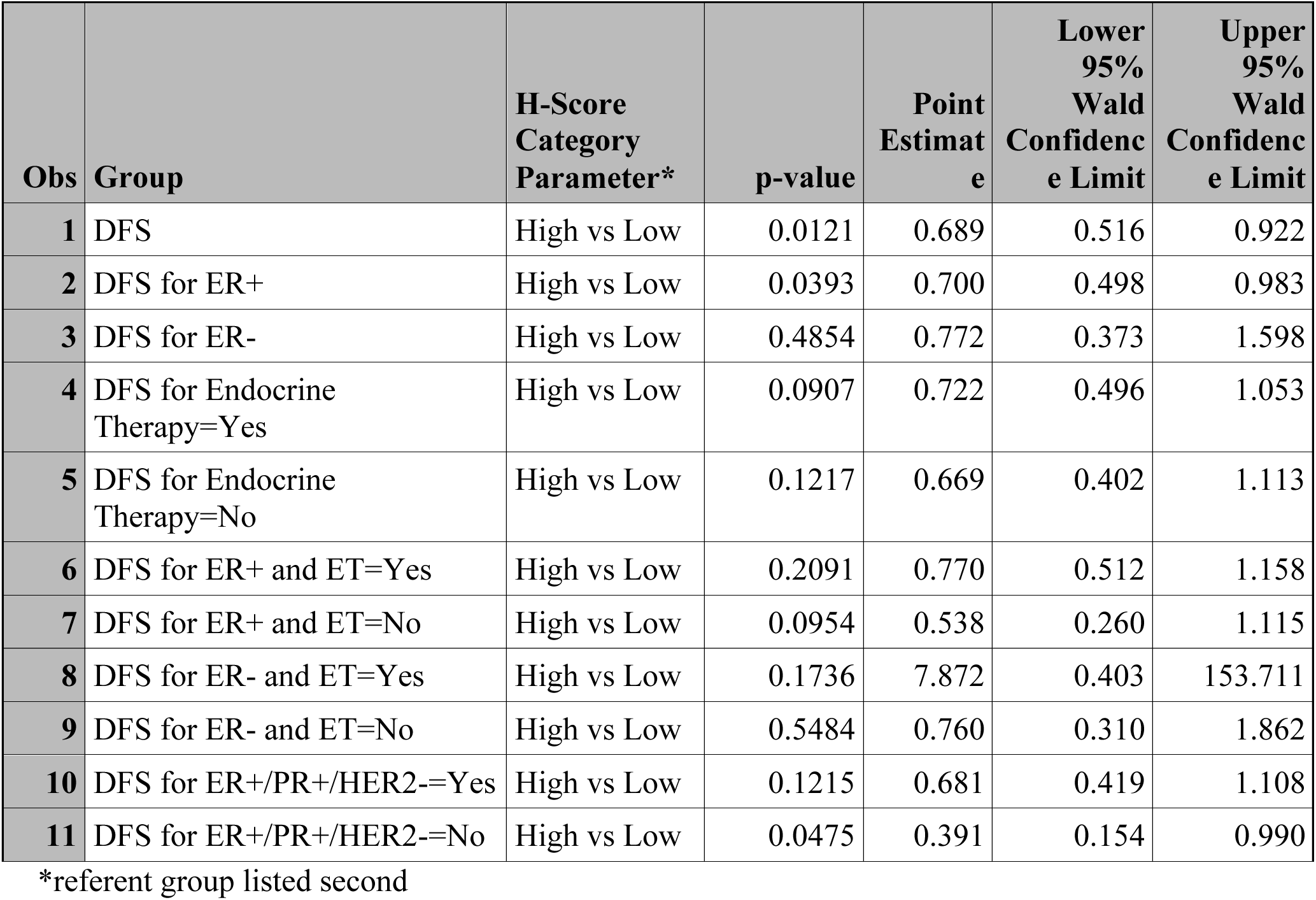
Disease Free Survival - PROC PHREG with H score as dichotomous variable in multivariable models

**Table 6a:** Overall Survival - PROC PHREG with H score as continuous variable in multivariable models

| **Obs** | **Group** | **Parameter** | **p-value** | **Point**<br>**Estimate** | **Lower**<br>**95%**<br>**Wald**<br>**Confidence**<br>**Limit** | **Upper**<br>**95%**<br>**Wald**<br>**Confidence**<br>**Limit** |
| --- | --- | --- | --- | --- | --- | --- |
| **1** | OS | score | 0.1414 | 0.997 | 0.994 | 1.001 |
| **2** | OS for ER+ | score | 0.6979 | 0.999 | 0.995 | 1.004 |
| **3** | OS for ER- | score | 0.0534 | 0.992 | 0.984 | 1.000 |
| **4** | OS for Endocrine<br>Therapy=Yes | score | 0.8304 | 0.999 | 0.994 | 1.005 |
| **5** | OS for Endocrine<br>Therapy=No | score | 0.4860 | 0.998 | 0.992 | 1.004 |
| **6** | OS for ER+ and ET=Yes | score | 0.9438 | 1.000 | 0.995 | 1.006 |
| **7** | OS for ER+ and ET=No | score | 0.6027 | 0.997 | 0.987 | 1.007 |
| **8** | OS for ER- and ET=Yes | score | 0.2253 | 0.952 | 0.880 | 1.031 |
| **9** | OS for ER- and ET=No | score | 0.2698 | 0.995 | 0.987 | 1.004 |
| **10** | OS for ER+/PR+/HER2-<br>=Yes | score | 0.9418 | 1.000 | 0.993 | 1.006 |
| **11** | OS for ER+/PR+/HER2-<br>=No | score | 0.0034 | 0.978 | 0.964 | 0.993 |

**Table 6b:** Disease Free Survival - PROC PHREG with H score as continuous variable in multivariable models

| **Obs** | **Group** | **Parameter** | **p-value** | **Point Estimate** | **Lower 95% Wald Confidence Limit** | **Upper 95% Wald Confidence Limit** |
| --- | --- | --- | --- | --- | --- | --- |
| **1** | DFS | score | 0.2099 | 0.998 | 0.994 | 1.001 |
| **2** | DFS for ER+ | score | 0.2528 | 0.997 | 0.993 | 1.002 |
| **3** | DFS for ER- | score | 0.4409 | 0.997 | 0.990 | 1.004 |
| **4** | DFS for Endocrine Therapy=Yes | score | 0.3920 | 0.998 | 0.992 | 1.003 |
| **5** | DFS for Endocrine Therapy=No | score | 0.8138 | 0.999 | 0.994 | 1.005 |
| **6** | DFS for ER+ and ET=Yes | score | 0.4605 | 0.998 | 0.992 | 1.003 |
| **7** | DFS for ER+ and ET=No | score | 0.4776 | 0.997 | 0.988 | 1.005 |
| **8** | DFS for ER- and ET=Yes | score | 0.2881 | 1.056 | 0.955 | 1.169 |
| **9** | DFS for ER- and ET=No | score | 0.7594 | 0.999 | 0.991 | 1.007 |
| **10** | DFS for ER+/PR+/HER2-=Yes | score | 0.4490 | 0.998 | 0.992 | 1.004 |
| **11** | DFS for ER+/PR+/HER2-=No | score | 0.0378 | 0.986 | 0.973 | 0.999 |

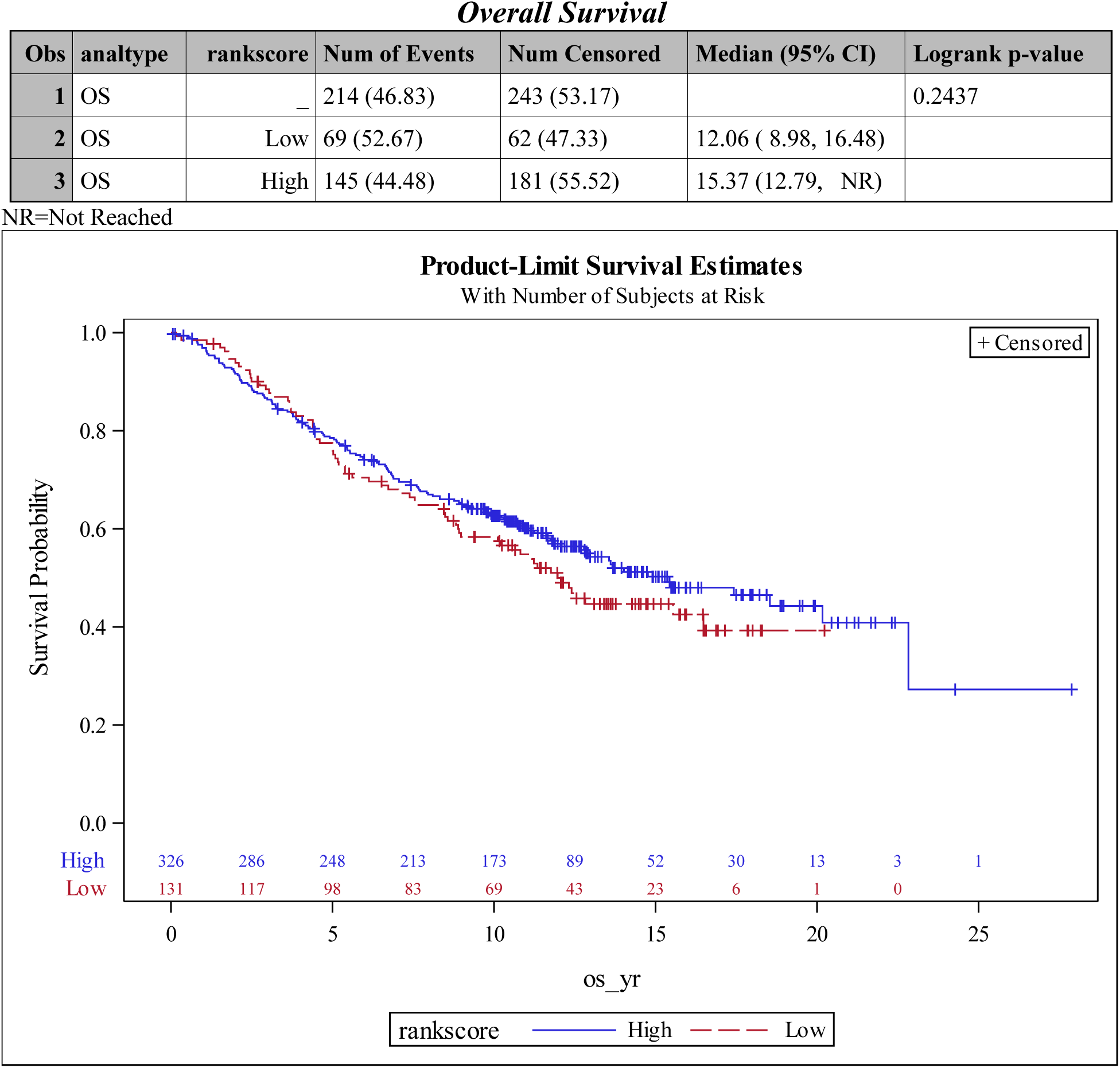
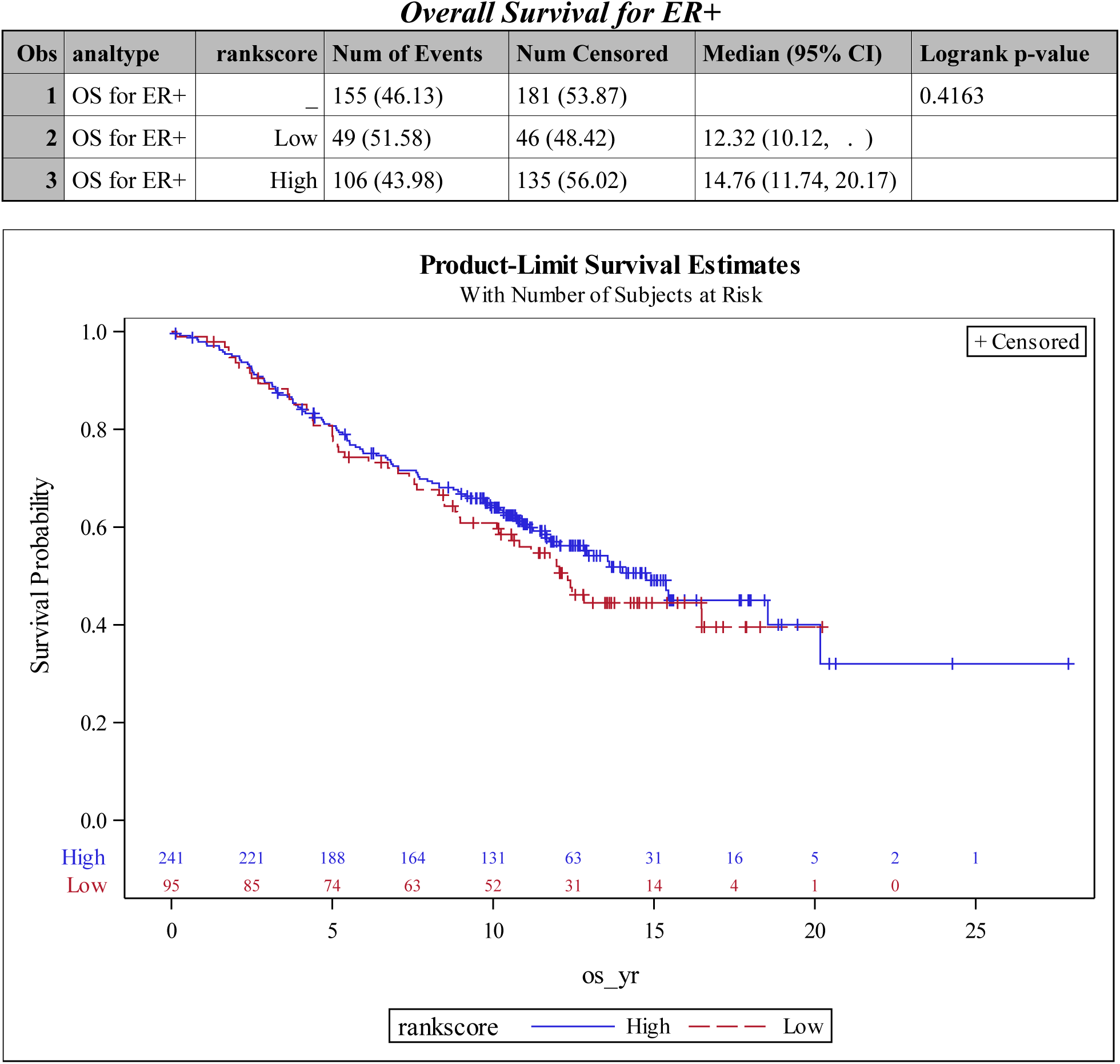
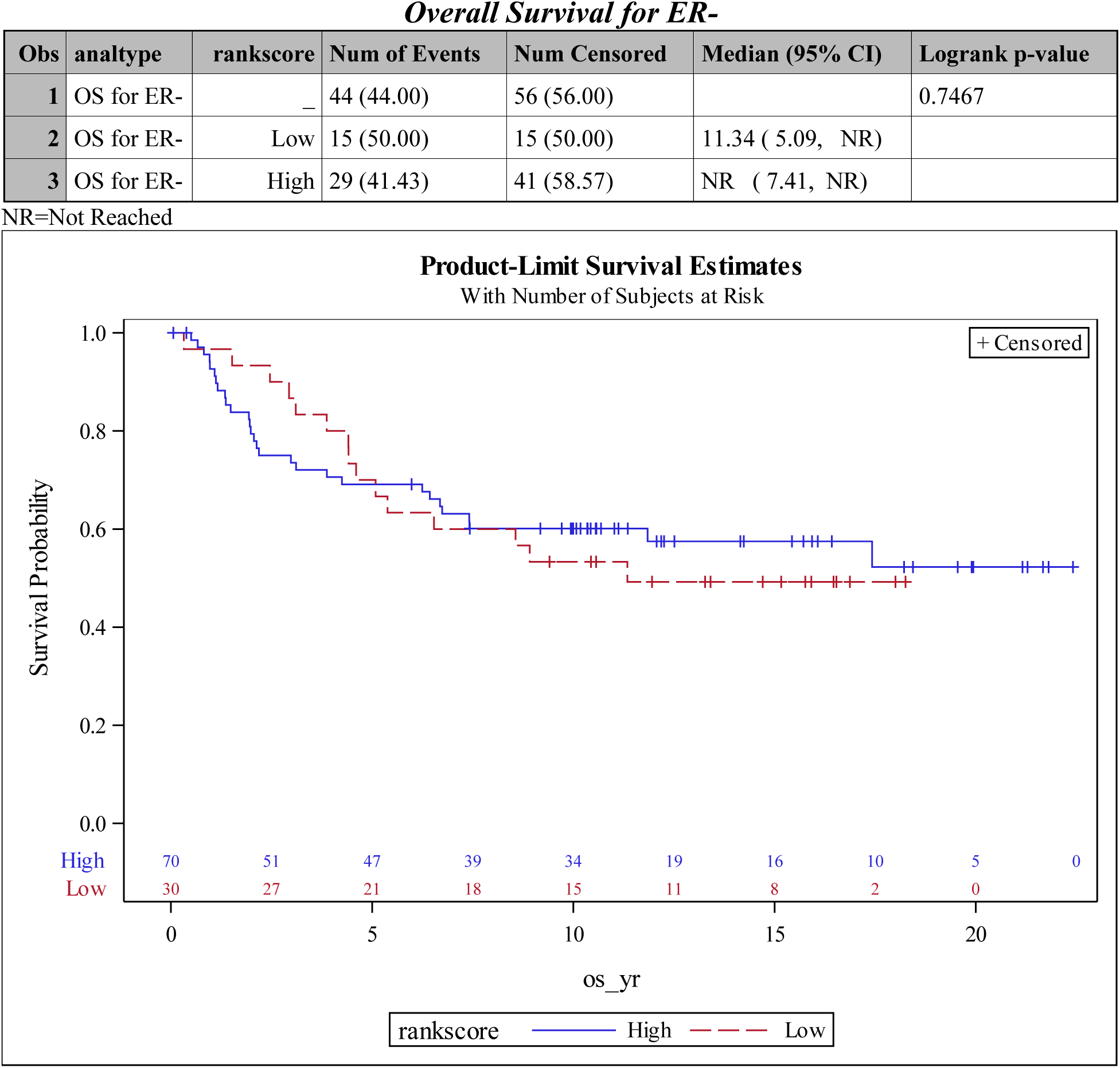
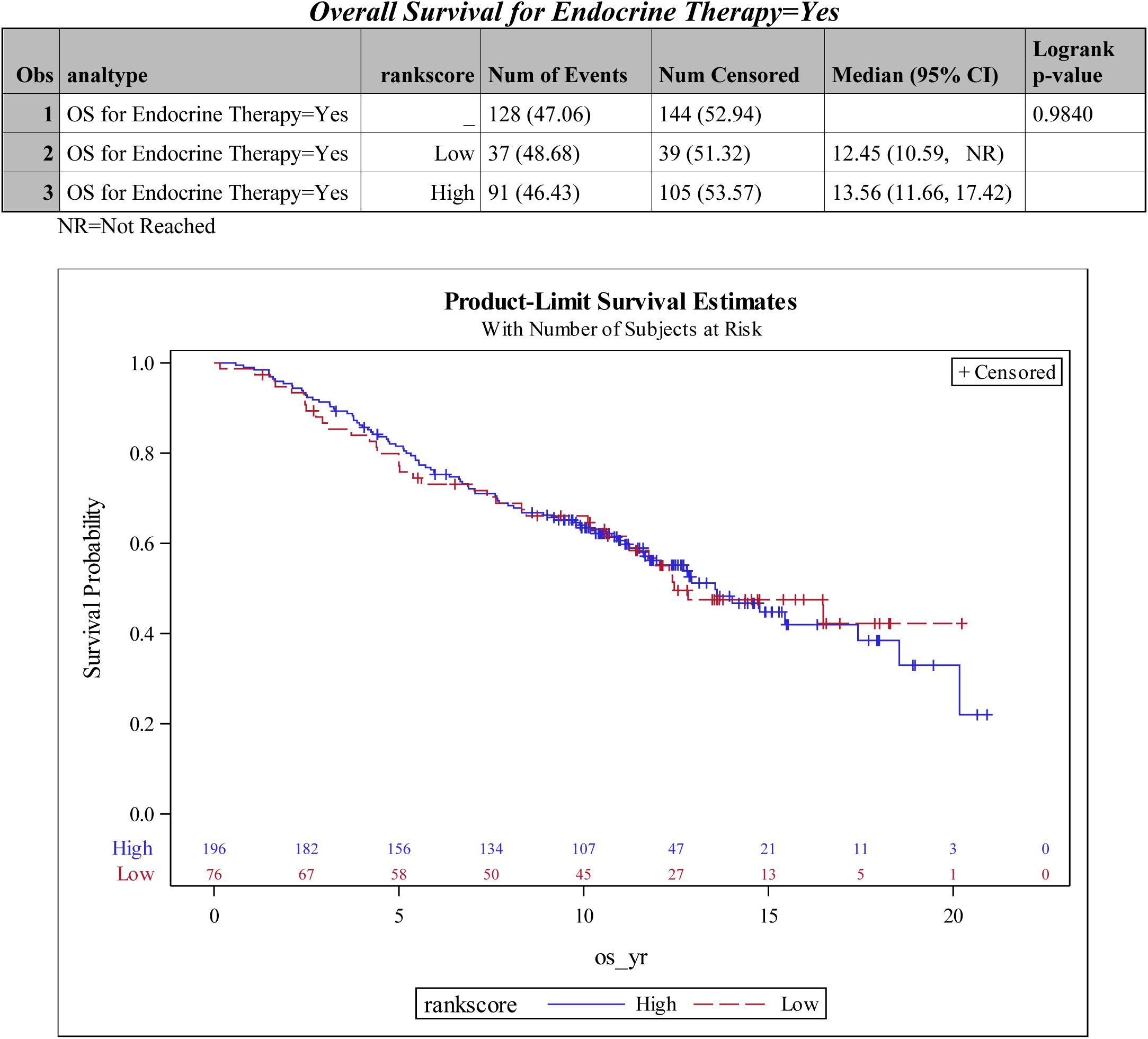
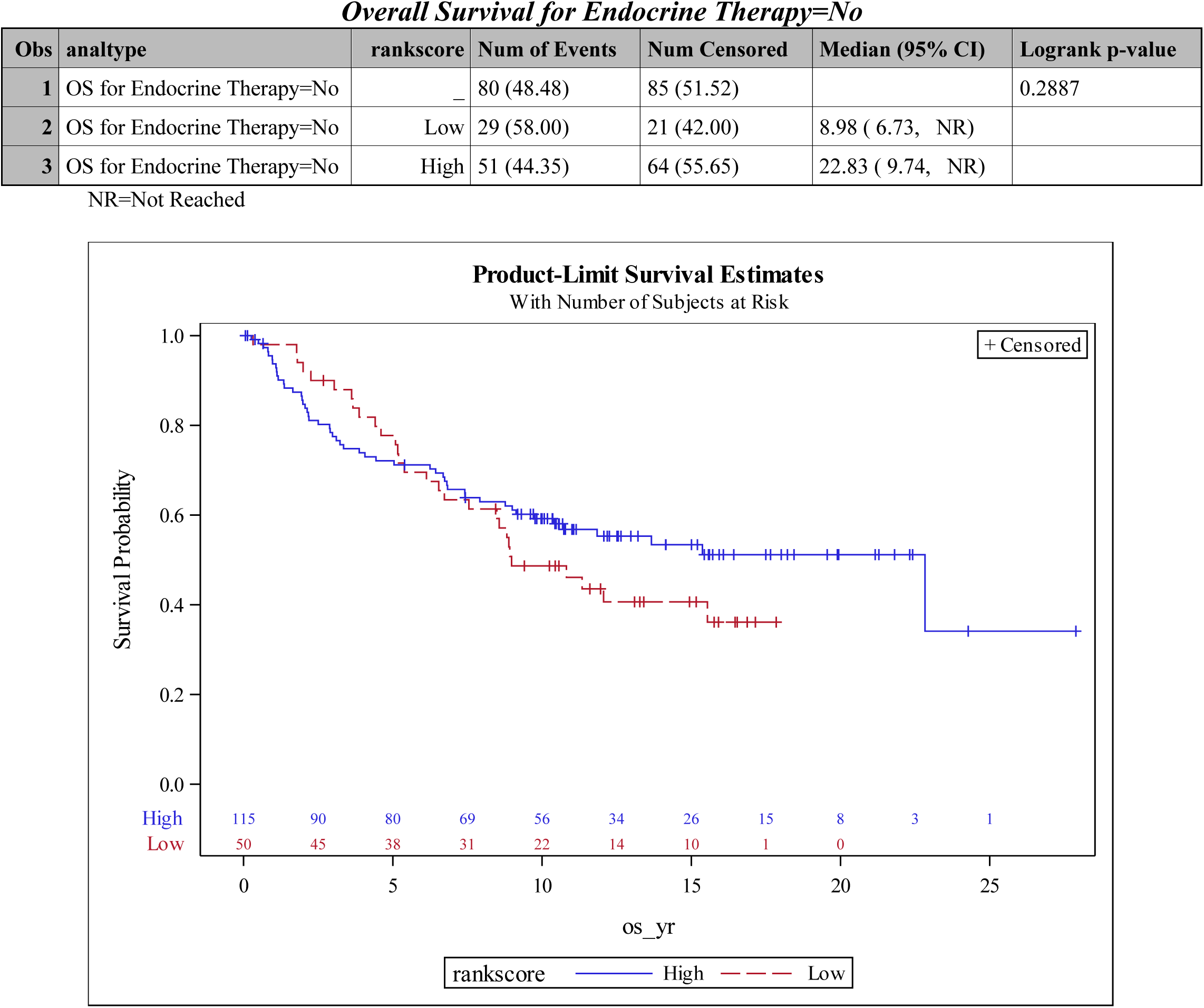
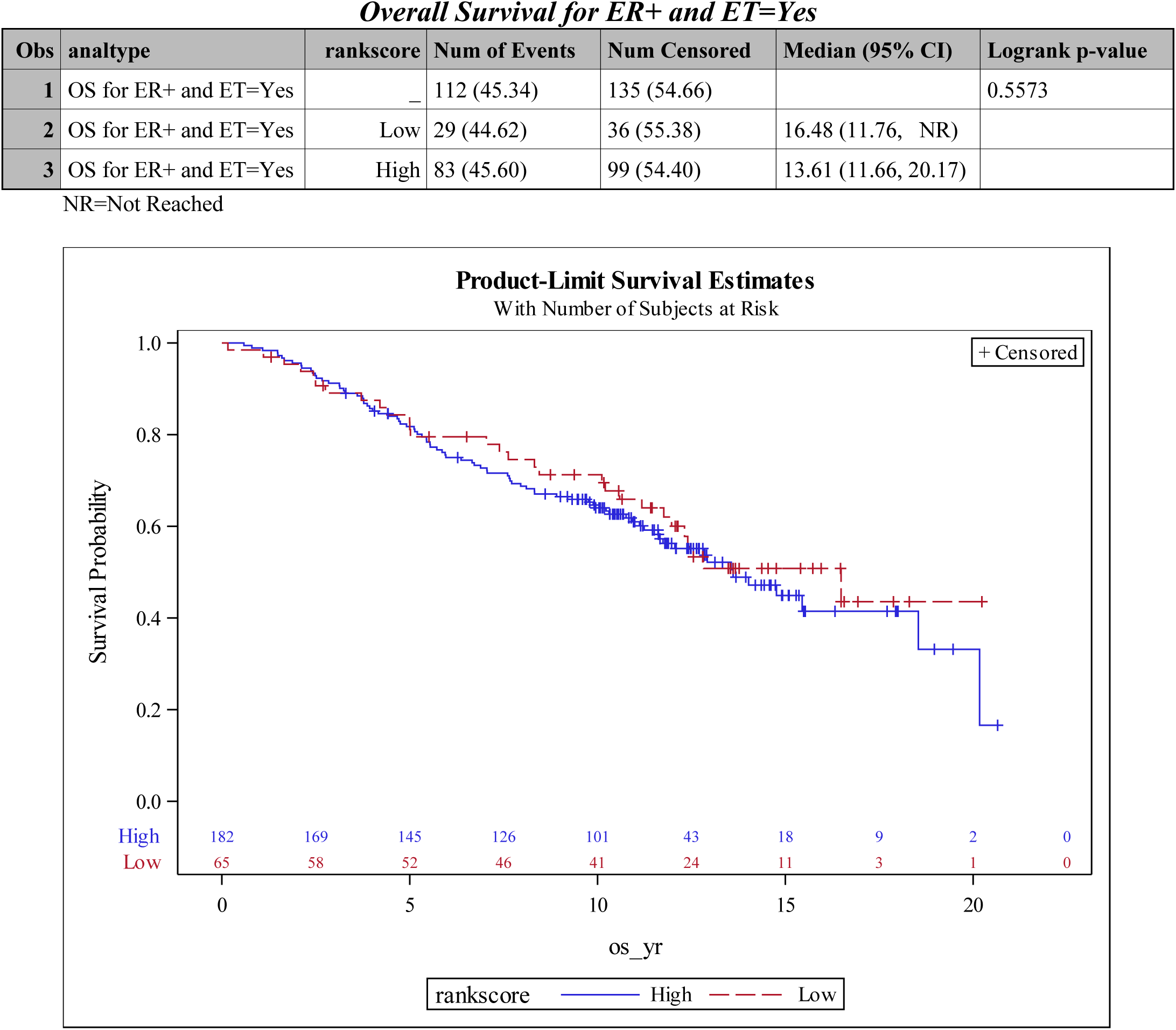
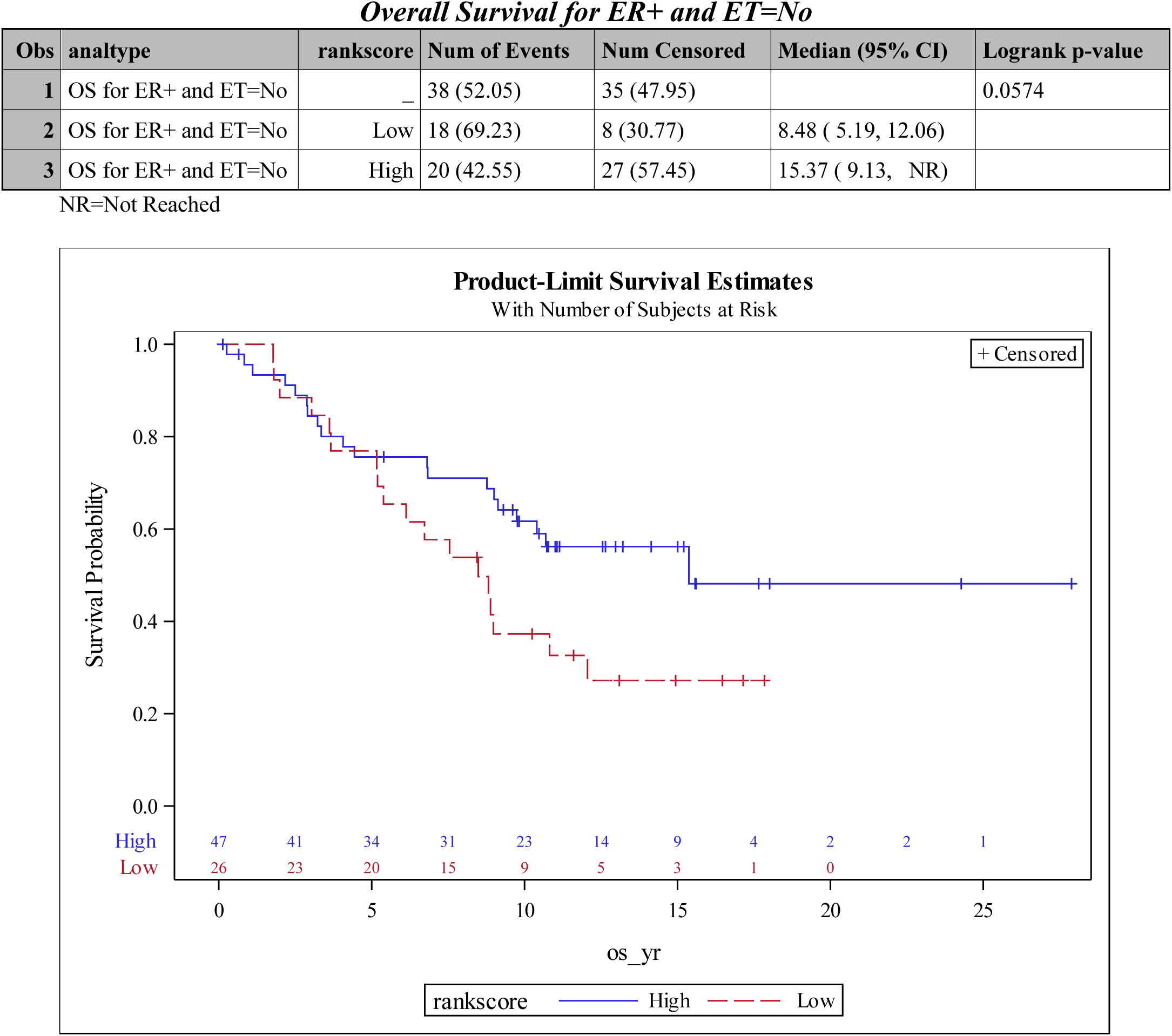
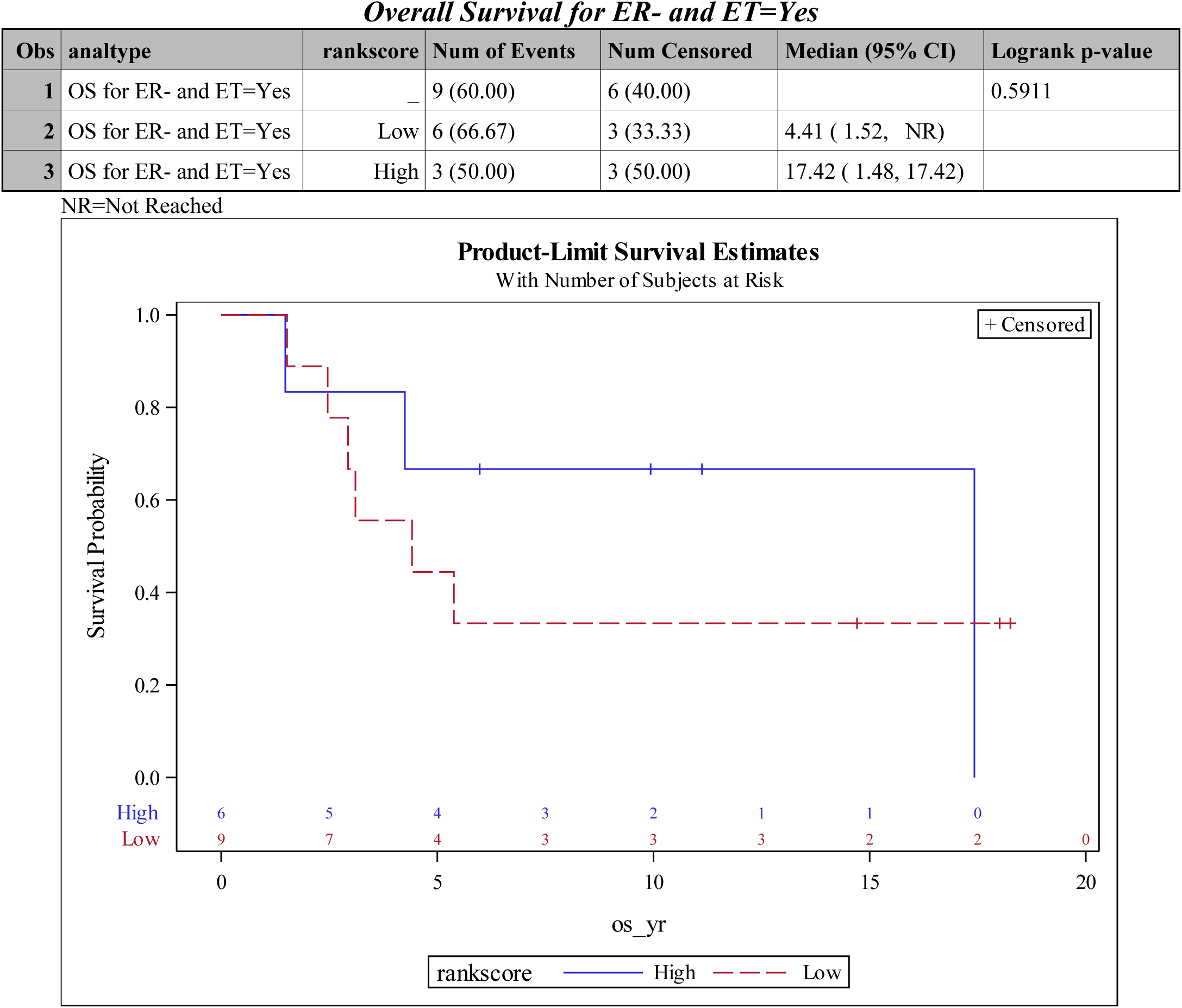
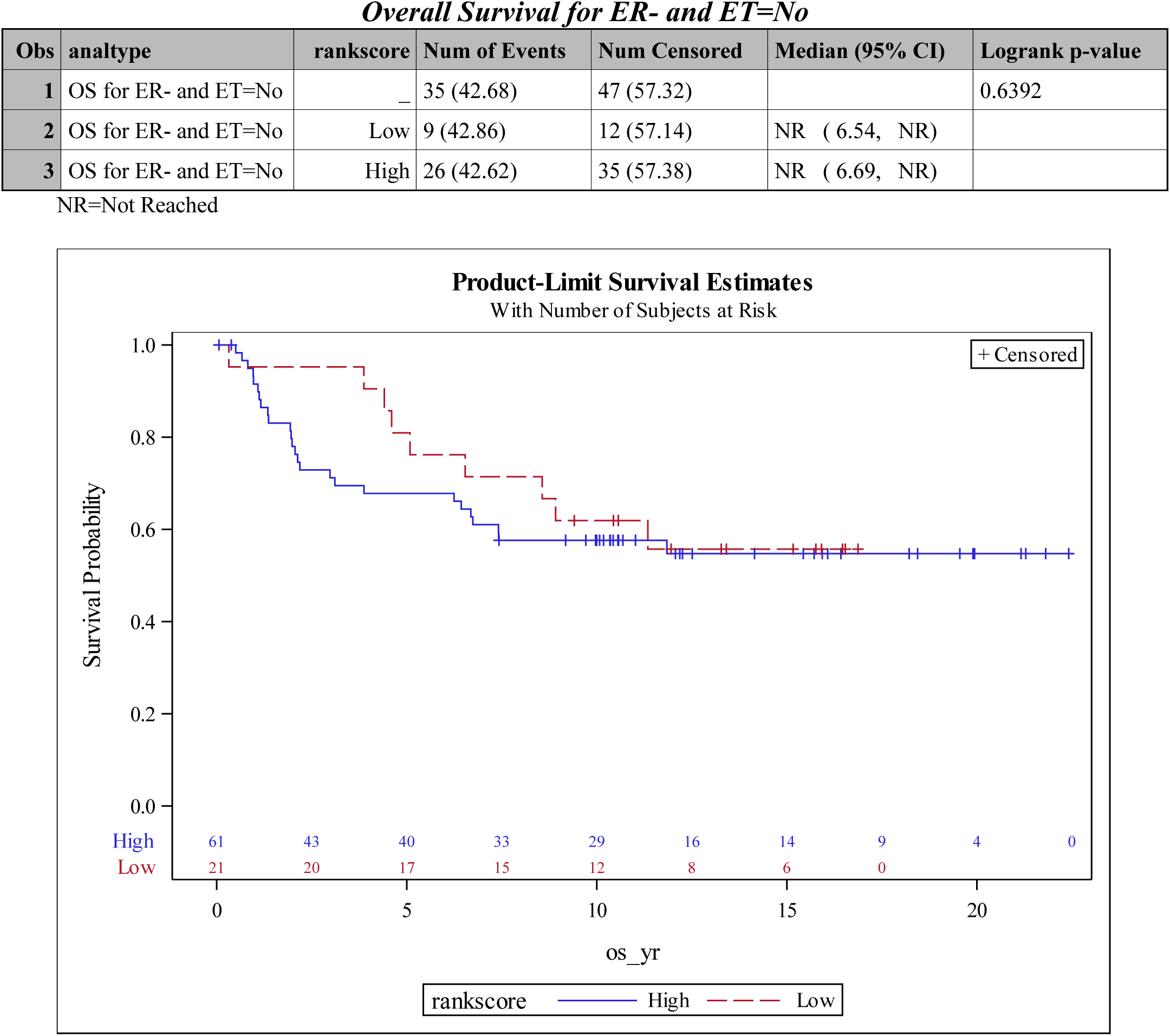
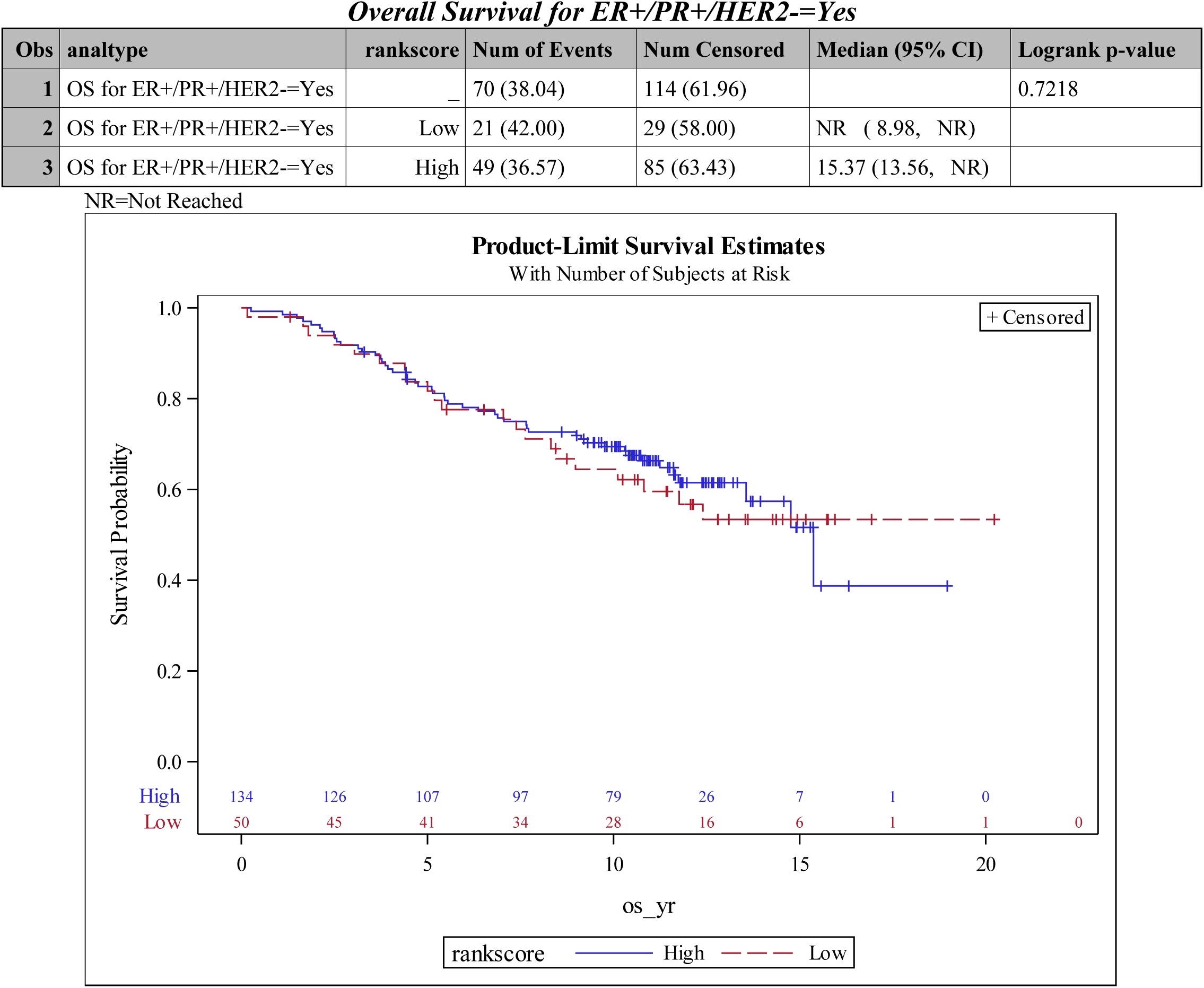
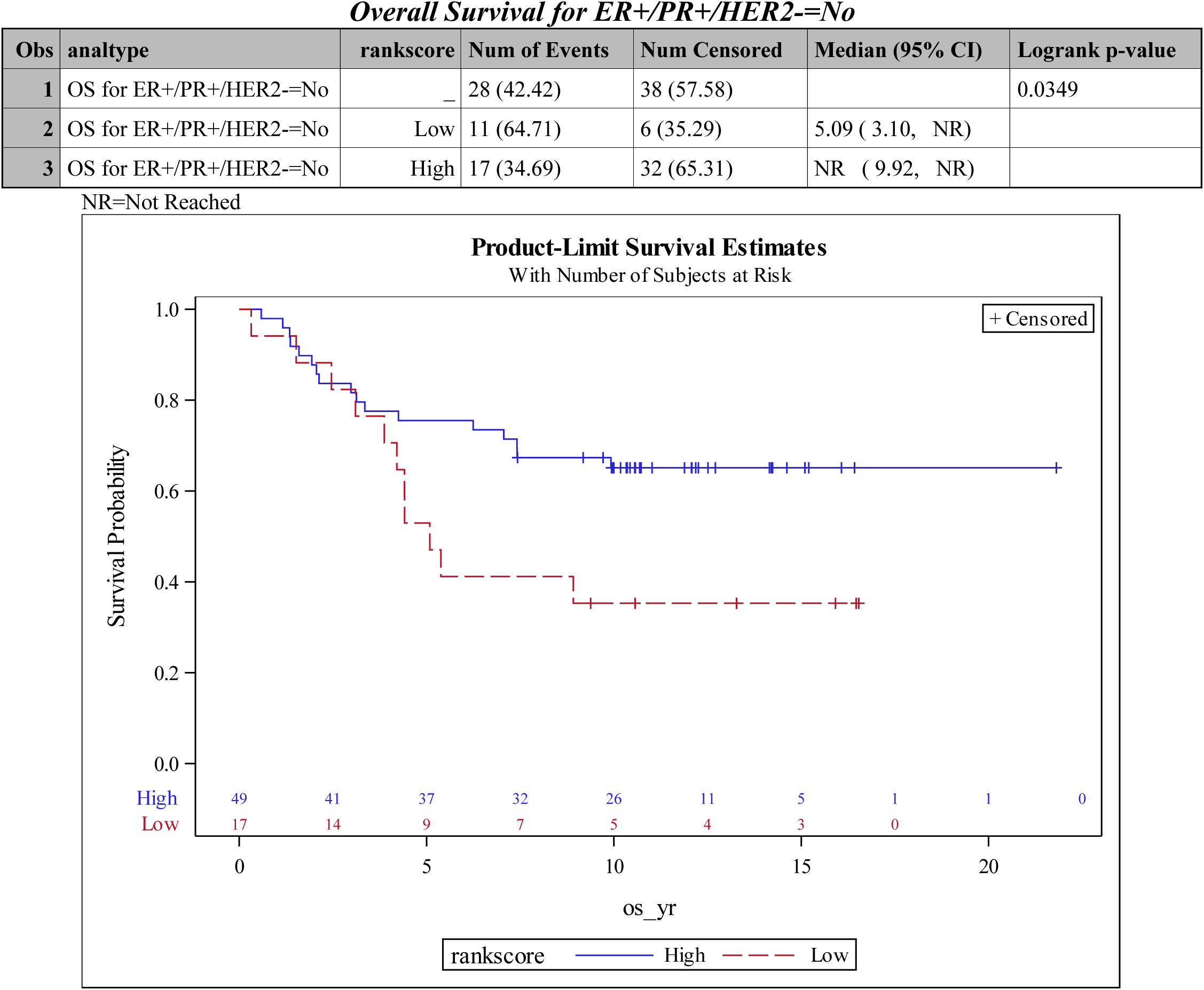
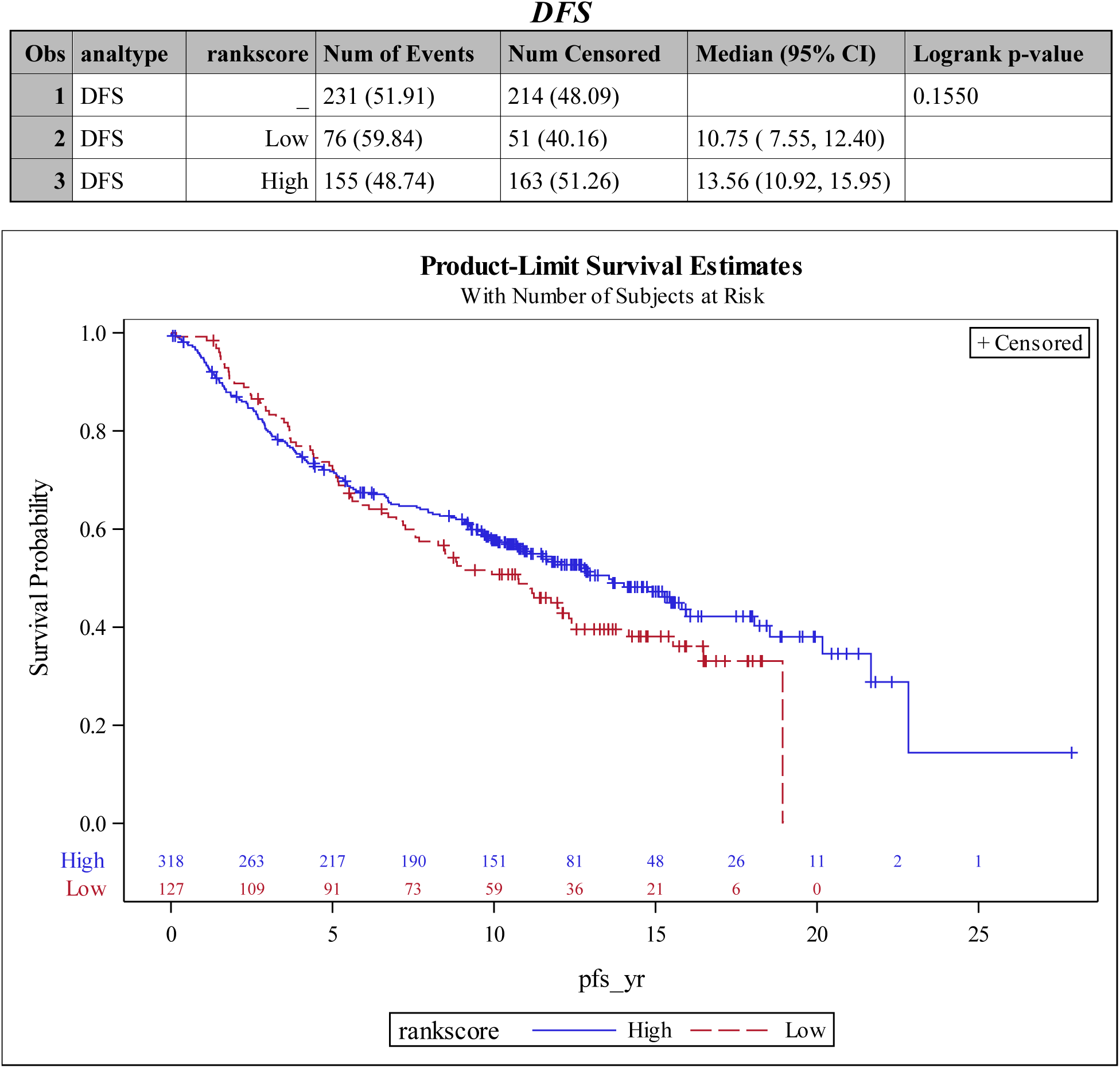
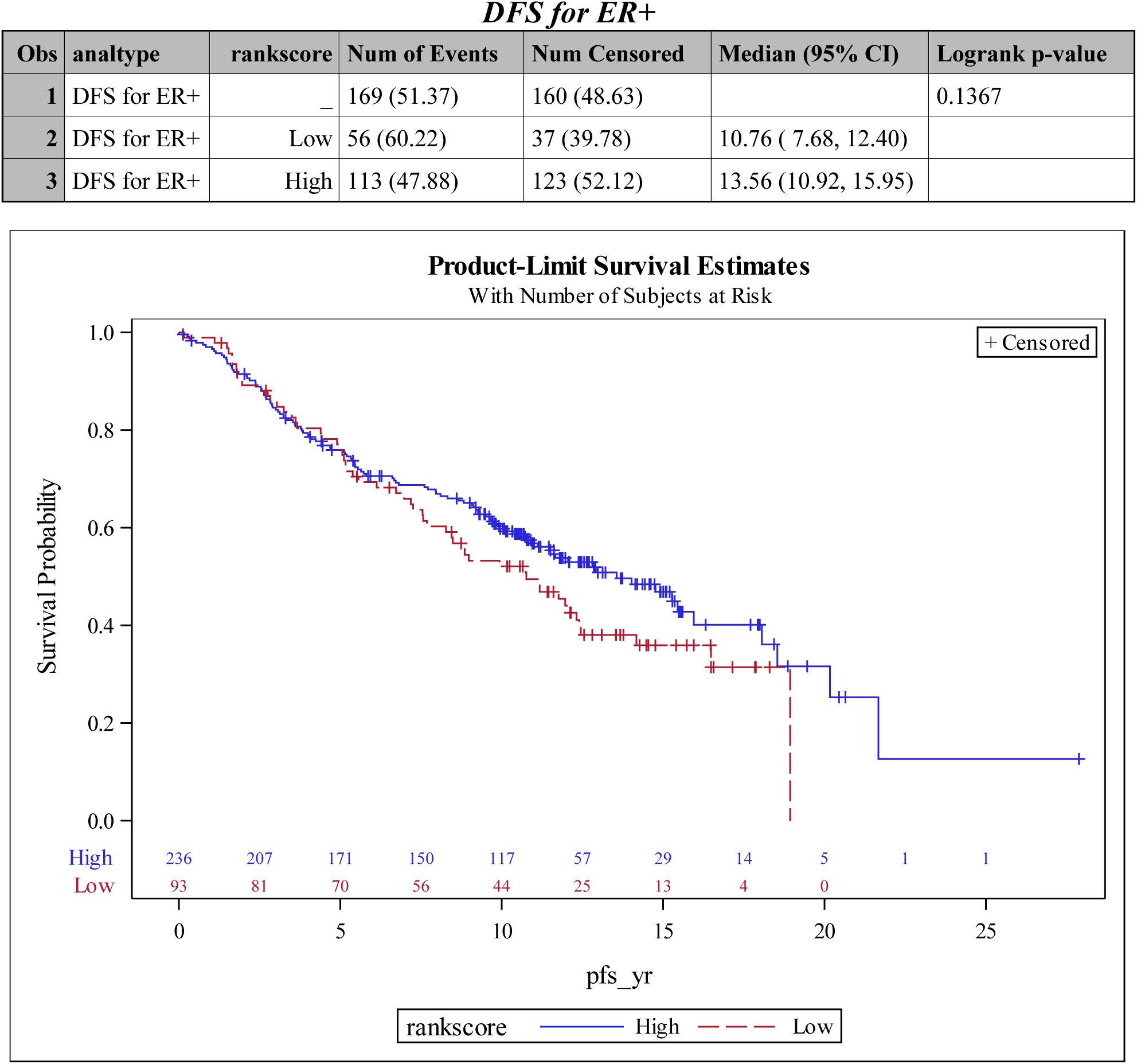
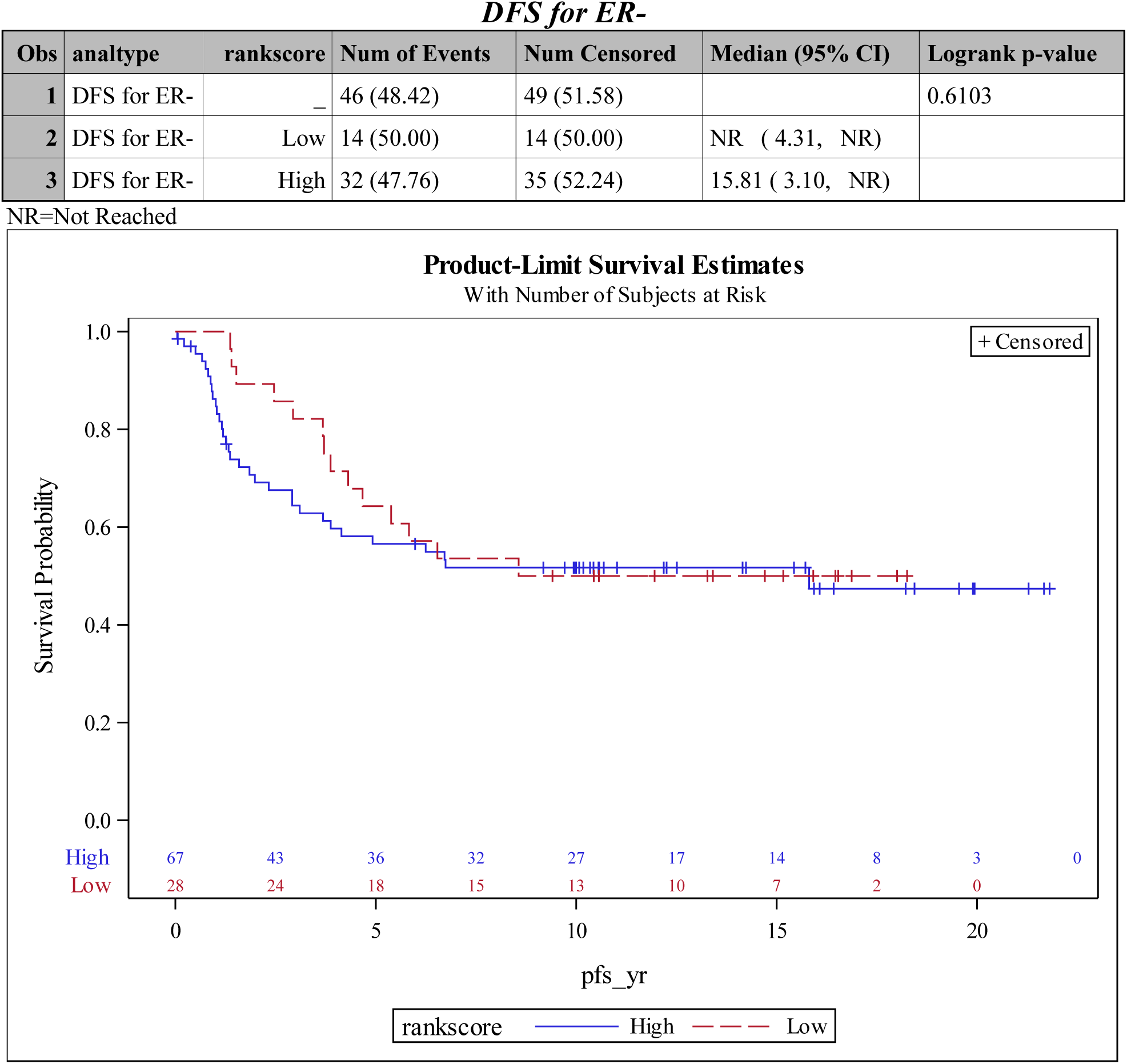
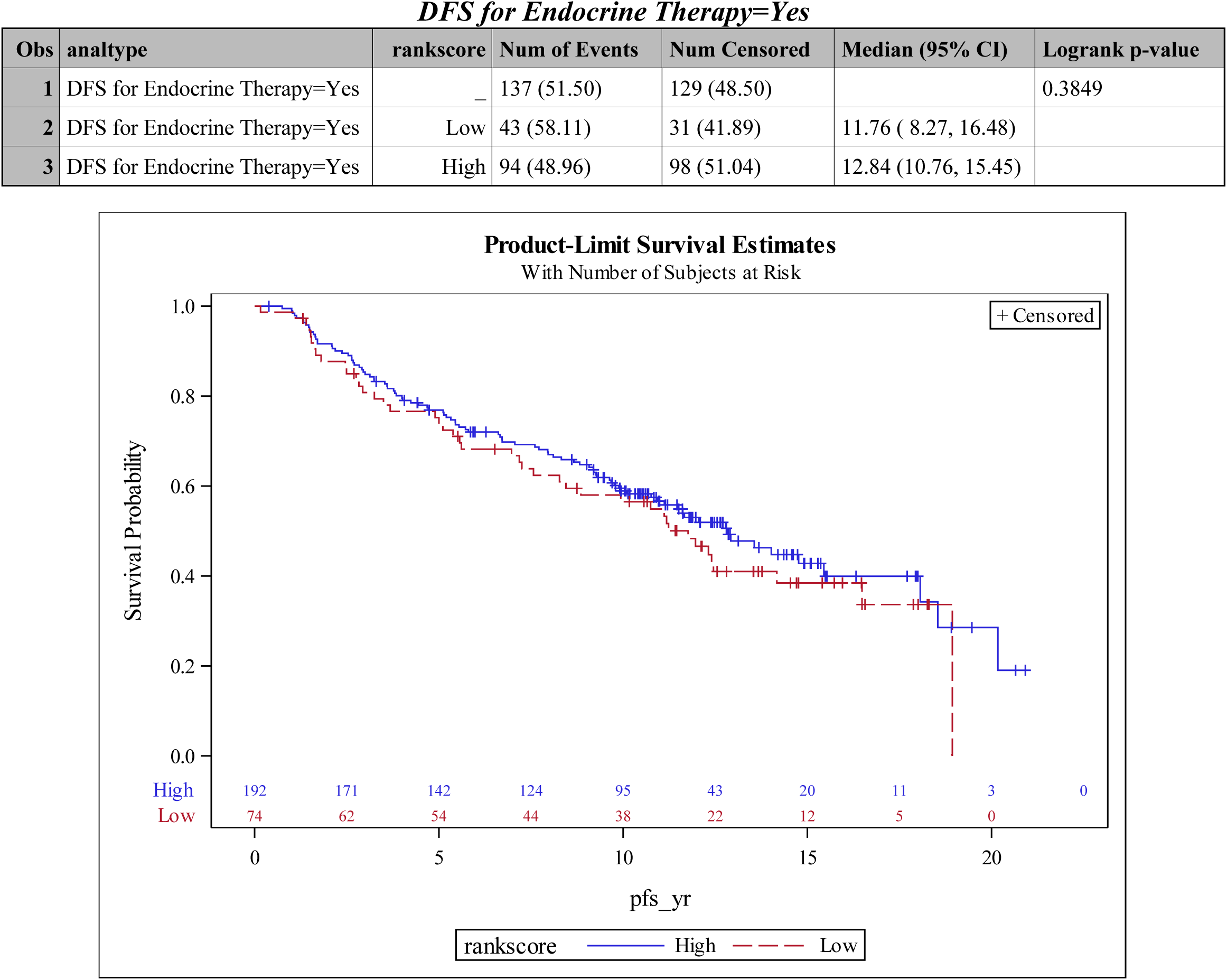
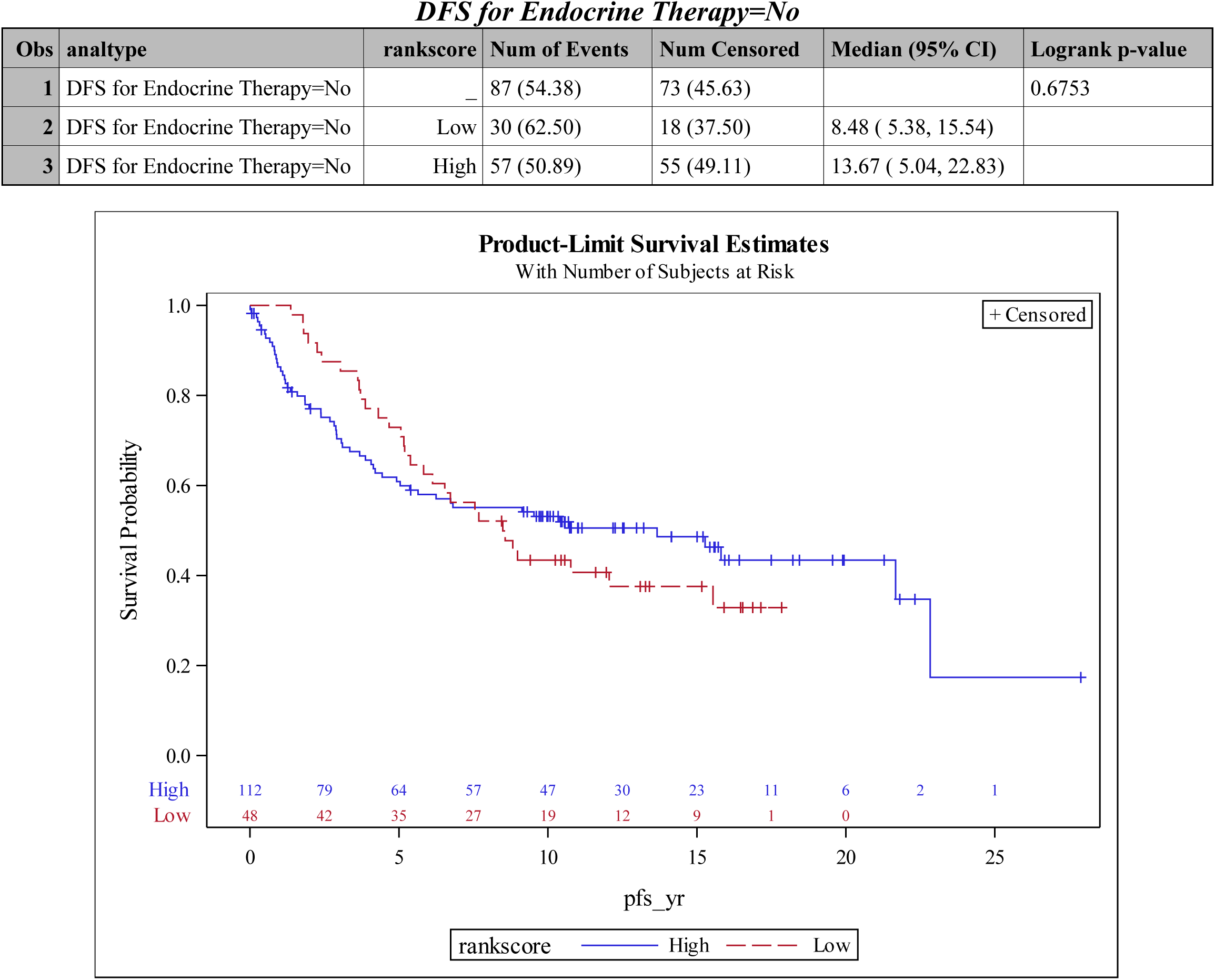
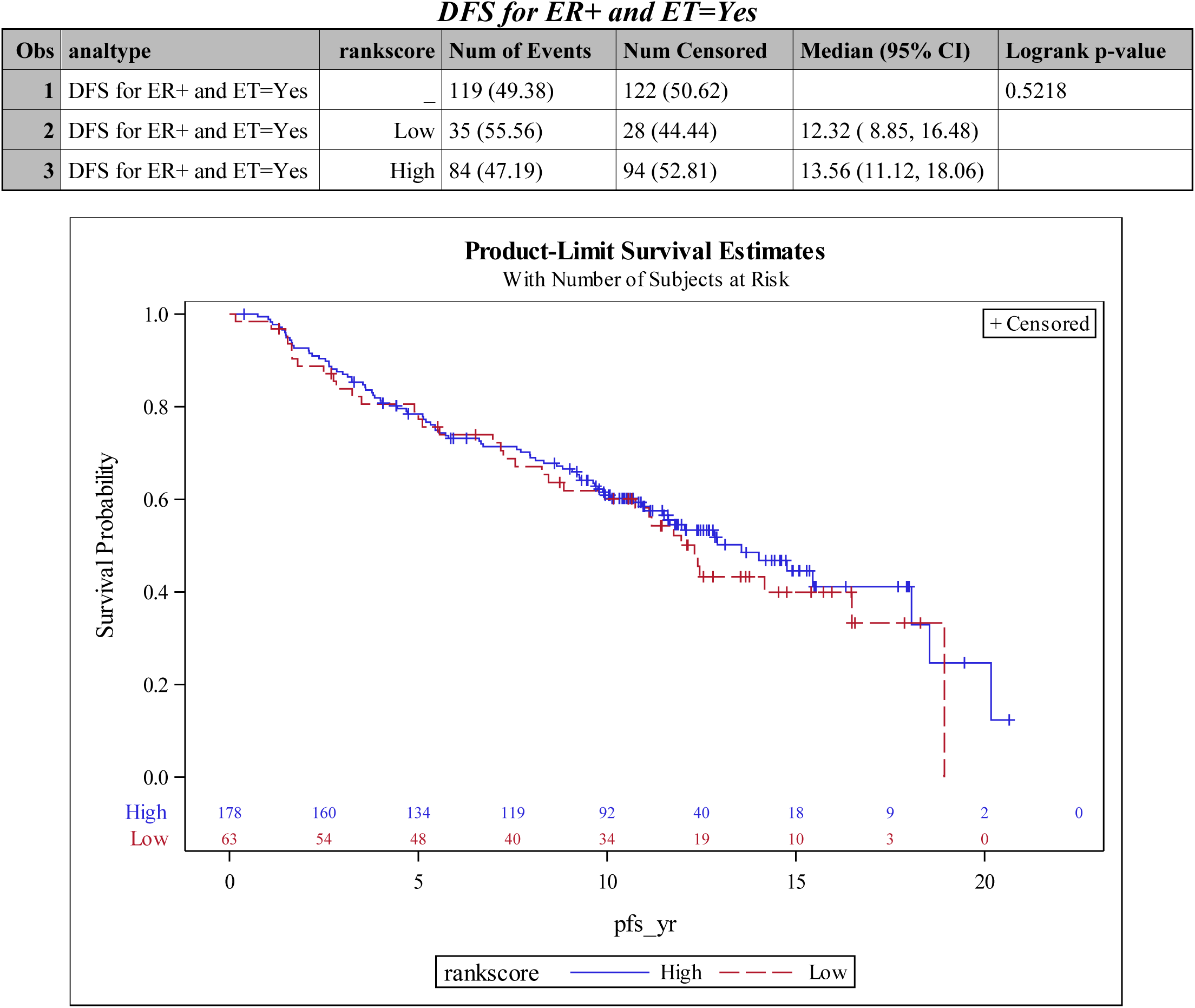
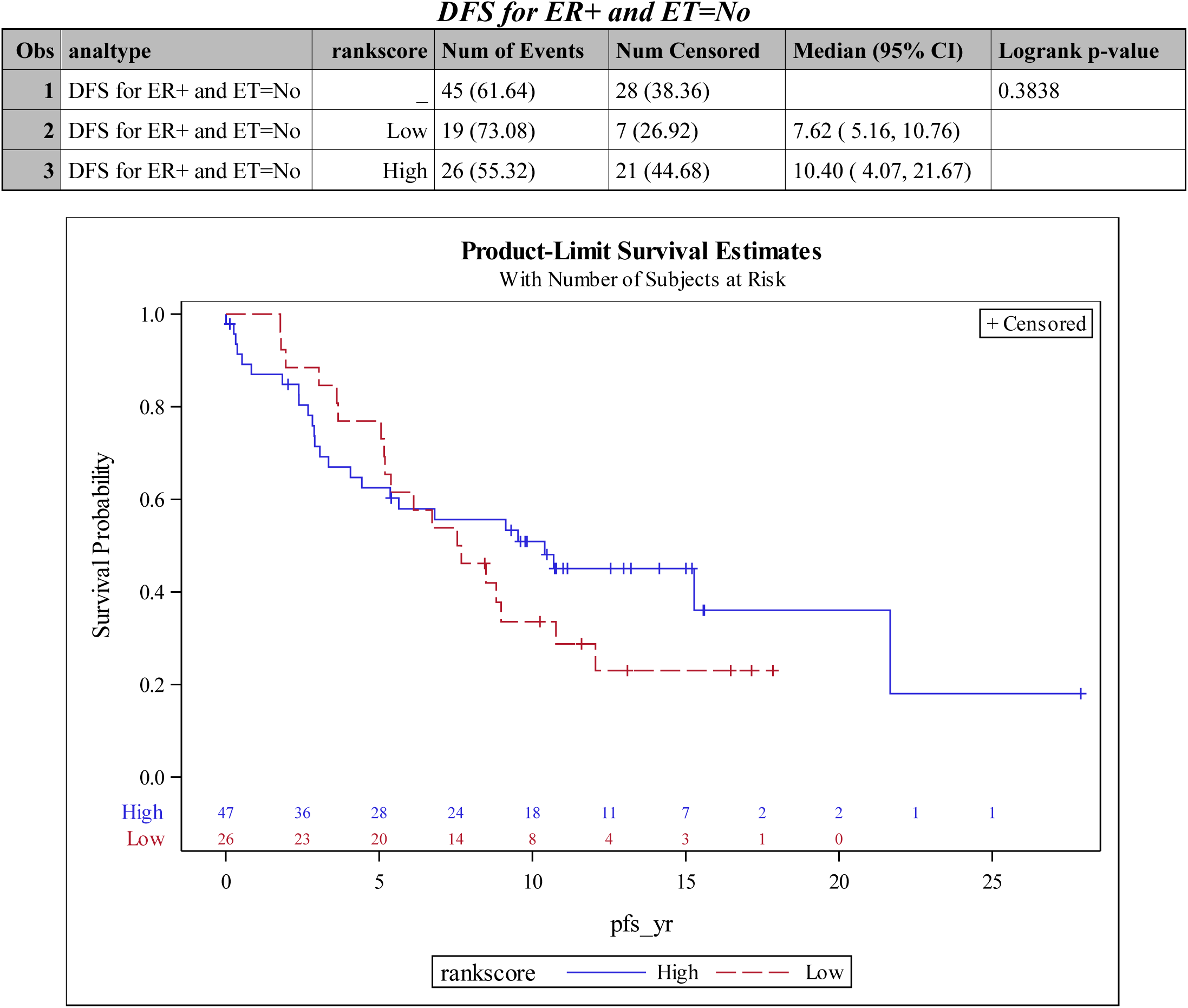
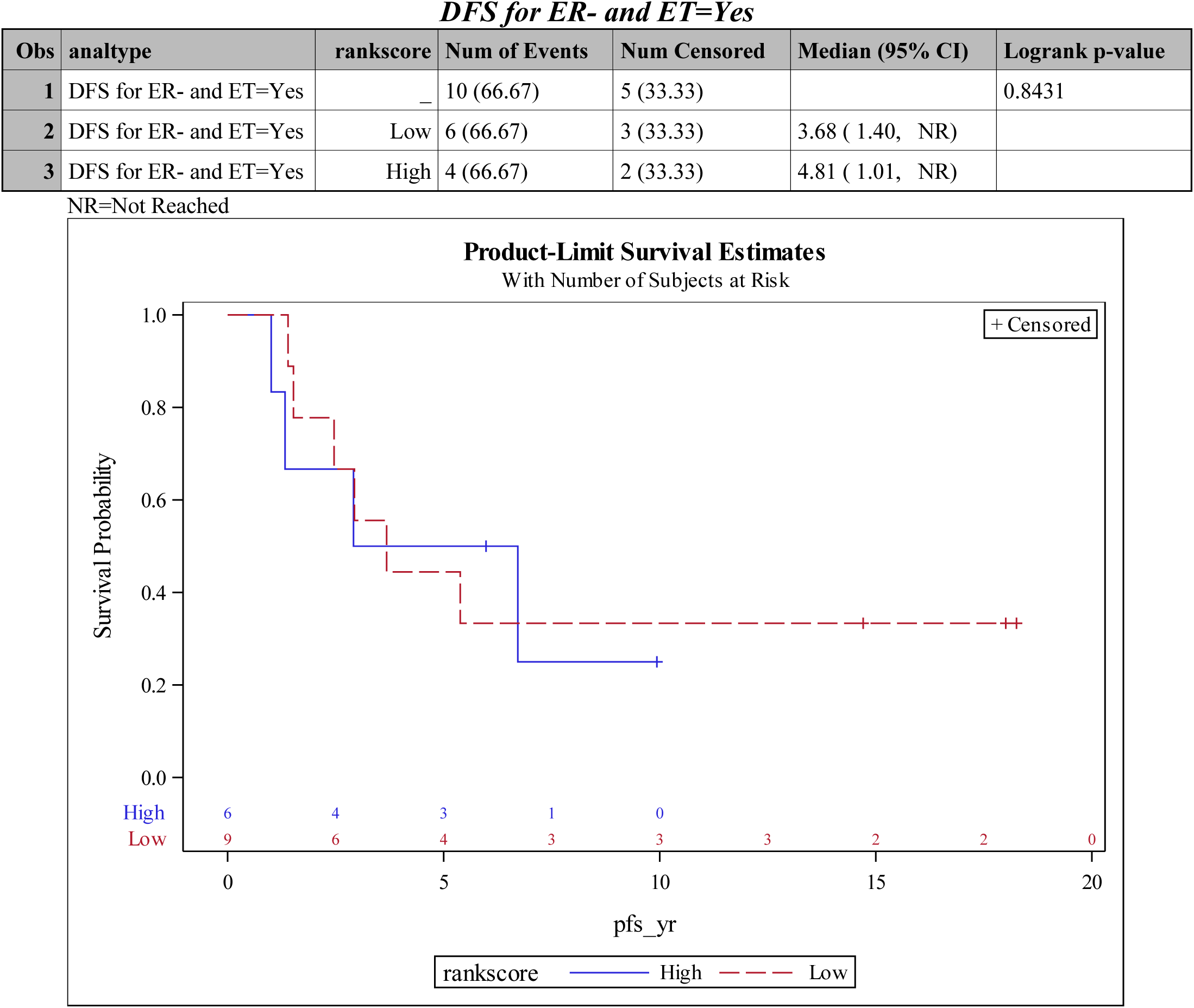
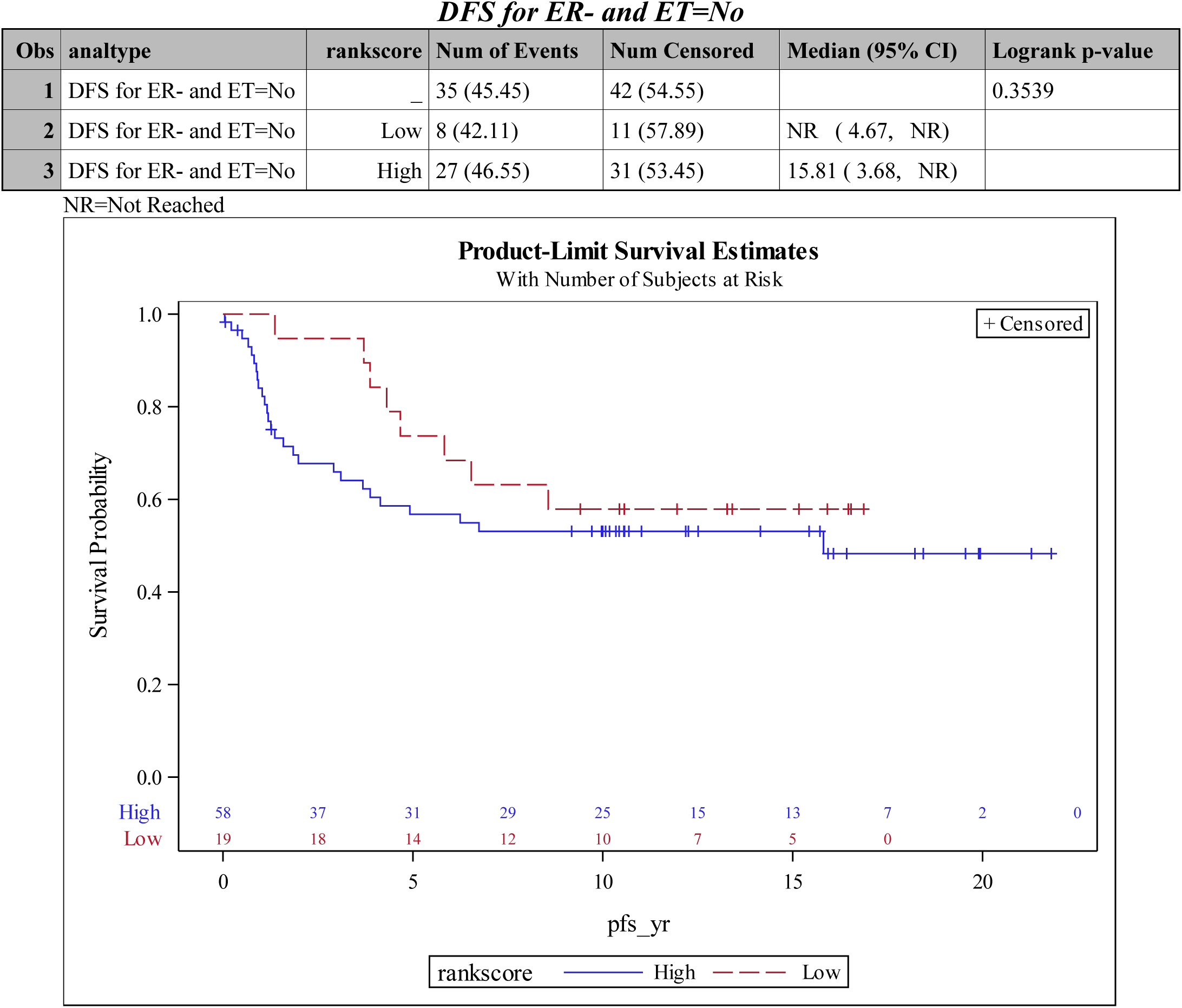
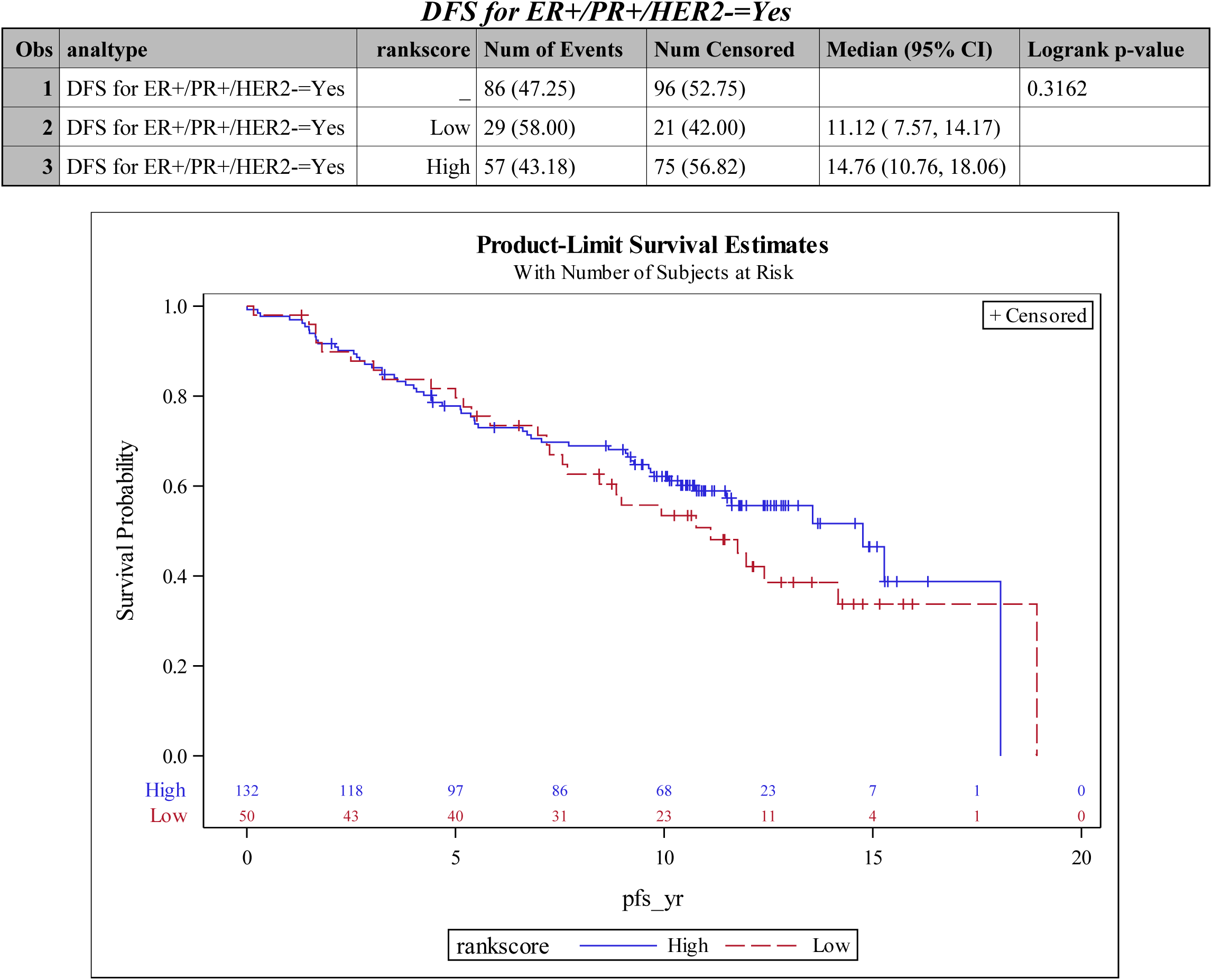
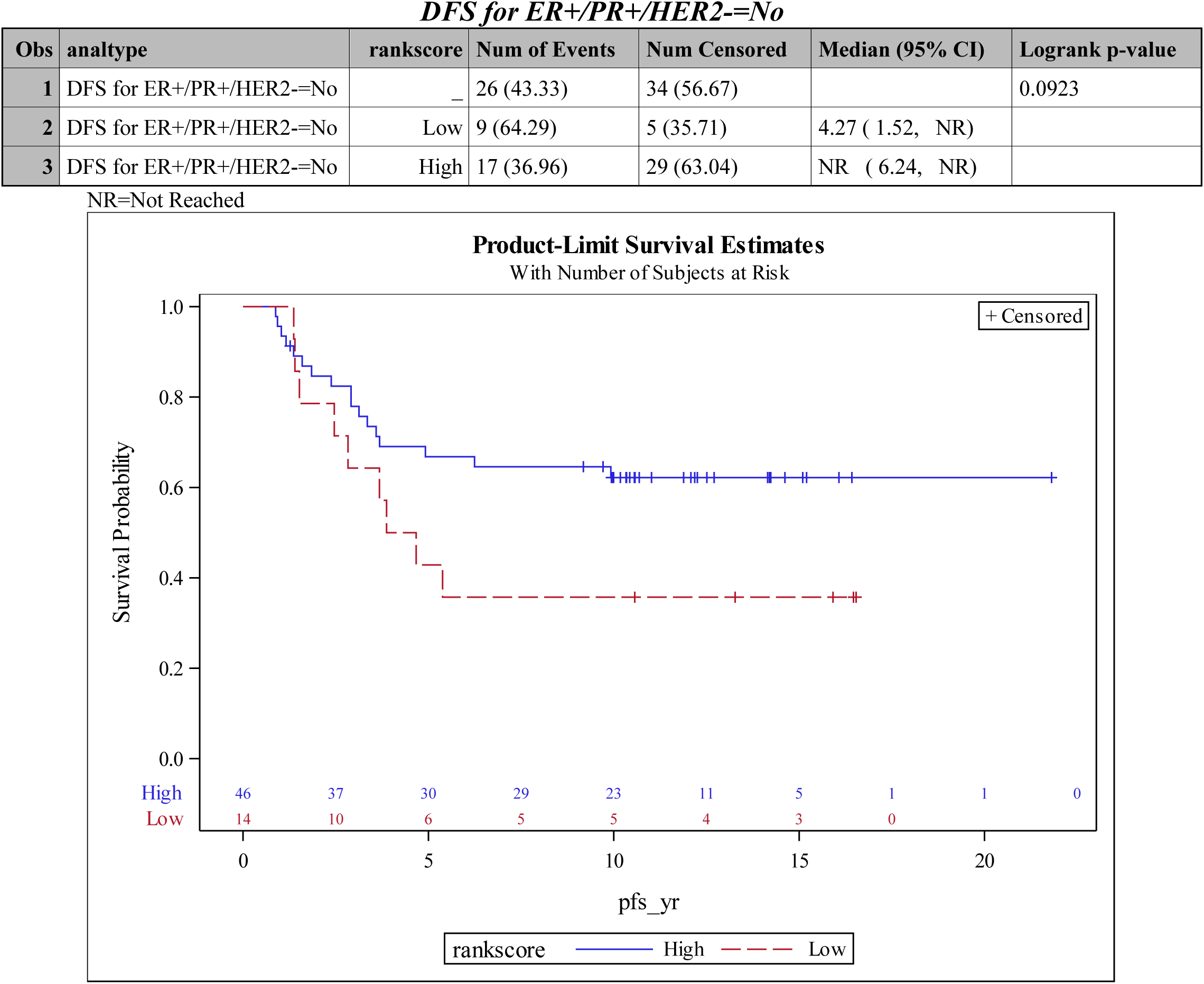
Summary of results for Overall Survival and DFS analysis – Univariate analyses on the TBX3 H-score category.

**Figure S1:**
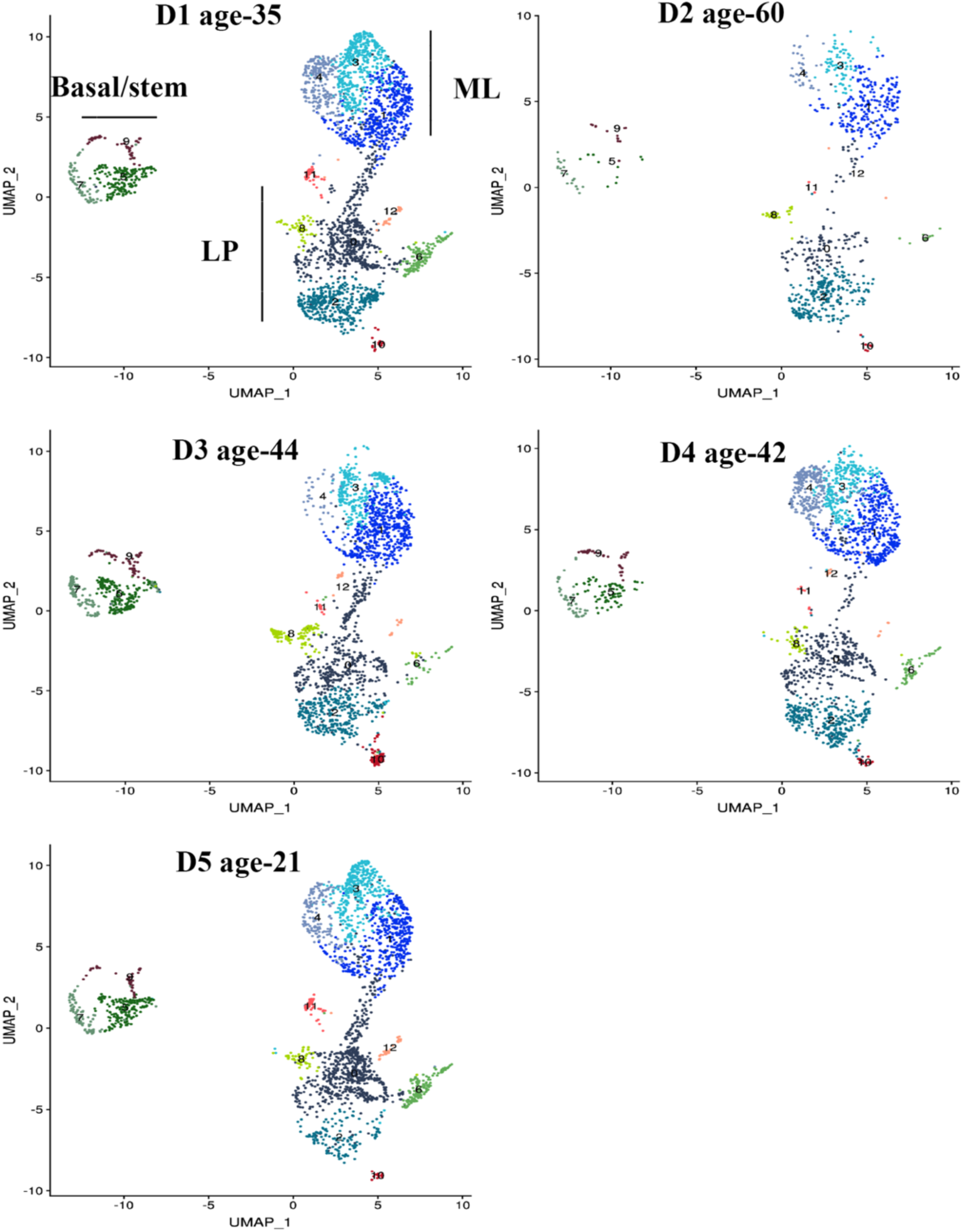
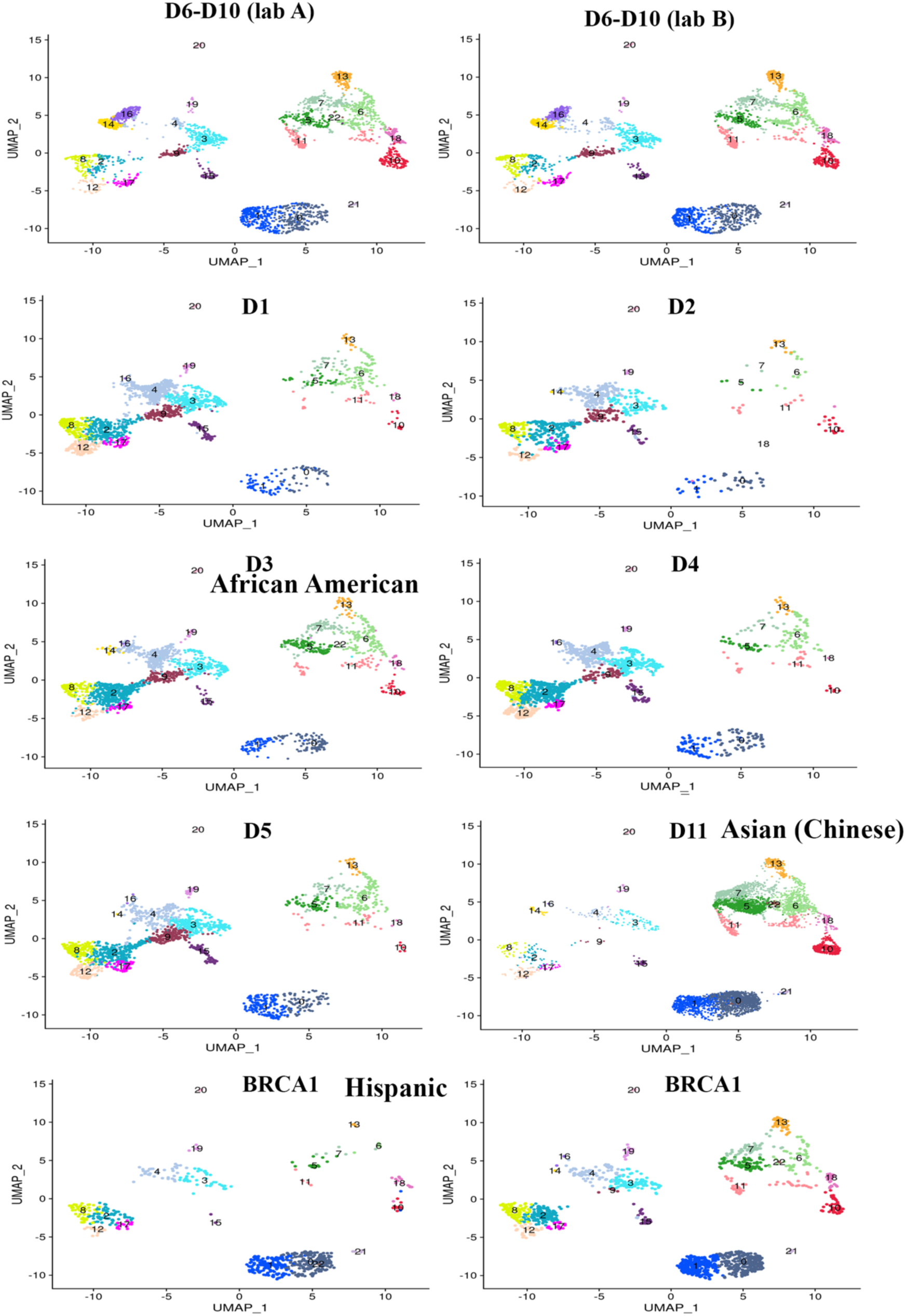
Epithelial clusters in individual samples. A) Epithelial cell clusters in breast tissues of D1 to D5. B) Epithelial cell clusters in first five samples sequenced individually, additional samples sequenced as a pool in two labs (D6-D10), individual sample from an Asian (Chinese), and a BRCA1 mutation carrier. Clustering was done with Seurat and Loupe browser was used to explore various gene expression.

**Figure S2:**
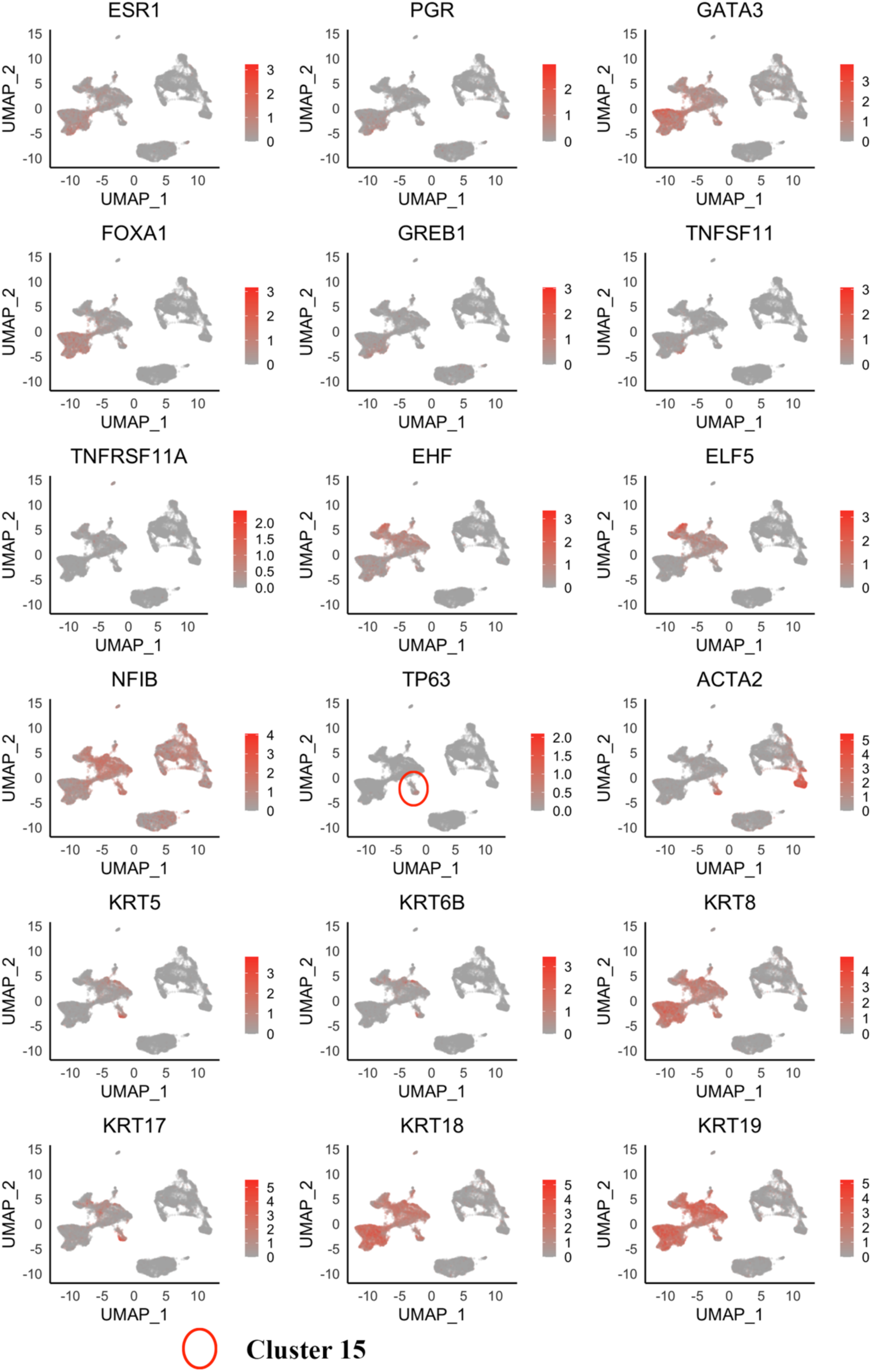
Expression patterns of known mature luminal (ML), luminal progenitor (LP) and basal/stem/myoepithelial cell-enriched marker genes and different keratins in various clusters.

**Figure S3:**
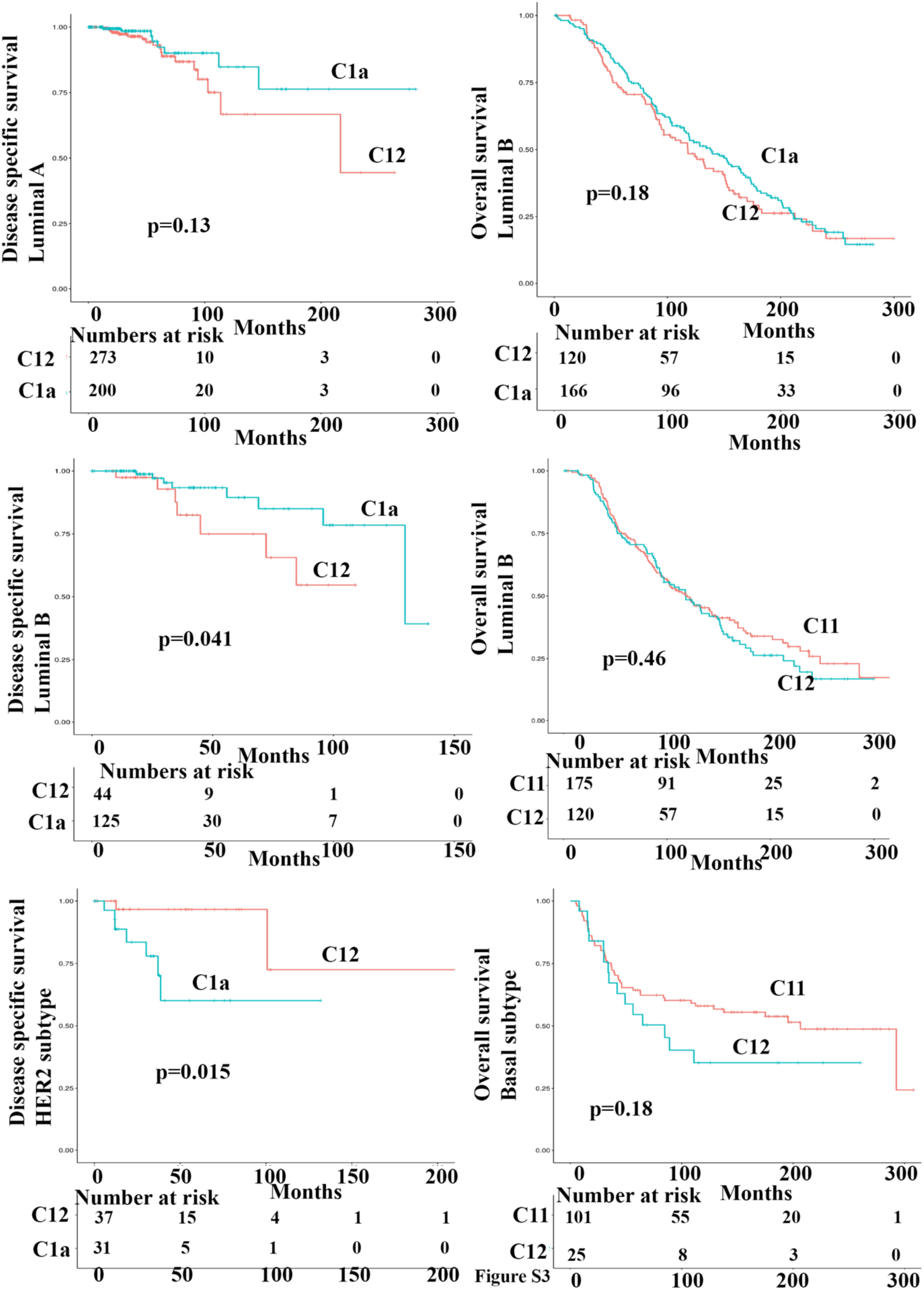
Prognostic value of cluster-enriched genes in various intrinsic subtypes of breast cancer. Data were generated using TCGA and METABRIC datasets.

Table S1: Breast tissue donor information.

Table S2: Sheet 1: Number of cells per cluster of the first five samples. Sheet 2: Number of cells per cluster in the second analysis; Sheet 3: Average expression levels of various genes in clusters of the first analysis. Sheet 4: Signaling network generated using cluster 11 enriched genes.

Table S3: Distribution of various intrinsic subtypes of breast cancers in each cluster in TCGA and METABRIC datasets.

Table S4: Mutation frequency in tumors enriched for specific cluster genes.

