## Supplementary Table 1 for "A single cell atlas of the healthy breast tissues reveal clinically relevant clusters of breast epithelial cells"

**Table S1: Information on breast tissue donors**

| **Donor number** | **Race** | **Age** | **BMI** | **Times pregnant** | **Family history of BC** |
| --- | --- | --- | --- | --- | --- |
| D1 | White | 35 | 24.5 | 1 | NA |
| D2 | White | 60 | 24.9 | 4 | Sisters |
| D3 | African American | 44 | 29 | 4 | Mother, Grandmother and Aunt |
| D4 | White | 42 | 27.8 | 1 | No |
| D5 | White | 21 | 35.4 | 0 | Paternal Grandmother |
| D6 | White | 33 | 23.4 | 2 | Paternal Grandmother |
| D7 | White | 27 | 22.7 | 0 | No |
| D8 | White | 56 | 23.5 | 3 | No |
| D9 | White | 24 | 30.5 | 3 | NA |
| D10 | White | 34 | 26.8 | 2 | No |
| D11 | Asian/Chinese | 43 | 25.6 | 3 | Mother |

**NA=Not available.**
